# Adipose triglyceride lipase is regulated by CAMKK2-AMPK signaling and drives advanced prostate cancer

**DOI:** 10.1101/2022.11.02.514910

**Authors:** Dominik Awad, Thomas L. Pulliam, Meredith Spradlin, Pham Hong-Anh Cao, Elavarasan Subramani, Tristen V. Tellman, Caroline F. Ribeiro, Hubert Pakula, Jeffrey J. Ackroyd, Mollianne M. Murray, Jenny J. Han, Badrajee Piyarathna, Justin M. Drake, Michael M. Ittmann, Cristian Coarfa, Mary C. Farach-Carson, Massimo Loda, Livia S. Eberlin, Daniel E. Frigo

## Abstract

Lipid metabolism plays a central role in prostate cancer. To date, the major focus on prostate cancer lipid metabolism has centered on *de novo* lipogenesis and lipid uptake with little consideration for how cancer cells access these lipids once they are created or taken up and stored. Patient-derived phosphoproteomics identified adipose triglyceride lipase (ATGL), a previously suspected tumor suppressor, as a CAMKK2-AMPK signaling target that, conversely, promotes castration-resistant prostate cancer (CRPC) progression. Phosphorylation of ATGL increased its lipase activity, cancer cell proliferation, migration, and invasion. Shotgun lipidomics and mass spectrometry imaging demonstrated ATGL’s profound regulation of lipid metabolism *in vitro* and *in vivo*, remodeling membrane composition. Inhibition of ATGL induced metabolic plasticity, causing a glycolytic shift that could be exploited therapeutically by co-targeting both metabolic pathways. Together, these data nominate ATGL and intracellular lipolysis as potential therapeutic targets for the treatment of CRPC and provide insights for future combination therapies.

## Introduction

Prostate cancer is the most common non-dermatological cancer in U.S. men, ranking second in cancer-related deaths^1^. The androgen receptor (AR), a transcription factor, remains the central driver of prostate cancer even in the advanced stages of the disease^2^. While AR targeted therapies are initially effective for the treatment of advanced prostate cancers, they are not curative^3^, largely due to the emergence of resistance mechanisms that allow AR signaling to remain active^4^. As such, new therapeutic targets downstream or independent of AR are needed to improve current clinical outcomes.

A direct target of AR is the gene *CAMKK2* (*calcium/calmodulin-dependent protein kinase kinase 2*)^5^, which can promote prostate cancer cell proliferation, survival, and migration in part through the phosphorylation and activation of AMP-activated protein kinase (AMPK), a master regulator of cellular homeostasis^5–9^. AMPK was first described as a tumor suppressor, due to its association with the upstream activator liver kinase B1 (LKB1), a *bona fide* tumor suppressor^10–12^. However, recent reports indicate that AMPK has context-dependent oncogenic roles in cancer^5,7,8,13–20^. Because of the paradoxical tumor suppressor and oncogenic roles of AMPK in cancer, and its importance in physiology, targeting AMPK directly may be suboptimal. Therefore, we sought to identify the downstream targets of CAMKK2-AMPK signaling that could promote prostate cancer progression and test whether these substrates would represent potential novel therapeutic targets.

Using patient-derived phosphoproteomic data, we identified adipose triglyceride lipase (ATGL, encoded by the gene *PNPLA2*), the initial and rate-limiting step in the of the breakdown of triglycerides (TG)^21–23^, as being highly phosphorylated in metastatic castration resistant prostate cancer (mCRPC) and a potential AMPK substrate. ATGL was of particular interest given the importance of lipid metabolism in prostate cancer^24^. To date, the majority of research in prostate cancer lipid metabolism has focused on *de novo* lipogenesis or fatty acid uptake^8,25–32^ with little consideration for how prostate cancer cells access these lipids after they are stored in lipid droplets in part as a mechanism to help avoid lipid toxicity. Notably, ATGL was described as a tumor suppressor in lung cancer^33,34^, while in prostate cancer conflicting reports have been published^35,36^. Although some studies showed a dramatic effect upon targeting ATGL *in vitro* via siRNA or shRNA^36^, additional studies observed only minor effects upon targeting ATGL in cell culture^35^. These prior reports, combined with our correlative data, indicated that a deeper investigation of ATGL’s regulation and role in prostate cancer was warranted.

## Results

### ATGL is highly phosphorylated in human mCRPC at Ser404, a target of AR-CAMKK2-AMPK signaling

To identify potential AMPK targets that could play a role in prostate cancer progression in an unbiased manner, we mined clinical phosphoproteomic data^37^ using an improved AMPK substrate motif^38^ to detect changes in the phosphorylation of predicted AMPK targets across benign, hormone-sensitive prostate cancer (HSPC) and mCRPC (isolated via a rapid autopsy program^37^) disease states. Through the application of an updated position weight matrix (PWM), a mathematical algorithm based on the improved AMPK target phosphorylation motif^38^, we could nominate high-confidence AMPK targets across patient groups. This resulted in the identification of classic AMPK targets, such as the acetyl-CoA carboxylase (ACC)^6^, as well as potential new targets of AMPK in prostate cancer (**Figure 1; Supplementary File 1**).

**Figure 1:**
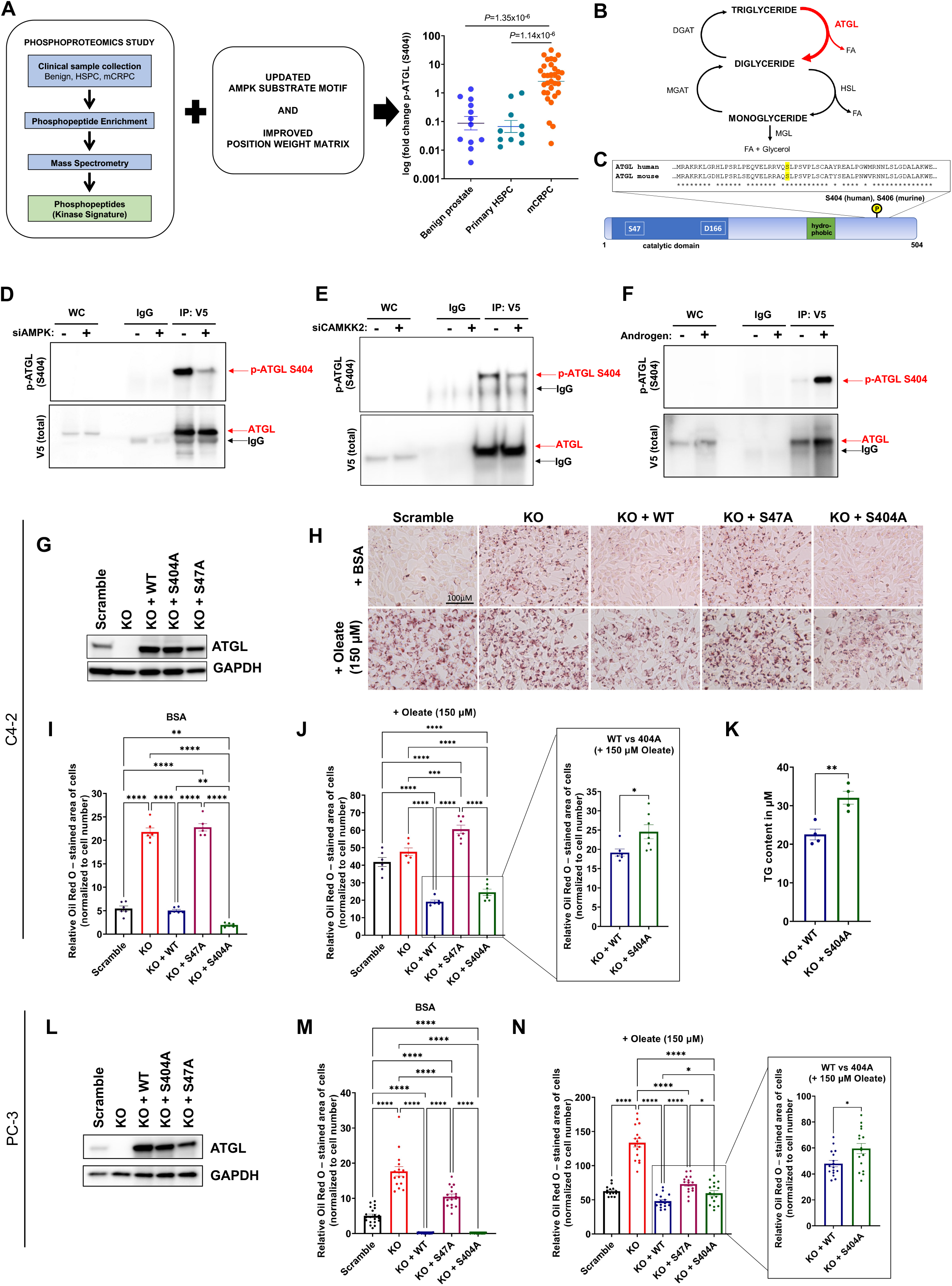
ATGL is highly phosphorylated at S404 in human mCRPC, is targeted by AR-CAMKK2-AMPK signaling, and fine-tunes access to intracellular lipids. (A) ATGL (encoded by *PNPLA2*) is highly phosphorylated at S404 in tumor samples from men with mCRPC (n=31) compared to benign (n=12) or primary, treatment-naïve hormone sensitive prostate cancer (HSPC) (n=10). (B) ATGL is the rate-limiting step in the breakdown of triglycerides (TGs), stored in intracellular lipid droplets. (C) Murine ATGL has high similarity to human ATGL and in particular the murine S406/human S404 site. (D-F) ATGL-V5 was overexpressed in C4-2 cells and then treated with siRNAs targeting scramble control, the AMPK a catalytic subunits (*PRKAA1/2*) or *CAMKK2* (siAMPK and siCAMKK2) or synthetic androgen (R1881) for 72h (validation of siRNA efficacy and androgen-mediated effects on CAMKK2-AMPK signaling are shown in Supplemental Fig. S1). Immunoprecipitated ATGL was tested for phosphorylation status using a developed p-ATGL S404 antibody (validation in Supplemental Fig. S1). (G-N) Oil Red O staining demonstrated that knockout of ATGL (*PNPLA2*) increased neutral lipids in AR+ C4-2 (G-K) and AR-PC-3 (L-N) cells. (I and M) This phenotype can be rescued in the presence of BSA control when ATGL wildtype (WT) or a S404A mutant is re-expressed but not the lipolytic-inactive S47A mutant. 150 μM oleate (complexed with BSA) led to an increase of detected triglycerides in S404A mutant cells compared to WT as assessed by (J and N) Oil Red O staining or (K) directly measuring TG content. Images and graphs are representative results of at least three independent experiments. One-way ANOVA or unpaired two-tailed t-test. \**P* < 0.05, \*\**P* < 0.01, \*\*\**P* < 0.001, \*\*\*\**P* < 0.0001, ns = no significance.

The substrate that exhibited the greatest change was ATGL (encoded by the gene *PNPLA2*), which was highly phosphorylated in mCRPC compared to treatment-naïve, localized prostate cancer and/or benign prostate tissue (**Figure 1A**). The levels of ATGL phosphorylated at S404 exhibited a ~40-fold increase in mCRPC patient samples compared to primary HSPC and/or benign prostate tissue (**Figure 1A**), higher than any other candidate AMPK substrate (**Supplementary File 1**). ATGL’s known role in the breakdown of TGs has been reported in other tissues^39^ (**Figure 1B**). Prior work in murine adipose tissue demonstrated that the phosphorylation at S406, thought to be the murine ortholog of human S404 (**Figure 1C**), stimulated ATGL’s lipolytic activity^40,41^. While human ATGL demonstrates high sequence similarity with murine ATGL (**Figure 1C**), the phosphorylation of human ATGL by AMPK has not been studied in detail. Previous studies have expressed murine ATGL constructs in human cell lines to demonstrate its regulation by AMPK^41^ or have reported on human ATGL’s phosphorylation at S404 by AMPK using a murine-specific p-ATGL S406 antibody^42^. When we tested the specificity of a popular murine-specific p-S406 antibody to detect changes in human p-ATGL (S404) levels, this murine-target antibody could not detect human p-ATGL (**Supplemental Figure S1A**). Thus, to determine if human ATGL was targeted by AMPK in human prostate cancer cells, we created and validated our own highly specific p-ATGL S404 antibody (**Supplemental Figure S1B**). While highly specific, this antibody did not detect endogenous p-ATGL. Hence, we expressed V5-tagged ATGL constructs to test the regulation of ATGL at S404 by the AR-CAMKK2-AMPK signaling axis in human CRPC C4-2 cells. We immunoprecipitated C-terminal V5-tagged human ATGL, since the C-terminal tagging of ATGL does not interfere with its activity^39^. Phosphorylation of ATGL at S404 was decreased following RNAi-mediated knockdown of the catalytic subunits of AMPK (*PRKAA1/2*) (**Figure 1D; Supplemental Figure S1C**) or *CAMKK2* (**Figure 1E; Supplemental Figure S1D**). We further confirmed that AMPK mediated the phosphorylation of ATGL(S404) using a second antibody that broadly recognizes proteins and peptides bearing the canonical LXRXXS AMPK substrate motif (**Supplemental Figure S1E**). Conversely, treatment of C4-2 cells with the synthetic androgen R1881 increased p-ATGL (S404) levels (**Figure 1F; Supplemental Figure S1F**), consistent with CAMKK2-AMPK signaling being a known downstream effector of AR in prostate cancer^5,7,8,20,43^. Taken together, these data indicate that ATGL is highly phosphorylated at S404 in mCRPC in patients and that this site is targeted by AR-CAMKK2-AMPK signaling.

### ATGL controls access to intracellular lipids in multiple prostate cancer cell models

We next evaluated ATGL’s role in prostate cancer lipid metabolism. We first tested the effect of ATGL knockdown on neutral lipid levels (assessed by Oil Red O staining) using chemical siRNAs targeting *PNPLA2* (encoding ATGL) in a variety of prostate cancer cell models (LNCaP (AR+hormone-sensitive), C4-2 (AR+ CRPC derivative of LNCaP), C4-2B (more aggressive variant of AR+ C4-2), 22Rv1 (AR+ CRPC), PC-3 (AR-CRPC)), and non-transformed prostate epithelial cells (RWPE1) (**Supplemental Figures S2A-B**). We observed an increase in neutral lipid accumulation following ATGL knockdown in multiple prostate cancer cell models, consistent with a block in the breakdown of TGs. However, ATGL knockdown did not change neutral lipid levels in non-transformed RWPE-1 prostate epithelial cells. As expected, hormone-sensitive LNCaP cells also demonstrated an increase in neutral lipids upon androgen treatment, consistent with AR’s known role in *de novo* lipogenesis, which was further increased following ATGL knockdown (**Supplemental Figures S2A-B**). To further investigate ATGL’s role in prostate cancer cell lipid metabolism, we created a series of isogenic CRPC *PNPLA2* (ATGL) knockout (KO) and addback cell lines, including a S404A mutant as well as a lipolytic-inactive S47A mutant^39^ (**Figures 1G-N; Supplemental Figure S2C and S3**). ATGL KO consistently increased intracellular neutral lipid levels, an effect that was reversed by the re-expression of ATGL wildtype (WT) but not lipolytic-inactive S47A mutant in three different CRPC cell lines, including the AR-CPRC cell model PC-3 (**Figures 1G-N, Supplemental Figure S3**). There was no difference in neutral lipid levels when comparing the WT and S404A addbacks in cells treated with bovine serum albumin (BSA) vehicle; however, there was a modest but consistent increase in neutral lipids in the S404A mutant-expressing cells compared to ATGL WT after cells were treated overnight with exogenous fatty acids (150 μM oleate) in all three cell models (**Figures 1J,N; Supplemental Figure S3D,H, L**). Since Oil Red O is an indirect measurement of triglycerides, broadly staining all neutral lipids, we also directly measured the triglyceride content of cells using a Triglyceride-Glo™assay and observed higher triglycerides levels in ATGL S404A-expressing prostate cancer cells compared to WT addback control (**Figure 1K**). As noted previously^39^, high expression of ATGL WT or even ATGL 404A was sufficient to decrease lipid levels below basal levels (**Supplemental Figure S4**). Together, these data suggest that ATGL is a major regulator of TG catabolism in prostate cancer cells and that phosphorylation of human ATGL at S404 can fine-tune ATGL’s lipolytic activity when cells are challenged with higher lipid levels. However, the posttranslational regulation of ATGL’s lipolytic activity can be overshadowed by high expression of ATGL.

### ATGL mediates prostate cancer growth *in vitro* and *in vivo*

Prior reports have demonstrated that ABHD5 (also called CGI-158), a co-activator of ATGL in murine adipose tissue, can function as a tumor suppressor in prostate cancer^36^, suggesting that perhaps ATGL could also function as a tumor suppressor. However, our phosphoproteomic analyses indicate that ATGL activity is increased in mCRPC (**Figure 1**). Moreover, humans and mice with *ABHD5* and *PNPLA2* (encoding ATGL) genetic alterations present different phenotypes suggesting that ABHD5 and ATGL may possess divergent functions^44,45^. Notably, genetic and transcriptomic data demonstrate that *PNPLA2* levels correlate with poor prognosis in a large cohort of men with mCRPC (**Figure 2A**; *n* = 444; divided into two RNA-Seq experimental methods; probe-capture and poly(A)+ selection), indicating that *PNPLA2* has the genomic and transcriptomic traits of an oncogene. Functionally, KO of *PNPLA2* had minimal effects on CRPC cell growth when cells were grown in the presence of standard cell culture media conditions (11.1 mM glucose) (**Figure 2B**). However, when glucose concentrations were lowered to mimic physiological levels found in the tumor microenvironment, ATGL KO impaired cell growth (**Figure 2B**). When we tested additional isogenic CRPC cell lines in colony formation assays, which introduces density-dependent metabolic stress^46^, we observed that ATGL KO decreased colony formation in all three cell models tested (C4-2, C4-2B-LT, PC-3) (**Figure 2C**). The impaired colony formation was rescued by ATGL WT addback but not S404A or S47A mutants (**Figure 2C**). This phenotype was observed under physiological glucose concentrations (5 mM) and was enhanced when glucose concentrations were lowered further (**Supplemental Figures S5**). Similar effects were observed in AR-PC-3 cells (**Figure 2C; Supplemental Figure S5C**), suggesting that CAMKK2-AMPK signaling may be intact and promoting prostate cancer cell growth independent of AR status. These data are consistent with a prior report demonstrating a pro-cancer role for CAMKK2 in AR-DU-145 cells^47^. To test the roles of CAMKK2 and AMPK in models of AR-prostate cancer cell growth, we used molecular knockdown (siRNA) or pharmacological inhibition (STO-609 and SGC-CAMKK2-1) and found that inhibition of CAMKK2-AMPK signaling suppressed the growth of AR-PC-3 cells as well as the established neuroendocrine prostate cancer (NEPC) cell models MDA-PCa-144-13 and NCI-H660 (**Supplemental Figure S6**).

**Figure 2:**
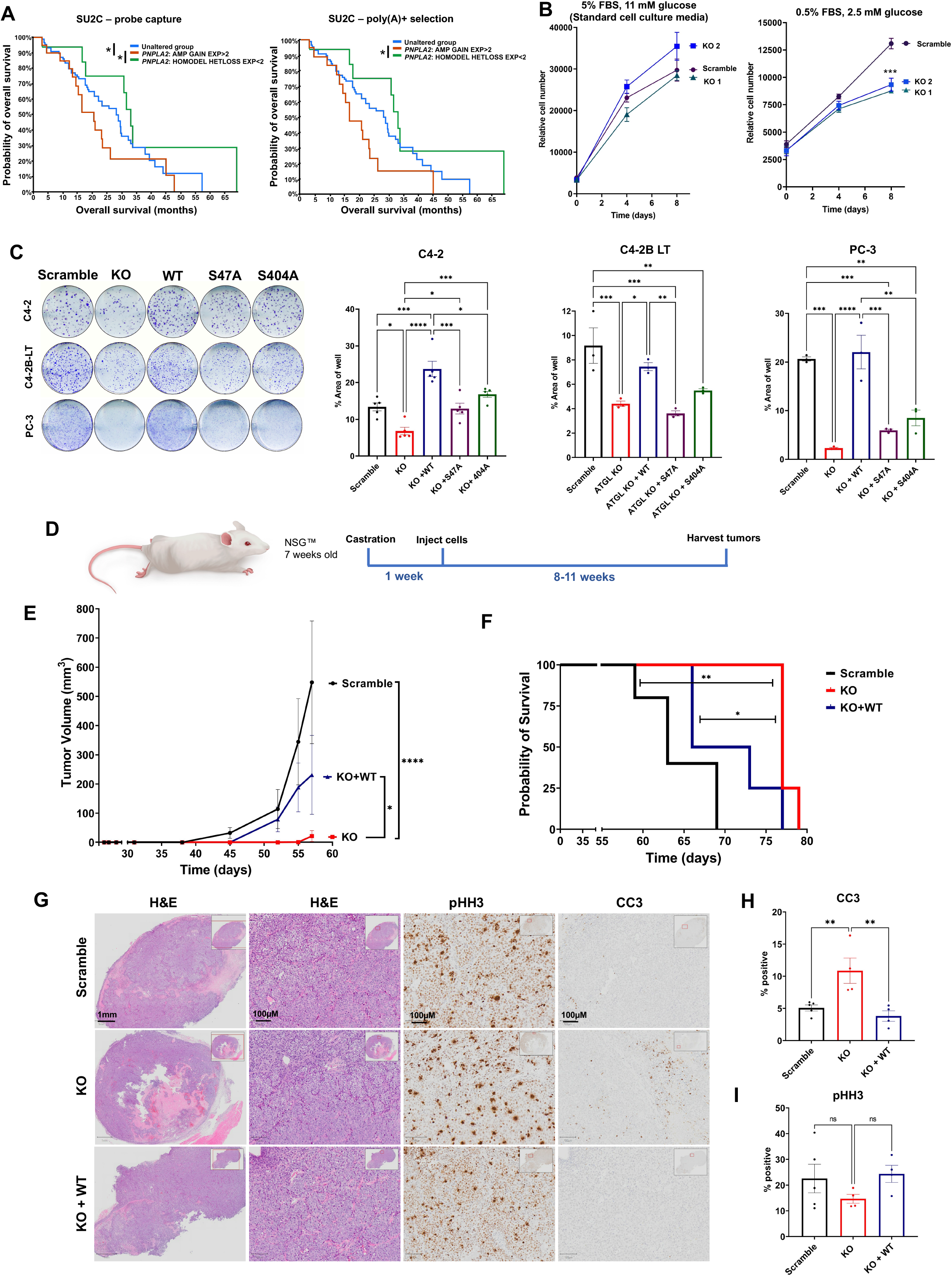
ATGL mediates prostate cancer growth *in vitro* and *in vivo*. (A-B) SU2C patient data demonstrates that the increased expression of *PNPLA2*, genetic amplification, or copy number gain correlates with decreased overall survival compared to unaltered or decreased expression, homozygous deletion, or copy number loss of *PNPLA2 (<2*). *n* = 444; results are shown using tumor samples that were subjected to probe capture or poly(A)+ selection RNA analysis. Logrank test. \**P* < 0.05. (B) Knockout of *PNPLA2* (ATGL KO) in C4-2 did not affect proliferation unless nutrient conditions were adjusted to more physiological conditions (2.5 mM glucose, 0.5% FBS). (C) Colony formation of AR+ (C4-2, C4-2B-LT) and AR- (PC-3) CRPC models. Scramble, ATGL KO and addbacks of ATGL wildtype (WT) or mutants were grown in the presence of 2.5 mM glucose and quantified for % area of well occupied by formed colonies at endpoint. See Supplemental Figure S5 for additional nutrient conditions. Images (*left*) and quantification (*right*) are representative results of at least three independent experiments. (D) Mice were castrated and subcutaneously injected with scramble control (n=5), ATGL KO (n=4), or ATGL KO + WT C4-2 addback cell line derivates (n=4). (E) Average tumor volume until day 57 of experiment (when first control mouse (from scramble group) had to be sacrificed due to tumor size) and (F) survival curve. (G-I) Representative images of tumor tissue. (G) Hematoxylin and eosin stain (H&E), (H) cleaved caspase 3 (CC3), (I) p-Histone H3 (pHH3). Kaplan-Meier curves analyzed by Logrank tests. Other data analyzed using one-way ANOVAs. \**P* < 0.05, \*\**P* < 0.01, \*\*\**P* < 0.001, \*\*\*\**P* < 0.0001, ns = no significance. Data are represented as mean ± SEM (n=3 or as indicated in figure).

To test ATGL’s role in tumor growth *in vivo*, we used a human xenograft mouse model of CRPC where we injected C4-2 scramble control, C4-2 ATGL KO and C4-2 ATGL KO + WT addback cells into castrated 7-week old NOD *scid* gamma (NSG) mice (**Figure 2D**) and monitored tumor growth. ATGL KO decreased tumor growth and subsequently prolonged survival, which was rescued by the addback of ATGL WT (**Figures 2E,F**). IHC analyses of tumors revealed that ATGL KO increased apoptosis as detected via cleaved caspase-3 (CC3) and had a non-significant trend towards decreased proliferation as assessed by phosphorylated histone H3 (Ser10) (pHH3) levels (**Figures 2G-I**), effects that were reversed by the re-expression on ATGL WT. ATGL KO tumors were also noted to have large regions of necrosis that were not observed in control or ATGL WT addback tumors (**Figure 2G**).

### ATGL mediates the migration and invasion of prostate cancer cells

Lipid metabolism has been linked to prostate cancer metastasis^24,48–50^. Since these prior studies focused on the impact of extracellular lipids or *de novo* lipogenesis, we next wanted to test how regulating the access to stored lipids through the modulation of ATGL would influence prostate cancer cell migration and invasion. ATGL KO decreased CRPC cell migration in wound healing assays, an effect that could be rescued by re-expression of ATGL WT but often not ATGL S47A or S404A mutants and/or the addition of extracellular fatty acids (**Figures 3A-C; Supplementary Figures S7A-C**). Addition of extracellular fatty acids increased migration in ATGL WT-expressing C4-2 and C4-2B-LT cells (**Figures 4A-B; Supplemental Figures S7A-C**). Overexpression of ATGL (S404A) above basal levels (compared to expression nearer to endogenous ATGL levels (**Figure 1G**)) restored C4-2 migration (**Supplemental Figures S7D**), again indicating that ATGL total levels can supersede the impact of S404 phosphorylation. In addition, we also demonstrate here that knockdown of CAMKK2 and AMPK decreased AR-prostate cancer cell migration, similar to previous reports in AR+ prostate cancer cells^5^ (**Supplemental Figure S7E**).

**Figure 3:**
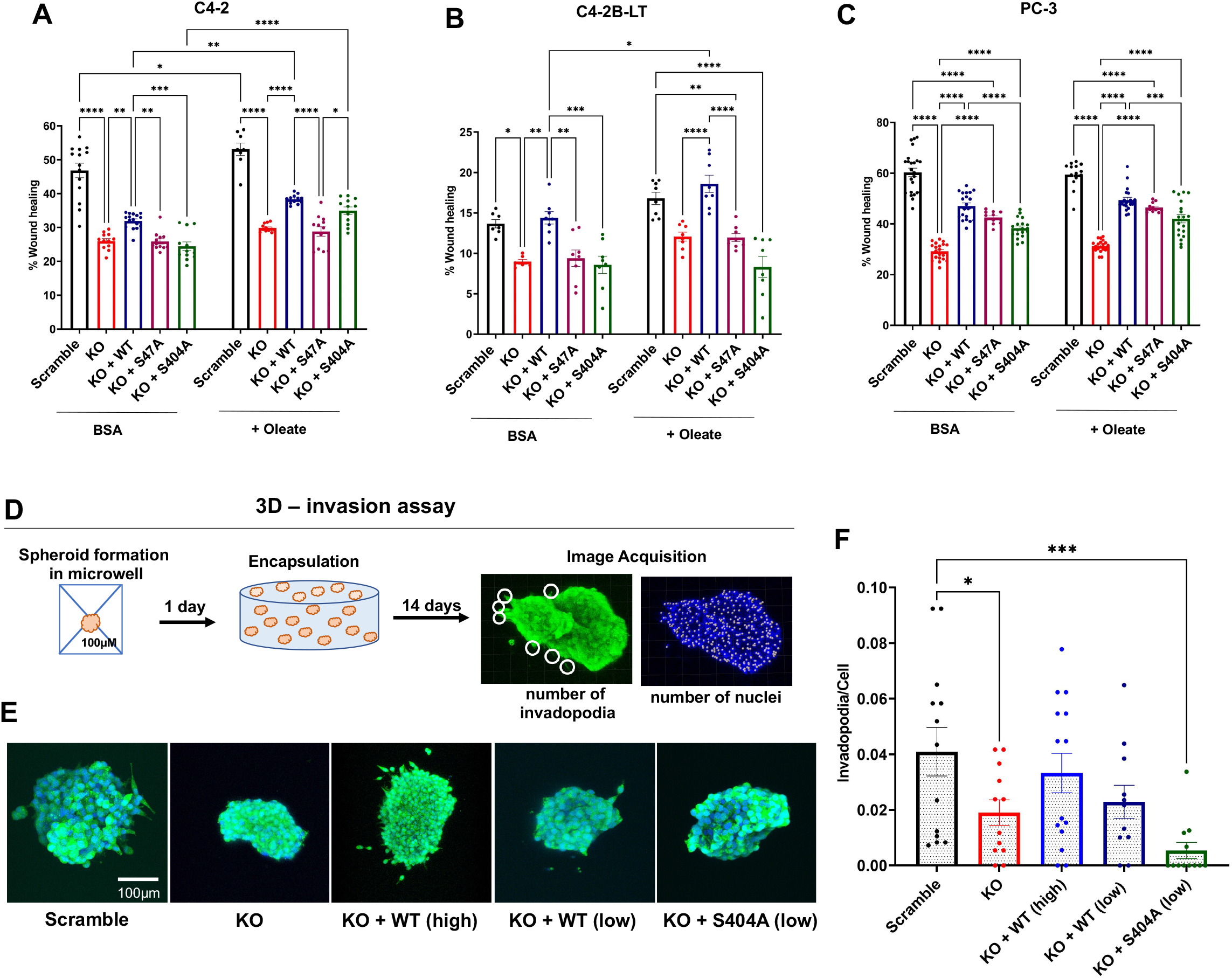
ATGL contributes to prostate cancer cell migration and invasion. Migration (wound healing/scratch test) of scramble, *PNPLA2* knockout (ATGL KO), ATGL KO with the addback of ATGL WT or S47A or S404A mutant ATGLs. Migration was analyzed over 24h in the presence of BSA or extracellular fatty acids (75 μM BSA-coupled oleate). Data are expressed as mean % scratch wound closure ± SEM. Data are representative results of at least three independent experiments. (D) Schematic of 3D invasion assay with a matrix metalloproteinase (MMP)-sensitive crosslinker embedded in hydrogel. (E) Representative images of clusters analyzed for invasion. *High* indicates overexpression, *low* indicates close to basal expression. (F) Quantification of invasion as invadopodia/cell. The graph combines the datapoints of five independent experiments. One-way ANOVA. \**P* < 0.05, \*\**P* < 0.01, \*\*\**P* < 0.001, \*\*\*\**P* < 0.0001, ns = no significance.

**Figure 4:**
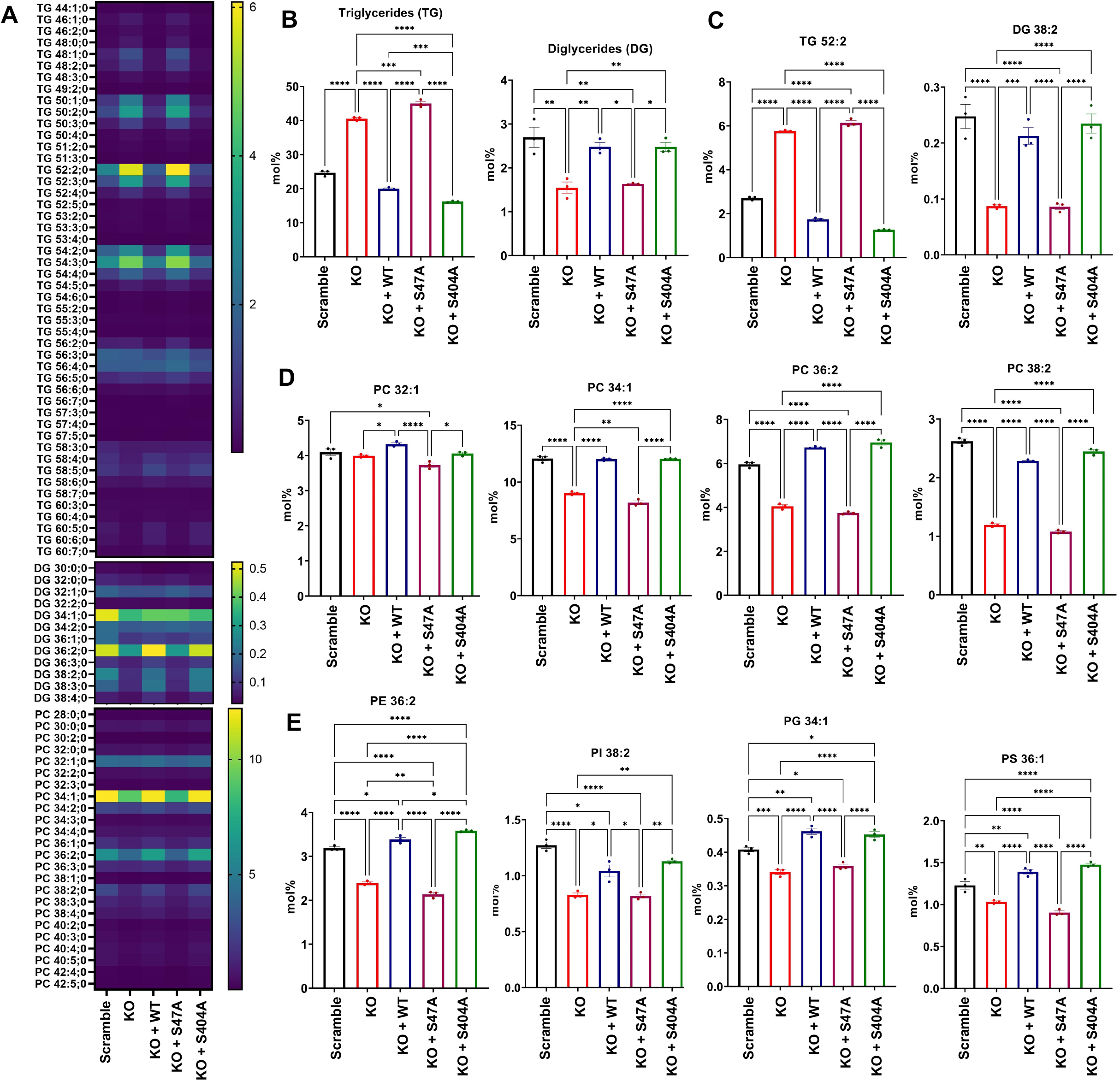
Shotgun lipidomics reveals ATGL lipolytic activity increases glycerophospholipids associated with prostate cancer. C4-2 cells with scramble sgRNA, *PNPLA2* knockout (ATGL KO) or ATGL KO with addback of ATGL wildtype (WT) or mutants (S47A, S404A) were analyzed via shotgun lipidomics. Cells were supplemented with 75 μM BSA-coupled oleate overnight. (A) Heatmaps of triglycerides (TG), diglycerides (DG) or phosphatidylcholine (PC) of detected lipid species in mol%. (B) Sum of TG and DG detected. (C) TG and DG lipid species with high abundance in samples detected. (D) Phosphatidylcholine lipids associated with prostate cancer in patient samples (Butler et al. 2021). (E) Additional lipid subclasses associated with prostate cancer (Butler et al. 2021): Phosphatidylethanolamine (PE), Phosphatidylinositol (PI), Phosphatidylglycerol (PG), Phosphatidylserine (PS). Sample size n = 3. One-way ANOVA. \**P* < 0.05, \*\**P* < 0.01, \*\*\**P* < 0.001, \*\*\*\**P* < 0.0001, ns = no significance.

Next, we tested the role of ATGL in a 3D model of migration and invasion (**Figure 3D**). Here, cells were grown in 3D spheroids in the presence of 5 mM glucose while encapsulated in a well-established hydrogel model, simulating an extracellular tumor matrix environment and allowing us to monitor invasion by determining the formed invadopodia per cell. ATGL KO decreased prostate cancer cell invasion, reflected by a decrease in the number of invadopodia, an effect that could be rescued by the overexpression of ATGL WT in a dose-dependent manner, but not the ATGL S404A mutant (**Figures 3E,F**). Together, these data indicate functional roles for ATGL and p-ATGL (S404) in CRPC cell growth, migration, and invasion.

### ATGL’s lipolytic activity increases the levels of malignancy-associated glycerophospholipids

Our functional data confirmed a role for ATGL in prostate cancer biological processes *in vitro* and *in vivo*. To better understand ATGL’s mechanism of action, we first determined if ATGL activity impacts membrane composition, since the released fatty acids from the lipid droplet can serve as precursors for glycerophospholipids, a major component of cell membranes^51^. To do so, we performed shotgun lipidomics with C4-2 ATGL KO and addback cell models treated for 24 hours with BSA-coupled oleate (75 μM) (**Figure 4; Supplemental Figures S8-10**). As expected, we observed an increase in detected TGs and decreases in diglycerides (DGs) upon ATGL knockout that could be reversed by re-expression of ATGL WT but not the catalytically inactive S47A mutant (**Figures 4A-C**). We did not observe notable differences in TG content when comparing ATGL WT and ATGL S404A mutant addbacks. This further supports our observation that ATGL’s phosphorylation at S404 has a modest effect on lipolytic activity that is mostly observed in the presence of higher fatty acid concentrations (ex. 150 μM). We also detected an increase in cholesterol esters (CE; **Supplemental Figures S8-9**) in the KO and ATGL S47A mutant addback cells, often observed when TGs are accumulated in the lipid droplet^52^. We observed clear decreases of several enriched glycerophospholipids in the ATGL KO and S47A mutant addback cells including phosphatidylcholines (PC), phosphatidylserines (PS), phosphatidylglycerols (PG), and phosphatidylinositols (PI) (**Figures 4D,E; Supplemental Figures S8-10**). This included several lipid subspecies that were also enriched in patients in the membranes of malignant prostate tumors compared to benign tissue^48^. One of the most prominent glycerophospholipid classes associated with prostate cancer is phosphatidylcholine. The most commonly found phosphatidylcholines associated with prostate cancer malignancy in patient samples (PC 32:1, PC 34:1, PC 36:2, PC 38:2)^48^ were clearly regulated by ATGL in our isogenic preclinical models (**Figures 4A,D**). Additional glycerophospholipids found to be increased in malignant prostate cancer patient samples^48^ (PI 36:1, PI 38:2, PS 36:1, PS38:2, PG 34:1, PG36:2, PE 34:1 and PE 36:2) likewise were regulated by ATGL (**Figures 4A,E, Supplemental Figures S8,10**). Collectively, shotgun lipidomics demonstrated that ATGL activity is a major regulator of phospholipid levels in prostate cancer cells. Notably however, phosphorylation of S404 did not influence ATGL lipolytic activity, at least with regards to the lipids profiled in this study. These data thus also suggest that S404 regulates another currently unknown function of ATGL that influences prostate cancer cell biology.

We next determined if the ATGL-mediated changes in metabolism observed in cell culture were maintained *in vivo*. To do this, we used mass spectrometry imaging (desorption electrospray ionization-mass spectrometry, DESI-MS) to spatially evaluate ATGL-mediated metabolic changes in xenograft tumors derived from the experiment described in Figures 2D-I (**Figure 5; Supplemental Figures S11-12**). This approach provides *in situ* spatial information on tumor metabolism that can be precisely correlated with tissue histopathology. Hence, DESI-MS imaging enables correlation between lipid distribution in specific tissue regions composed of cancer cells – apart from stroma and other neighboring cell compartments that may confound data analysis. As expected and observed in our shotgun lipidomics, DESI-MS revealed a clear increase in the relative abundance of structurally diverse TG species following ATGL KO, an effect that was reversed by the re-expression of ATGL (**Figure 5; Supplemental Figures S11-12**). Conversely, DGs were generally decreased or unchanged by ATGL KO. As observed in cell culture, we saw decreases in the malignancy-associated phosphatidylcholines PC 32:1, 34:1, 36:2 and 38:2 in ATGL KO tumors (**Figure 5**). This correlated with ATGL-mediated changes in choline and acetylcholine levels (**Supplemental Figures S11-12**). Additional changes were observed following ATGL knockout in PI 38:2, PS 36:1, PG 34:1 and PE 34:1 (**Figure 5**), which have all been associated with prostate cancer^48^ and were identified in our *in vitro* lipidomics (**Figure 4; Supplemental Figures S8,10**). DESI-MS was also able to detect ATGL-dependent changes in the relative abundances of signaling lipids derived from glycerophospholipids, including lysophosphatidylcholines, lysophosphatidylethanolamines and lysophosphatidylglycerols (**Supplemental Figure S12**). Interestingly, the relative abundance of longer-chain monoglycerides and fatty acids were increased in scramble and ATGL WT tumors relative to ATGL KO tumors, whereas short-chain species were less abundant or unchanged (**Supplemental Figure S12**).

**Figure 5:**
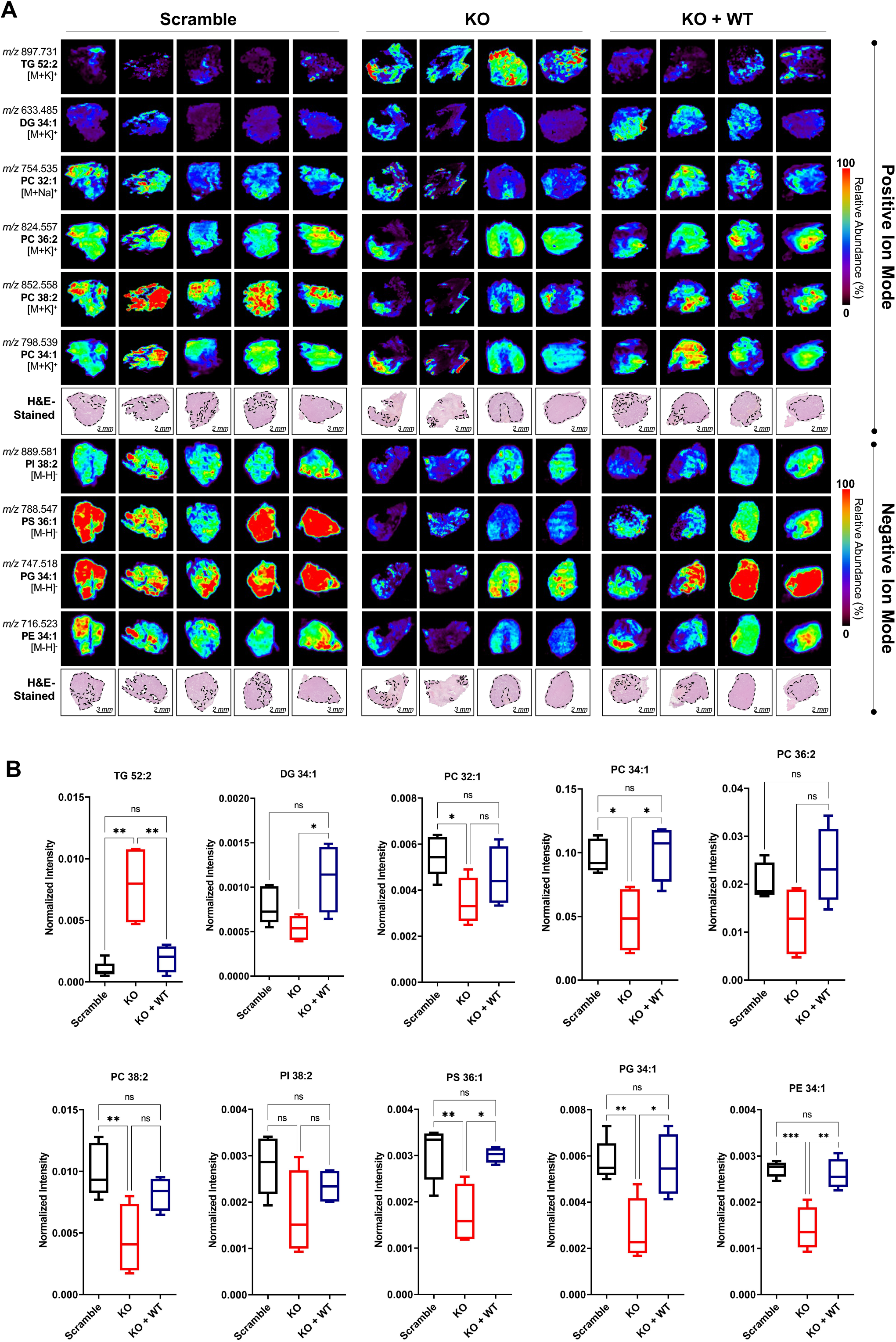
DESI-MS analysis of human CRPC xenografts confirms findings of *in vitro* shotgun lipidomics. (A) Representative DESI-MS images from human CRPC xenografts described in Figure 2. (B) Normalized ion intensities for malignancy-associated phosphatidylcholine (PC), phosphatidylinositol (PI), phosphatidylserine (PS), phosphatidylglycerol (PG), and phosphatidylethanolamine (PE) lipids subclasses detected in the human CRPC xenografts. Scramble (n = 5), KO (n = 4), KO + WT (n=4). One-way ANOVA and Tukey. \**P* < 0.05, \*\**P* < 0.01, \*\*\**P* < 0.001, \*\*\*\**P* < 0.0001, ns = no significance.

### Pharmacological or molecular targeting of ATGL inhibits growth in diverse preclinical models of prostate cancer

Our genetic data indicate that ATGL is required for prostate cancer cell migration, invasion, proliferation, colony formation, and tumorgenicity. As such, we next wanted to assess ATGL as a potential therapeutic target in advanced prostate cancer. To test this, we used the ATGL-specific inhibitor atglistatin^54^. While atglistatin is highly selective for ATGL over other lipases, it is specific to murine ATGL^55,56^ and will not inhibit human ATGL. Therefore, we tested the efficacy of atglistatin in murine prostate cancer models. Consistent with our genetic data in human cells, atglistatin increased neutral lipids as assessed by Oil Red O staining in murine RM-9 prostate cancer cells (**Figure 6A; Supplemental Figure S13A**). In addition, atglistatin decreased prostate cancer colony formation and cell migration in a dose-dependent manner (**Figures 6B-C**). Glucose starvation again sensitized cells to ATGL inhibition in both cell growth and colony formation assays (**Figures 6B,G; Supplemental Figure S13B**). To test the effect of ATGL inhibition on invasion, we subjected RM-9 3D-cell clusters embedded in hydrogels to atglistatin and measured the number of escaped cells per cell cluster following a 5-day treatment, which resulted in a decrease of escaped cells (**Figure 6D**).

**Figure 6:**
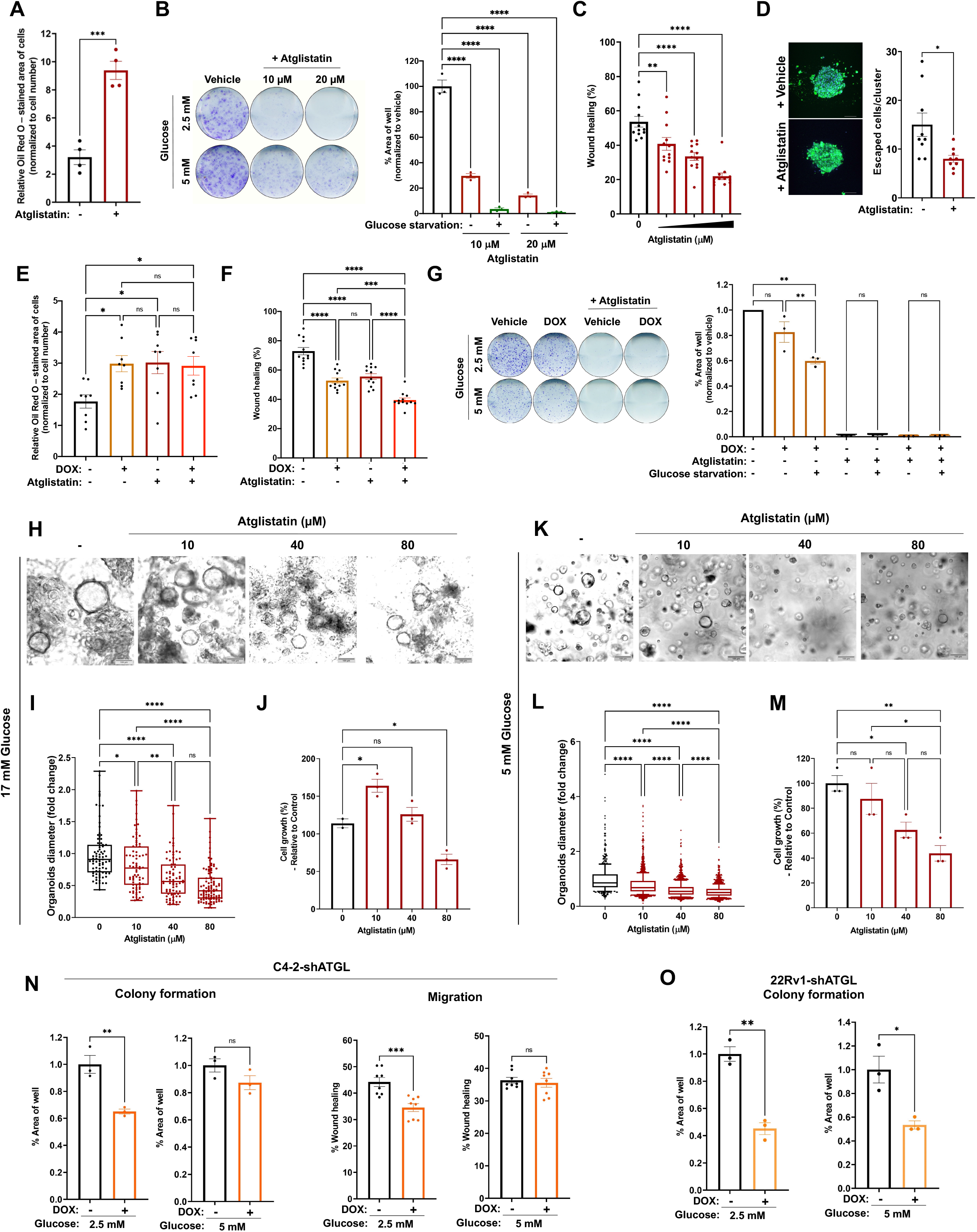
ATGL can be molecularly or pharmacologically targeted in diverse prostate cancer models. (A) Murine prostate cancer cells (RM-9) stained with Oil Red O to determine the TG content upon overnight treatment with atglistatin (40 μM). (B) Colony formation of RM-9 cells grown in RPMI 1640 with adjusted glucose concentrations (5 mM or 2.5 mM). (C) Migration (wound healing/scratch assay) of RM-9 cells quantified after 24h treated with atglistatin (20, 40 and 80 μM) (D) 3D invasion assays using a hydrogel with MMP sensitive crosslinker to assess RM-9 invasion (escaped cells from cluster) upon treatment with 40 μM atglistatin. (E-G) RM-9 inducible shRNA constructs targeting scramble control or *PNPLA2* (ATGL) quantified for (E) TG accumulation via Oil Red O staining (F) migration (wound healing/scratch assay) and (G) colony formation, upon knockdown and/or treatment with 10 μM atglistatin. (H-M) Organoid studies using Hi-MYC model-derived organoids grown in (H-J) typical organoid media containing high glucose (17 mM) or (K-M) adjusted glucose concentrations (5 mM). (N) Knockdown of *PNPLA2* (shATGL) via doxycycline (DOX)-inducible shRNA decreases C4-2 colony formation and migration in a glucose-dependent manner. (O) Knockdown of ATGL in 22Rv1 cells decreases colony formation. Note, the migration of 22Rv1 cells was too low to quantify over 24 hours. One-way ANOVA. \**P* < 0.05, \*\**P* < 0.01, \*\*\**P* < 0.001, \*\*\*\**P* < 0.0001, ns = no significance.

A previous report in lung cancer cells observed opposing effects after atglistatin treatment compared to knockdown of ATGL^33^, suggesting the potential for off-target effects. To test for potential off-target effects, we created stable RM-9 cells in which we could silence ATGL in a doxycycline (DOX)-dependent manner (**Supplemental Figure S13C**). Both ATGL knockdown and atglistatin increased neutral lipid accumulation and decreased cell migration and colony formation (**Figures 6E-G; Supplemental Figure S13D**). Moreover, the effects of ATGL knockdown and atglistatin appeared mostly redundant or at most additive, consistent with atglistatin’s effects being on-target (**Figures 6E-G; Supplemental Figure 13D**). As an additional control, doxycycline-mediated induction of scramble control (shControl) did not impact neutral lipid levels, migration, or colony formation, indicating that the phenotypes observed are not due to doxycycline-mediated off-target effects (**Supplemental Figures S13E-G**). Notably, ATGL knockdown *increased* colony formation when cells were switched to supraphysiological levels of glucose (25 mM) often found in cell culture media (ex. DMEM) (**Supplemental Figure S13H**), which may help explain prior discrepancies in the field. To test if ATGL could be targeted in an orthogonal, more physiological setting, we treated organoids derived from Hi-MYC genetically engineered mice with atglistatin and observed a dose-dependent decrease in organoid diameter and cell growth, the efficacy of which was again increased when glucose concentrations were lowered from standard high-glucose organoid media (17 mM glucose) to physiological (5 mM) levels (**Figures 6H-M**). Lipidomics performed on the organoids revealed that atglistatin again decreased the levels of malignancy-associated phosphatidylcholines^48^ such as PC32:1, PC34:1 when the media was adjusted to physiological glucose levels (**Supplemental Figure S14**).

The species specificity of atglistatin limited testing of this compound to mouse models. To test the efficacy of temporal ATGL inhibition in human cells, we created human prostate cancer cell lines containing DOX-inducible shRNAs targeting scramble control (shControl) or *PNPLA2* (shATGL) (**Supplemental Figures S15-16**). Inducible knockdown of ATGL, but not scramble control, impaired C4-2 and 22Rv1 CRPC colony formation and migration, effects that were again magnified in low glucose conditions (**Figures 6N-O, Supplemental Figure S15**). Interestingly, knockdown of ATGL in hormone-sensitive LNCaP cells had minimal effects on basal and androgen-mediated colony formation (**Supplemental Figure S16**), consistent with a heightened role for ATGL in CRPC relative to HSPC.

### Inhibition of ATGL creates a therapeutically targetable metabolic shift towards glycolysis

Because ATGL inhibition decreased migration and proliferation/colony formation, we wanted to further understand the contribution of ATGL to prostate cancer metabolism. ATGL’s effects on prostate cancer biology were influenced by glucose concentration (**Figures 2 and 6**). Previous studies have described a metabolic shift towards glycolysis upon ATGL inhibition^57^ but also upon ATGL activation^58,59^. Due to the conflicting reports in adipose and other non-prostate tissues, we wanted to assess how ATGL inhibition impacted glycolysis in prostate cancer cells. We first tested our C4-2 ATGL KO and addback cell lines using metabolic flux analysis (Seahorse™ glycolysis stress test) and observed an increase in the extracellular acidification rate (ECAR), which correlates with glycolysis, in ATGL KO cells (**Figures 7A-B**). Like in cell culture, ATGL KO increased intratumoral levels of glucose and lactate, indicating that blockade of ATGL caused a similar metabolic shift towards glycolysis *in vivo* (**Figures 7C-D**). Pharmacological inhibition of murine prostate cancer cells with atglistatin also increased ECAR, which could be reversed upon treatment with the glycolysis inhibitor AZ-PFKFB3-26^60,61^ (**Figures 7E-F**). Together, these results indicated that targeting ATGL in CRPC creates a metabolic shift towards glycolysis that we hypothesized could be therapeutically exploited. To test this idea, we co-treated prostate cancer cells with atglistatin and AZ-PFKFB3-26 and assessed the effects on colony formation (long-term treatment) (**Figure 7G**). Coefficient of drug interaction (CDI) scores were calculated to be 0.563 between treatments suggesting a synergistic effect (CDI values <1) between the two treatments (**Figure 7G**). We also performed short-term (4 days) treatment cell growth assays with increasing doses of atglistatin and AZ-PFKFB3-26, which resulted in a highest single agent (HSA) combined synergy score of 8.133 with maximal scores (>15) between 1-2 μM AZ-PFKFB3-26 and 40 μM atglistatin (**Figures 7H-I**). Thus, glycolysis inhibitors further sensitize CRPC cells to ATGL inhibitors and vice versa. As such, the combination of glycolytic and ATGL inhibitors represents a potential novel therapeutic approach that we propose warrants further investigation.

**Figure 7:**
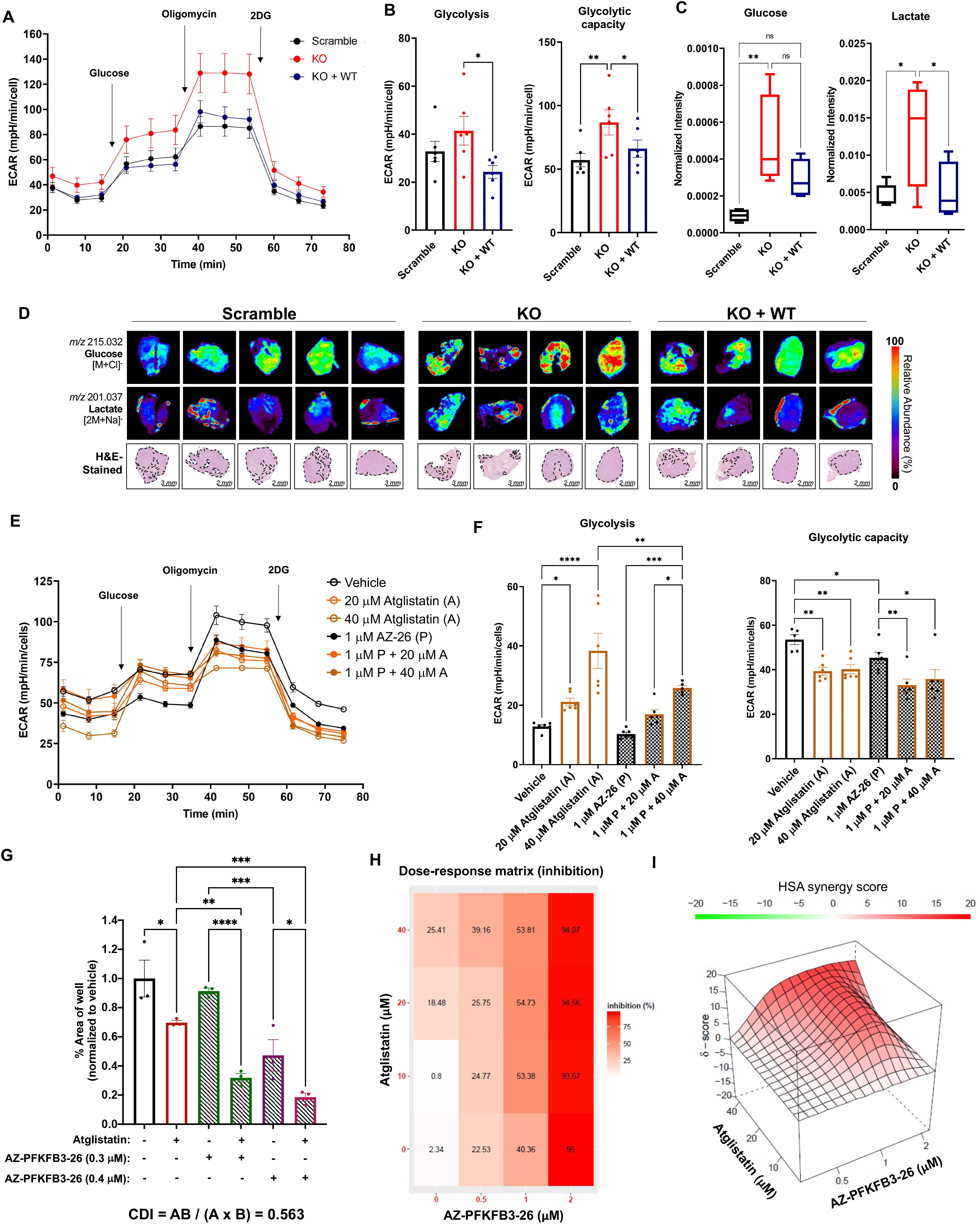
ATGL KO or inhibition creates a metabolic switch towards glycolysis that can be therapeutically exploited. (A-B) Metabolic flux analysis of C4-2 Scramble, *PNPLA2* knockout (ATGL KO), and ATGL KO with ATGL wildtype (WT) addback. (A) ECAR of glycolytic stress assay, (B) Glycolysis (ECAR measurement after glucose injection but before oligomycin injection), and glycolytic capacity (maximum ECAR after oligomycin injection). (C-D) CRPC xenografts demonstrate an increase in glucose and lactate upon *PNPLA2* knockout (ATGL KO), which can be rescued by ATGL addback. (E-F) Metabolic flux analysis of RM-9 cells treated for 24h with atglistatin (A) and the glycolytic inhibitor AZ-PFKFB3-26 (AZ-26; P). (E) ECAR of glycolytic stress assay which was used to quantify (F) glycolysis and glycolytic capacity. (G) RM-9 cells co-treated with atglistatin and AZ-PFKFB3-26 had decreased colony formation and resulted in a calculated coefficient of drug interaction (CDI) of 0.563. (H-I) Short term co-treatment of atglistatin and AZ-PFKFB3-26 was used to calculate synergy. One-way ANOVA. \**P* < 0.05, \*\**P* < 0.01, \*\*\**P* < 0.001, \*\*\*\**P* < 0.0001, ns = no significance.

## Discussion

ATGL’s role in cancer has remained enigmatic due to conflicting reports across different cancer types. In lung cancer, ATGL has been described as a tumor suppressor based on the early observation that ATGL expression is reduced in lung cancer^34^. This study reported that 25% of the *Pnpla2/ATGL* knockout mice spontaneously developed lung adenocarcinoma, while ATGL WT mice were cancer-free^34^. Two additional *in vitro* studies suggested that the knockout of ATGL in lung cancer results in an increase in lung cancer cell migration and proliferation^57,62^. However, only a single cell line was used for both studies and a prior study found that inhibition of ATGL in the same cell line decreased proliferation and migration^63^. A more comprehensive study looked at the role of ATGL in different cancer cell lines, including lung, melanoma, colon, and liver^33^. While ATGL overexpression had tumor suppressive functions, shRNA-mediated knockdown of ATGL did not have any effects *in vitro* or *in vivo*^33^. Adding to the confusion, in pancreatic cancer, ATGL levels are increased in obese patients in both the cancer and surrounding stroma and associated with worse outcome^64^. Another study found that ATGL has a pro-tumor role in hepatocellular carcinoma in preclinical models, since the knockdown of ATGL reduced tumor size, unless supplemented with free fatty acids, and overexpression of ATGL increased tumor volume^65^. In prostate cancer, the ATGL coactivator CGI-58 has a tumor suppressive function^36,66^. However, CGI-58’s role as a tumor suppressor was uncoupled from ATGL^36^. Notably, CGI-58, which is known to activate murine ATGL was shown to only have a minor impact on human ATGL’s activity^33,67^. Prior studies utilizing prostate cancer cell culture models found that ATGL knockdown increased neutral lipids but had only minor effects on proliferation compared to DGAT1^35,68^. We speculate that the modest effects of ATGL in these studies may be attributable to the high glucose concentrations used in standard cell culture media that don’t reflect physiological levels of glucose in the tumor microenvironment. Finally, in contrast to the above findings, a previous study found a pro-tumor role for CGI-58^35^. Together, these data indicate potential context-dependent roles for ATGL that might be influenced by cancer type, disease stage, and/or experimental conditions. To the latter, our data suggests that using culture conditions that better mimic the nutrient-challenged tumor microenvironment is critical to assessing ATGL’s role in cancer. Our findings also raise the question of whether proteins such as ATGL have been overlooked in large CRISPR or RNAi loss-of-function screens due to the use of high glucose- and serum-containing media.

The data presented here provide additional insights into the regulation of human ATGL’s lipolytic activity. This study started with the observation that ATGL is highly phosphorylated at S404 in mCRPC patient samples (**Figure 1**). The orthologous phosphorylation site was previously described in murine adipose tissue to be phosphorylated by AMPK, leading to an increase in its lipolytic activity^41^. Since there is 87.5% homology between murine and human ATGL, it was largely assumed that this lipolytic-stimulating event would be conserved between species. However, this was never investigated in human cells using a human ATGL construct. Accordingly, murine-specific p-ATGL S404-targeted antibodies were used to determine changes in the phosphorylation of human ATGL. While we detected bands at the correct size for human p-ATGL using the most prominent, commercially available murine-specific p-ATGL (S406) antibody, siRNA-mediated knockdown of ATGL indicated that this antibody cannot recognize human p-ATGL (S404). We therefore created a new human-specific p-ATGL (S404) antibody and demonstrated that the AR-CAMKK2-AMPK axis phosphorylates this site in prostate cancer. To our knowledge, this is the first time that human ATGL has been shown to be phosphorylated by AMPK at S404 in human cells. In contrast to murine ATGL, we found that the phosphorylation of ATGL by AMPK at S404 only leads to a modest increase in lipolytic activity, observed most at high concentrations of extracellular fatty acids (150 μM). By comparison, expression of ATGL appears to have a more prominent role in lipid metabolism. This is in line with previous findings investigating ATGL in non-adipose tissue, where the overexpression of human wildtype, but not lipolytically inactive (S47A), ATGL, decreased lipid size, suggesting that total ATGL activity may correlate best with ATGL expression^39^. However, this study did not investigate the S404 site and its impact on ATGL lipolytic activity. Interestingly, beyond ATGL KO and expression of a catalytically inactive mutant of ATGL (S47A), CRPC cells expressing the ATGL (S404A) mutant exhibited decreased colony formation and invasion/migration compared to ATGL WT-expressing cells. These data suggest that the S404A mutation, in addition to impairing ATGL’s lipolytic activity in the presence of excess lipids (ex. 150 μM oleate), disrupts an important function of ATGL that is independent of its function in the breakdown of triglycerides. In addition to AR+ CRPC models, we also confirmed ATGL-dependent effects in AR-prostate cancer cell models. Likewise, CAMKK2 and AMPK inhibition also decreased AR-prostate cancer cell growth and migration, indicating that CAMKK2-AMPK signaling could be activated in prostate cancer cells independent of AR. Of note, disruption of ATGL had no effect on intracellular neutral lipid levels in non-transformed prostate epithelial cells and colony formation in hormone-sensitive prostate cancer cells, indicating that ATGL’s pro-cancer roles are more specific to advanced prostate cancer (**Supplemental Figures S2,16**). Additional functions of ATGL were described when it was first discovered including CoA-independent acyl-glycerol transacylase activity (two monoglyceride to one diglyceride or one monoglyceride and one diglyceride to one triglyceride)^23^ and phospholipase A2 activity^69^. However, these additional enzymatic activities have been poorly studied. ATGL has also been described to play a role in autophagy/lipophagy^70^. Given the previous reports on the role of autophagy in CRPC^20^, it would be of interest to test if ATGL plays a role in this process, and if the S404A mutant interferes with this function. Recently, a novel non-lypolytic function of ATGL was described to facilitate the biosynthesis of branched fatty acid (FA) esters of hydroxy FAs^71^. Our lipidomic platforms did not explore these lipid species, but certainly this is an area of interest for future studies. A major challenge in the study of ATGL structure-function is that there is currently no published ATGL structure^72^. Further, structure prediction models such as alpha fold^73^ cannot predict the structure of ATGL, and in particular the C-terminus where the S404 site is located. Most understanding about the role of ATGL’s C-terminus comes from mutational studies based on isolated murine and human ATGL, which suggest that mutations could impair its activity^74^. Therefore, it is challenging to determine if the ATGL S404A mutant would lead to changes in other protein binding partners based on our current knowledge about this protein’s structure. A caveat of our work is that we did not examine any additional phosphorylation sites of ATGL that could, for example, interfere with its location as suggested in murine ATGL^75^. We also did not test additional upstream kinases of ATGL that have been reported to phosphorylate murine ATGL at S406, such as PKA^76^.

ATGL inhibition caused a shift towards glycolysis *in vitro* and *in vivo* (**Figure 7**). This phenotype has been observed in lung cancer cells^57^. This finding was important in the context of prostate cancer because while many types of cancer have been characterized as highly glycolytic, this is not the case for most prostate cancers^77^.

Our data indicate that the co-targeting of ATGL-mediated lipid metabolism and glycolysis could represent a novel approach to overcome the metabolic plasticity of mCRPC, a disease stage that currently has no cure.

One of the major concerns in the systemic targeting of ATGL is its potential impact on the heart. Systemic deletion of *Pnpla2* leads to premature lethality unless ATGL expression is rescued in the heart^78–82^. However, systemic targeting of ATGL in mice did not show any signs of cardiomyopathy during long-term treatment with atglistatin^56^. While more studies are needed to evaluate the potential side effects of targeting ATGL systemically in the context of prostate cancer, it also still needs to be determined if enough of the inhibitor would reach the tumor. In our preclinical models, concentrations of atglistatin lower than those used in adipocytes to block ATGL lipolysis (80 μM) were sufficient to decrease prostate cancer colony formation, migration, invasion, and organoid growth (**Figure 6**). Whether an inhibitor of ATGL can safely block human prostate cancer progression *in vivo* remains to be tested. While preparing this manuscript, the first human specific ATGL inhibitor NG-497 was published^83^, which will allow further investigation for the potential treatment of mCRPC. To our knowledge, however, NG-497 has not yet been shown to have activity *in vivo*.

In this study we demonstrate clear pro-cancer roles for ATGL in advanced prostate cancer using multiple murine and human 2D and 3D models, as well as human xenograft mouse models. Moreover, early studies in mice have demonstrated that the systemic targeting of ATGL can counteract models of induced insulin resistance^56^, a known co-morbidity of anti-AR therapy^84,85^. As such, ATGL inhibitors could have dual benefit, targeting both the cancer itself and standard of care treatment-associated comorbidities.

## Acknowledgments

We would like to thank Kelly Kage (UT MD Anderson Cancer Center) for artistic assistance in preparing the graphic abstract. We want to thank Dr. Sue-Hwa Lin (UT MD Anderson Cancer Center) and Dr. Mikhail Kolonin (UT Health Science Center Houston) for helpful conversations about lipid metabolism in prostate cancer. We would like to thank the MD Anderson Cancer Center Functional Genomics Core (Houston, TX, USA) for the shRNA constructs and the MD Anderson Cancer Center Science Park Research Histology, Pathology, and Imaging Core (RHPI) (Smithville, TX, USA) for histology services.

This work was supported by grants from the National Institutes of Health (NIH R01CA184208 and P50CA140388 (D.E.F.); P30CA125123, P30ES030285, P42ES0327725 (C.C.); P01CA098912 (M.C.F-C)), American Cancer Society (RSG-16-084-01-TBE (D.E.F.), CPRIT (RP170005 and RP200504 (C.C.)), and a grant from the Department of Defense/Prostate Cancer Research Program (W81XWH-22-1-0686 (D.E.F.)). This work was also supported by an American Legion Auxiliary Fellowship (D.A.), the Austrian Scientist in North America Mentoring Program (D.A.), and the Larry Deaven PhD Fellowship in Biomedical Sciences (T.V.T.). Some of the histology was performed with the CCSG-funded MDACC Research Histology Core Laboratory, NIH grant P30CA016672.

## Author Contributions

Conceptualization, D.A. and D.E.F.; Investigation: D.A., M.S., T.L.P, T.V.T., C.F.R., H.P., E.S., J.J.A., M.M.M., J.J., B.P., C.C., M.C.F-C., M.L., L.S., D.E.F; formal analysis: D.A., M.S., T.L.P., T.V.T., C.F.R., H.P., E.S., B.P., C.C., M.C.F-C., M.L. and D.E.F; resources: C.F., M.L, L.S.E, D.E.F; data curation: D.A. and D.E.F.; writing—original draft preparation, D.A., and D.E.F.; writing—review and editing, D.A., M.S., T.L.P., T.V.T., C.F.R., P.H.A.C, H.P., E.S., J.J.A, M.M., J.J.H., J.M.D., B.P., C.C., M.M.I., M.C.F-C., M.L. and D.E.F; resources: C.F., M.L, L.S.E, J.M.D., D.E.F; visualization, D.A. and D.E.F.; supervision: C.C., M.C.F-C., M.L., L.S.E. and D. E.F.; All authors have read and agreed to the published version of the manuscript.

## STAR Methods

### Institutional Review Board Statement

The animal study protocol was approved by the Institutional Review Board of the University of Texas MD Anderson Cancer Center (IACUC protocol#: 00001738-RN01, approved 10/4/2021).

### Bioinformatics

Phosphoproteomic data^37^ was mined using an updated AMPK substrate motif and PWM^38^. AMPK targets were ranked by PWM score, *P* value, and change in phosphorylation. A complete list can be found in Supplemental File 1.

### Cell lines

RWPE-1, LNCaP, C4-2, PC-3, 22Rv1, and NCI-H660 were obtained from the American Type Culture Collection (ATCC, Manassas, VA, USA) and cultured according to ATCC protocols. C4-2B and C4-2B-LT (C4-2B cells stably expressing firefly luciferase and tdTomato) (gifts from Dr. Sue-Hwa Lin (UT MD Anderson Cancer Center)), MDA-PCa-144-13 (gift from Dr. Nora M. Navone (UT MD Anderson Cancer Center)), and RM-9 cells (gift from Dr. Timothy Thompson (UT MD Anderson Cancer Center)) were cultured as previously described^86,87^. Cell lines were maintained without any addition of antibiotics, authenticated using short tandem repeat analyses, and tested for mycoplasma upon thawing out fresh vials.

### CRISPR knockout and addback strategies

Single guide RNAs (sgRNA) were designed using the Dharmacon CRISPR design tool (2018) and tested for off-target effects using ChopChop (https://chopchop.cbu.uib.no/2019). Two sgRNAs targeting Exon2 of *PNPLA2* (sgRNA1: 5’-GATGTTCCACGTCTTCTCGC-3’ (-)-strand, sgRNA2: 5’-CGCCGACGTAGTAGACGCCG-3’ (-) strand) and Exon 3 (sgRNA3 5’-TTGAGGTATCTAAAGAGGCC-3’) and a scramble sgRNA (https://doi.org/10.1038/nature20565) were ordered via Sigma as oligonucleotides. SgRNAs were then cloned into the pLentiCRISPRv2 as previously described^88^. The different pLentiCRISPRv2 constructs containing each sgRNA were virus packaged using HEK293 (Lentviral packaging, 3^rd^ generation) and then transfected into different prostate cancer cell lines and selected using puromycin. Prostate cancer cell lines containing the scramble sgRNA were used as a pool. For each of the three different sgRNAs, several single colonies were confirmed via sequencing, immunoblot, and then tested for proliferation, migration and TG accumulation using Oil Red O staining.

**Table 1:**
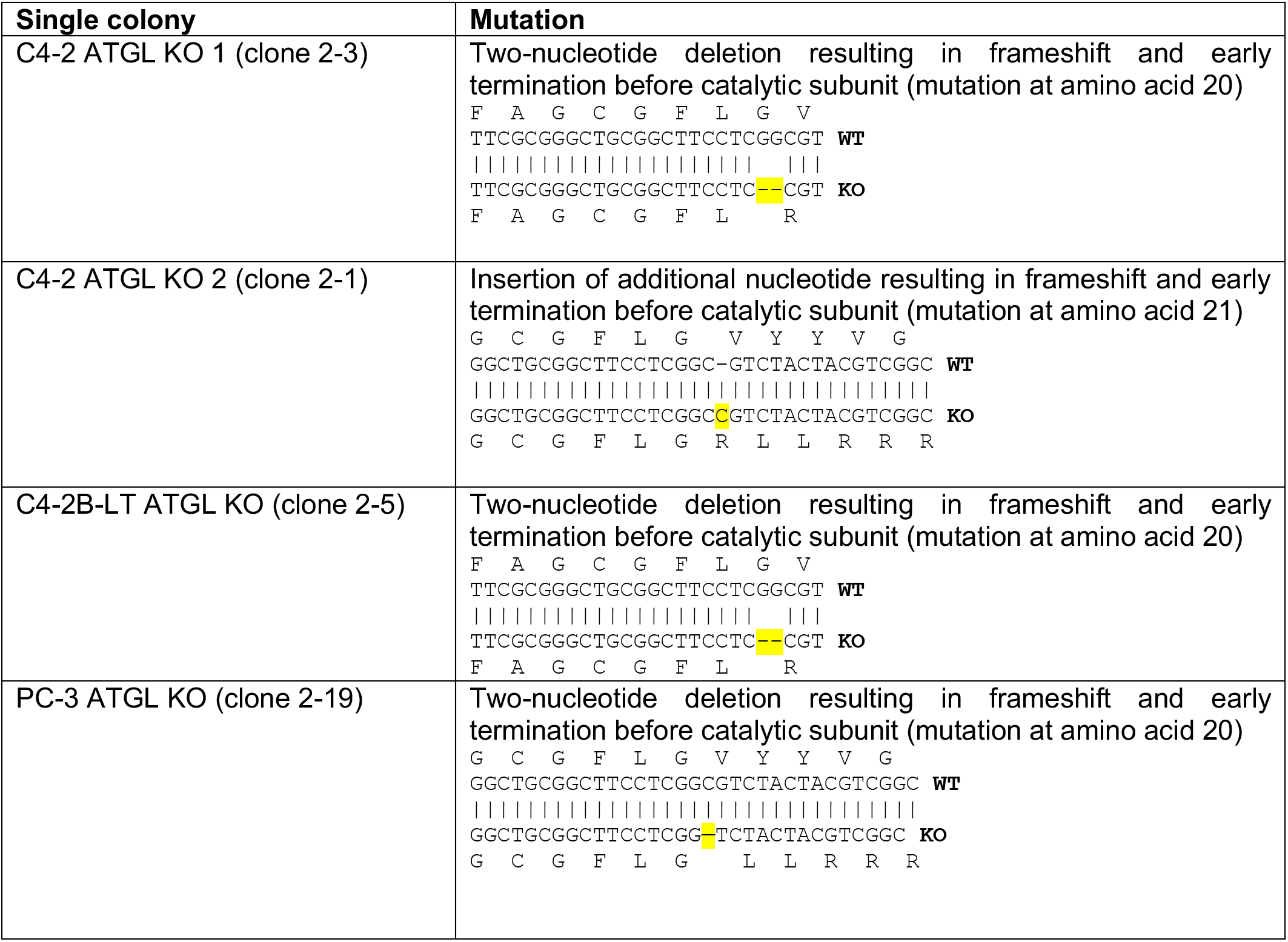
ATGL KO cell lines used in this study.

Addback constructs were created by first mutating the PAM sequence used by the sgRNA for the ATGL knockout. This was achieved by site direct mutagenesis in the pDONR223-*PNPLA2* plasmid using primers containing a mismatch, resulting in the mutation of nucleotide 51 (C to T) while maintaining the amino acid residue (F). In addition to the PAM mutation, we also created a S404A (TCG to GCG) mutation and a S47A (TCG to GCG) mutation using site-directed mutagenesis. Constructs were then inserted into the pLenti-PGK-Neo-DEST backbone via an LR reaction (Gateway™ LR Clonase™ II Enzyme mix, Cat#11791020, Invitrogen, Waltham, MA, USA) virus packaged in HEK293T cells and then stably integrated into the ATGL KO single colonies via G418 selection.

### shRNA cell lines

pGIPZ-shRNA constructs were purchased from the MDACC Functional Genomics Core and cloned via XhoI (Cat#R0146, NEB, Ipswich, MA, USA) and MluI (Cat# R0198, NEB, Ipswich, MA, USA) into pINDUCER10 (gift from Thomas Westbrook). Cells were transfected viral lentiviral packaging and selection via puromycin (2 μg/ml). For human prostate cancer cell lines seven different shRNAs and for murine prostate cancer cell lines three different shRNAs targeting ATGL were tested. The shRNA construct that showed the best knock down efficiency via immunoblot was selected for further experiments.

**Table 2:**
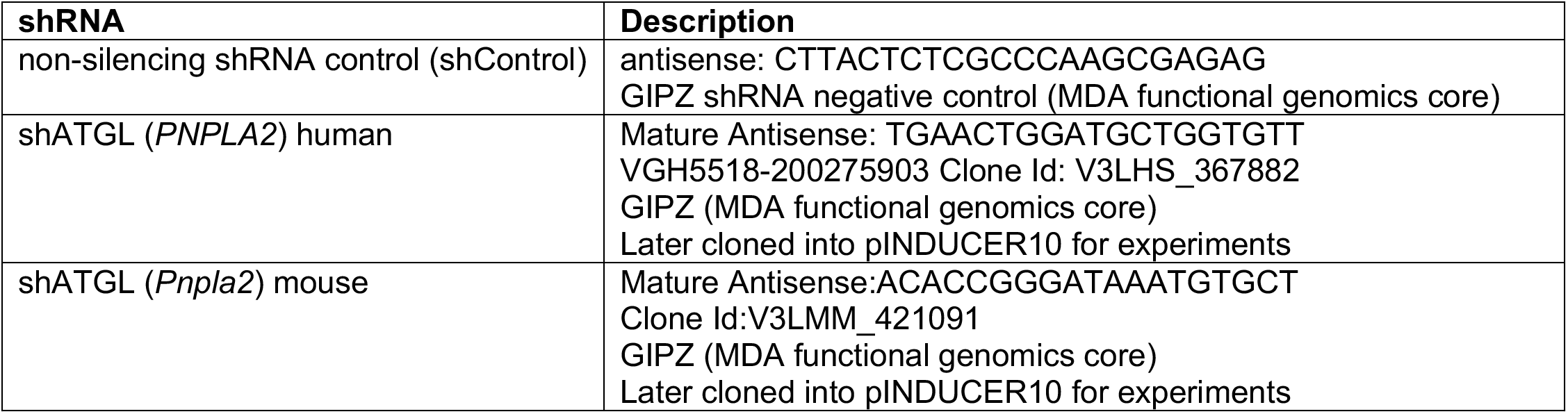
List of shRNAs used in this study.

### siRNA treatment

Chemical siRNAs (Silencer select) were obtained via Invitrogen. For immunoprecipitation, cells were plated at 60% confluency in RPMI 1640 (no glucose) supplemented with 5 mM glucose and 5%dialyzed FBS. Cells were transfected with silencer RNAs for 72 h before cells were collected as described in the immunoblot section.

**Table 3:** List of siRNAs used for the ATGL study.

| siRNA | Target | Catalog number<br>(Invitrogen, Waltham, MA, USA) |
| --- | --- | --- |
| select negative control #1 | - | Cat#4390843 |
| silencer select *CAMKK2*<br>(labeled as #1 in here) | CAMKK2 | assay id: S20927, validated,<br>Cat# 4390824 |
| silencer select *CAMKK2*<br>(labeled as #2 in here) | CAMKK2 | assay id: s20926, validated,<br>Cat# 4390824 |
| silencer select *PRKAA1* | AMPK alpha subunit 1 | assay id: s100, Cat # 4392420 |
| silencer select *PRKAA1* | AMPK alpha subunit 1 | assay id: s101, Cat # 4392420 |
| silencer select *PRKAA2* | AMPK alpha subunit 2 | assay id: s11056, Cat # 4390824 |
| silencer select *PRKAA2* | AMPK alpha subunit 2 | assay id: s11057, Cat # 4390824 |
| silencer select *PNPLA2*<br>(labeled here as #1) | PNPLA2 (ATGL) | assay id: s32682, Cat # 4392420 |
| silencer select *PNPLA2*<br>(labeled here as #2) | PNPLA2 (ATGL) | assay id: s32684, Cat # 4392420 |

### Colony formation assays

5000 cells were plated in RPMI 1640 (no glucose) supplemented with 5% (LNCaP, C4-2, C4-2B-LT) or 0.5% (PC-3, RM-9, 22Rv1) dialyzed FBS and glucose adjusted as noted in figures. Cells were fed every 48 h with media or retreated with inhibitors as noted in figures. Assays were run for various lengths of time (ex. low glucose conditions took longer to form colonies) but data were always normalized to controls on the same plate series. Cells were fixed using 10% formalin (Cat# HT501128, Sigma, St. Louis, MO, USA) and then washed with PBS. After drying overnight, fixed cells were stained with crystal violet (0.0125% dissolved in water) and washed with water until excessive crystal violet solution was removed. The plate was then dried and scanned at a resolution of 600 dpi. The colony formation assay was quantified using ImageJ by converting the image into grayscale, processing to binary, and then measuring the area % per well that is occupied by the colonies.

### Wound healing (scratch test) assays

Cells were plated in 6-well plates at 80-90% confluency in RPMI 1640 (no glucose) supplemented with 5% (LNCaP, C4-2, C4-2B-LT) or 0.5% (PC-3, RM-9, 22Rv1) dialyzed FBS, as well as 5 mM glucose, unless otherwise indicated. If supplemented with fatty acids, cells were incubated 48h before scratch and then pretreated the day of scratch. Doxycycline-inducible shRNA-containing cell lines were pretreated in flasks with 600 ng/ml doxycycline for at least 7 days prior to the assay. RM-9 or RM-9 shRNA cell lines were treated with atglistatin (cat#5301510001, Calbiochem, Sigma, St. Louis, MO, USA) immediately after the scratch (t=0). Scratch was performed using a 200 μl pipette tip. Images were taken with a Cytation 5 (BioTEK, Serial # 1705171D, 4xPL FL, Phase contrast) at timepoint 0 h and 24 h. Gaps from scratch were quantified via ImageJ using the MRI wound healing feature. Area of gap was recorded and then the % area wound healing was calculated using the formula 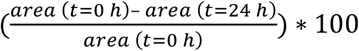.

### Cell viability assays

NCI-H660 and MDA-PCa-144-13 cells were plated in RPMI-1640 media containing 5 μg/ml insulin, 1 μg/ml transferrin, 30 nM sodium selenite, 10 nM hydrocortisone, 10 nM beta-estradiol, 4 mM L-glutamine, 10 mM HEPES, 1 mM sodium pyruvate and 5% FBS at a density of 5×10^3^ in 96-well plates. After 48 h, cells were treated with STO-609 (Cat # 1551, Tocris, Bristol, UK) or SGC-CAMKK2-1 (Cat#SML2834, Sigma, St. Louis, MO, USA) and incubated for 3 or 7 days. For siRNAs experiments, NCI-H660 cells were transfected with 100 nM final concentration siControl #1, siCAMKK2 #1 and siCAMKK2 #3 for 3 or 7 days. Then, a resazurin reduction assay, a fluorometric method, was performed according to the manufacturer’s protocols (CellTiter-Blue assay; Cat#G8080, Promega, Madison, WI, USA) to determine the presence of metabolically active cells. Fluorescent signal was measured at a wavelength of 560Ex/590Em using a Synergy H4 microplate reader (BioTek Instruments, Synergy H4, serial number 140417F).

### Proliferation assays (Hoechst)

Media was removed and cell-containing 96-well plates were frozen at −80°C. Plates were then thawed and 100 μl of MilliQ water was added to each well. After an incubation at 37°C for 1 h, plates were frozen at −80°C. At the day of the assay, cell plates were thawed and 100 ul Hoechst stain working solution (Hoechst 33342, Cat#H3570, Invitrogen, Waltham, MA, USA; 2 μl diluted in 10 ml TNE buffer, pH 7.4) was added to each well. Plates were measured on a plate reader (Synergy H4, serial number 140417F) at the following settings: excitation: 360/20.0, emission: 460/20.0, optics: top, gain: 70, light source: xenon flash, read speed: normal, delay: 100 msec, measurements/data point: 10, read height: 8 mm.

### Organoid models

Hi-MYC mice (strain number 01XK8) were obtained from the National Cancer Institute Mouse Repository at Frederick National Laboratory for Cancer Research^88^. Five Hi-MYC males were sacrificed at 6 months of age. This line was bred on the same genetic background FVB/N.

#### <u>Single cell suspension</u> (based on ^89^)

Mouse prostate tissues were digested in Advanced DMEM/F12/Collagenase II (1.5mg/ml)/Hyaluronidase (1000 u/ml) (Life Technologies, Carlsbad, CA, USA) plus 10 μM Y-27632 (Tocris, Bristol, UK) for 1 h at 37°C with 1500 rpm mixing, continuously agitated. Subsequently, after centrifuging at 150 g for 5 min at 4°C, digested cells were suspended in 1 ml TrypLE with 10 μM Y-27632 and digested for 15 min at 37°C and neutralized in DMEM/F12/FBS (0.05%). Dissociated cells were subsequently passed through 70 μm and 40 μm cell strainers (BD Biosciences, San Jose, CA, USA) to obtain a single cells suspension. Samples were resuspended in 1x PBS and sorted by Flow Cytometry (Becton-Dickinson Aria II and/or Becton-Dickinson Influx) for 4’,6-diamidino-2-phenylindole (DAPI) to enrich living cells.

#### <u>3D culture</u> (based on ^90,91^)

Next, Hi-MYC DAPI negative, prostate epithelial cells were cultured in ADMEM/F12 supplemented with B27 (Life Technologies, Carlsbad, CA, USA), 10 mM HEPES, Glutamax™ (Life Technologies, Carlsbad, CA, USA) and penicillin/streptavidin and contained following growth factors: EGF 5-50 ng/ml (Peprotech, Cranbury, NJ, USA), R-spondin1 conditioned medium or 500 ng/ml recombinant R-spondin1 (Peprotech), 100 ng/ml recombinant Noggin (Peprotech, Cranbury, NJ, USA) and the TGF-ß/Alk inhibitor A83-01 (Tocris, Bristol, UK). Dihydrotestosterone (Sigma, St. Louis, MO, USA) was added at 1 nM final concentration. Murine prostate organoids were passaged via trituration with trypsinization with TrypLE for 15 min at 37°C. Passaging was performed every week with a 1:5-1:10 ratio.

#### Organoid treatment

Following dissociation with TryplE for 15 min at 37°C, Hi-MYC organoids were seeded at 50,000 cells embedded in 20 μL of growth factor-reduced Matrigel^®^ per well in a 12-well plate. Organoids were cultured with aDMEM/F12 for high-glucose medium (17 mM glucose) or DMEM for low-glucose (5 mM glucose), in the presence of atglistatin or DMSO (vehicle). Organoids were grown for 14 days, and medium was refreshed every 3 days. At the end of treatment, organoids were imaged with a Leica Thunder microscope, and their diameter manually measured with ImageJ. Organoids were then dissociated with TryplE and viable cells determined with Vi Cell BLU analyzer (Beckman Coulter, Brea, CA, USA) for cell growth analysis.

#### Organoid lipidomics

Lipid species were extracted and analyzed by Lipometrix, at KU Leuven, Belgium. Briefly, lipid extraction was performed with 1 N HCl:CH3OH 1:8 (*v/v*), 900 μl CHCl_3_ and 200 μg/ml of the antioxidant 2,6-di-tert-butyl-4-methylphenol (BHT; Sigma-Aldrich). A mixture of deuterium-labeled lipids SPLASH® LIPIDOMIX® Mass Spec Standard (#330707, Avanti Polar Lipids, Alabaster, AL, USA) was spiked into the extract mix. The organic fraction was evaporated using a Savant Speedvac spd111v (Thermo Fisher Scientific) at room temperature and the remaining lipid pellet was stored at −20°C under argon. Just before mass spectrometry analysis, lipid pellets were reconstituted in 100% ethanol. Lipid species were analyzed by liquid chromatography electrospray ionization tandem mass spectrometry (LC-ESI/MS/MS) on a Nexera X2 UHPLC system (Shimadzu) coupled with hybrid triple quadrupole/linear ion trap mass spectrometer (6500+ QTRAP system; AB SCIEX). Chromatographic separation was performed on a XBridge amide column (150 mm × 4.6 mm, 3.5 μm; Waters) maintained at 35°C using mobile phase A [1 mM ammonium acetate in water-acetonitrile 5:95 (*v/v*)] and mobile phase B [1 mM ammonium acetate in water-acetonitrile 50:50 (*v/v*)] in the following gradient: (0-6 min: 0% B > 6% B; 6-10 min: 6% B > 25% B; 10-11 min: 25% B > 98% B; 11-13 min: 98% B > 100% B; 13-19 min: 100% B; 19-24 min: 0% B) at a flow rate of 0.7 mL/min which was increased to 1.5 mL/min from 13 minutes onwards. TGs and DGs were measured in positive ion mode with a neutral loss scan for one of the fatty acyl moieties. Lipid quantification was performed by scheduled multiple reactions monitoring, the transitions being based on the neutral losses or the typical product ions as described above. The instrument parameters were as follows: Curtain Gas = 35 psi; Collision Gas = 8 a.u. (medium); IonSpray Voltage = 5500 V and −4,500 V; Temperature = 550°C; Ion Source Gas 1 = 50 psi; Ion Source Gas 2 = 60 psi; Declustering Potential = 60 V and −80 V; Entrance Potential = 10 V and −10 V; Collision Cell Exit Potential = 15 V and −15 V. The following fatty acyl moieties were taken into account for the lipidomic analysis: 14:0, 14:1, 16:0, 16:1, 16:2, 18:0, 18:1, 18:2, 18:3, 20:0, 20:1, 20:2, 20:3, 20:4, 20:5, 22:0, 22:1, 22:2, 22:4, 22:5 and 22:6 except for TGs which considered: 16:0, 16:1, 18:0, 18:1, 18:2, 18:3, 20:3, 20:4, 20:5, 22:2, 22:3, 22:4, 22:5, 22:6.

Peak integration was performed with the MultiQuantTM software version 3.0.3. Lipid species signals were corrected for isotopic contributions (calculated with Python Molmass 2019.1.1) and were quantified based on internal standard signals and adheres to the guidelines of the Lipidomics Standards Initiative (LSI) (level 2 type quantification as defined by the LSI).

### Xenograft mouse models

This animal study was approved by and conducted under the Institutional Animal Care and Use Committee at the University of Texas MD Anderson Cancer Center (MDACC). NSG mice were obtained at the age of 6 weeks, castrated at week 7, and at 8 weeks 0.5 × 10^6^ cells in 50 μl PBS mixed with 50 μl Matrigel® were injected subcutaneously into the flank. Tumor growth was monitored using caliper measurement and tumor volume was calculated using the formula 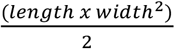. When tumors reached a length of 1.5 cm, tumors were dissected and cut into three pieces to 1) fix in formalin for IHC, 2) embed in OCT for DESI/MS, and 3) cryo-smash in liquid nitrogen for RNA/WB analysis.

### Histology and immunostaining

Tumor tissue processing, embedding, H&E and immune staining was conducted by the MD Anderson Cancer Center Science Park Research Histology, Pathology, and Imaging Core (RHPI) (Smithville, TX, USA). The following antibodies were used: cleaved caspase-3 (Cell Signaling cat #9661), p-HH3 (Ser10) (Millipore cat #06-570).

### Fatty acid treatments

100 mM oleate stocks were stored at −20°C and the day of the assay oleates were coupled to fatty acid-free BSA (Cat #A9205, Sigma, St. Louis, MO, USA) at 42°C for 20 min. BSA-coupled oleate was then mixed step wise into preheated media and sterile filtered (0.2 μM) before use. Fatty acid-containing media was prepared freshly the day of the cell treatment.

### Oil Red O staining

The Oil Red O staining protocol was modified from^92^. For the panel of prostate cancer cell lines treated with *siPNPLA2*, cells were seeded in 24-well plates to 60% confluency in RPMI 1640 supplemented with 5% charcoal-stripped FBS and then treated for 72 hours with chemical siRNAs and/or R1881. For TG assessment of ATGL KO and addback cells, cells were seeded in 24-well plates (80,000 cells/well) in RPMI 1640 no glucose supplemented with 5% or 0.5% dialyzed FBS and 5 mM glucose. After 24 h treatment with BSA-coupled oleate or BSA, cells were fixed and stained with Oil Red O as described below. RM-9 (RM-9, RM-9-sh*PNPLA2*, RM-9 shControl) cells were plated in RPMI 1640 no glucose supplemented with 0.5% dialyzed FBS and 5 mM glucose and treated atglistatin for 24 h hours and/or 600 ng/ml doxycycline, fixed with 10% formalin and stained with Oil Red O. In brief, the 6 parts of the Oil Red O stock solution (2.5 g/l Oil Red O, Cat#O0625, Sigma, St. Louis, MO, USA, diluted in isopropyl alcohol) was diluted the day of the assay with 4 parts of MilliQ water and filtered. Cells that were treated with BSA or different amounts of BSA-coupled oleate for 24 h, were fixed with 10% buffered formalin (Cat #HT501128, Sigma, St. Louis, MO, USA) for 30 min. Cells were washed with 60% isopropyl alcohol and incubated with the Oil Red O working solution for 20 min. After washing with 60% isopropyl alcohol, cells were kept in PBS and brightfield images at 20× were taken using the BioTek Cytation 5. To quantify the Oil Red O stains, images were converted to grayscale and the area of stain for each picture was determined using ImageJ and normalized to cell number per picture (manually counted).

### Triglyceride measurements

The TG content was determined using the Triglyceride-Glo™ Assay (Cat # J3160, Promega, Madison, WI, USA). In brief, C4-2 ATGL addback cell lines (WT and 404A) were seeded in 96-well plates (10,000 cells/well) and treated with BSA or BSA-coupled oleate. 24 hours after treatment, the triglyceride content was determined following the manufacturer’s recommendation. Measured TG content was normalized to cell number using a duplicate 96-well plate.

### Immunoblot analyses

Cells were washed twice with ice cold PBS and then lysed in RIPA buffer (50 mmol/L Tris (pH 8.0), 200 mmol/L NaCl, 1.5 mmol/L MgCl2, 1% NP-40, 1 mmol/L EGTA, 10% glycerol, 50 mmol/L NaF, 2 mmol/L Na_3_VO_4_, and protease inhibitors). After 0.5 – 1 h rotation at 4°C, the cell lysate was spun down and the supernatant was used for further analysis. Protein lysates were quantified via Bradford (Protein Assay Dye Reagent Concentrate Cat#5000006, Bio-Rad, Hercules, CA, USA) and 30 μg protein mixed with Laemmli buffer (62.5 mmol/L Tris-HCl (pH 6.8), 2% SDS, 25% glycerol, 5% β-mercaptoethanol, 0.01% bromophenol blue) were loaded per well on the SDS-PAGE (self-poured and adjusted to protein size (7.5-12.5%) or purchased premade (Criteron TGX precast gels, Cat#5671095, Bio-Rad, Hercules, CA, USA). The SDS-PAGE was separated at 100V or 150V (premade gels) in TRIS-Glycin buffer with SDS. Gels were transferred over night at 30V onto PVDF membranes (Bio-Rad Immun-Blot® PVDF Membrane 0.2 μM) at 4°C using a TRIS-Glycine/methanol buffer system. Membranes were then blocked at room temperature with Tris buffered saline (TBS) containing Tween 20 (TBST: 200 mM Tris. 1.5 mM NaCl) containing 5% (*w/v*) milk powder for one hour. Following the blocking the membranes were incubated with primary antibody in TBST-containing 0.5% milk powder for up to 48 h. After three, 5-minute washes with TBST, membranes were incubated with the secondary HRP conjugated antibody according to primary antibody species (Bio-Rad, Hercules, CA, USA) for up to 90 min at room temperature. After three washes with TBST, blots were developed using the Biosystems C-600 Imager (Azure, Dublin, CA, USA).

### Immunoprecipitations

Stable overexpression cell lines were created by lentiviral packaging plx304-ATGL WT or plx304-ATGL S404A in HEK293T cells and then transfecting C4-2 parental cells and selected using 10 μg/ml blasticidin. Cells were plated in 10 cm cell culture dishes at 60-70% confluency and the following day treated with R1881 or transfected with chemical siRNAs targeting both alpha subunits of AMPK or CAMKK2 (see siRNA section for sequences). After 48 h, cells were washed with ice cold PBS and lysed using RIPA lysis buffer (described above). Protein levels were detected via Bradford Assay (Bio-Rad) and 2 mg of each cell lysate was incubated with magnetic Dynabeads Protein G (Cat# 10003D, Invitrogen, Waltham, MA, USA) conjugated to either V5 (Cat #46-0705, Invitrogen, Waltham, MA, USA) antibody or mouse control IgG (Santa Cruz #sc 2025) for 2 h at room temperature. After three washes with PBS + Tween 0.02% using the Bio-Rad DynaMag2 magnet, samples were moved to a new tube, the supernatant removed via the magnet and 40 μl of Laemmli buffer (2×) was added. Samples were then boiled at 95°C for 6 min and stored at −20°C. Changes in ATGL phosphorylation at S404 were detected using the newly created pS404 ATGL antibody (Abclonal, Woburn, MA) as a primary antibody and HRP-conjugated secondary antibody. Membranes were stripped and then total levels of the protein were detected using the V5 antibody.

### 3D invasion assays

Cells were trypsinized and seeded at a density of 100,000 (C4-2) or 50,000 (RM-9) cells per well in the 5D spherical plate (Kugelmeiers, Zurich, Switzerland) and allowed to cluster in RPMI 1640 (C4-2) (Cat# 22400105, Gibco, Waltham, MA) with 10% (v/v) FBS, 1× penicillin/streptomycin or DMEM + glutamine (RM-9) in a 37°C in a humidified 5% (*v/v*) CO_2_ atmosphere^93^. After 24 h, spheroids were transferred to 1.75 mL centrifuge tubes and allowed to settle to the bottom of the tube via gravity. Media was removed and cells were resuspended in Glycosil (Advanced Biomatrix, Carlsbad, CA, USA; Cat# GS222) reconstituted in degassed water per the manufacturer’s instructions to 10 mg/mL. Peptides KGGGPQG;IWGQGK (PQ peptide) and GRGDS (RGD peptide) were purchased from Genescript USA Inc. (Piscataway, NJ, USA) and reacted as previously described^94^. RGD at 73.7 mg/mL was added to the Glycosil spheroid suspension and then mixed into solution with a pipette with a cut off end. The combined Glycosil-RGD mixture was adjusted to pH 7.8 with 1M NaOH. PQ at 11.243 mg/mL was added and then mixed as previously described^94,95^. The solution was allowed to begin crosslinking for 5 min prior pipetting in 50 μL pucks cast in PDMS molds and allowed to crosslink in the incubator. After 1 h, hydrogels were removed from the PDMS molds and cultured in 1 mL of media (C4-2: RPMI (no glucose) + 5 mM glucose + 5% FBS, RM-9: RPMI (no glucose) + 5 mM glucose + 0.5% FBS). RM-9 cells were treated with either atglistatin (40 μM) or DMSO (control) in assay media, replenished on days 3 and 5. C4-2 cells were assayed after 14 days and RM-9 cells were assayed after 5 days using live/dead assay reagents (4 μM EthD-1, 2 μM calcein AM, and 1 μg/mL Hoechst 33342 in PBS). Hydrogels laden with clusters were incubated in the live/dead reagents for 15 min prior to imaging on the Nikon A1R MP+. Images were acquired with .825 μm steps through the z-stack at 20×. The resonant scanner was used with 2× line average/integrate count and a pinhole size of 24.27 μm. Bitplane IMARIS 9.2.1 was used to analyze acquired z-stacks. C4-2 images were quantified using the spots feature (number of Hoechst positive nuclei) and surfaces (total encompassing cluster volume of calcein AM) to calculate number of cells per cluster (number of Hoechst-positive nuclei within the encompassing calcein AM volume). Invadopodia were manually counted, and final numbers were reported as invadopodia/cell. “Invadopodia” were defined as thin cellular processes extending outward from a cell cluster that were clear enough to be easily identified by eye. RM-9 z-stacks were acquired using the above acquisition settings then uploaded to IMARIS and counted manually. Cell counts include only those cells that contain a blue nucleus and green cytoplasm and were reported as the number of escaped cells per cluster. Both experiments are reported with three clusters per hydrogel done in three experimental replicates, n=9 clusters.

### Shotgun lipidomics

C4-2 ATGL KO and addback cell lines were plated to 70% confluency in 10 cm cell culture plates and treated with BSA or 75 μM BSA-coupled oleate for 24 h. Cells were harvested by washing twice with ice cold PBS, cell number and viability was measured and then diluted in PBS before snap freezing in liquid nitrogen. The frozen cells were then shipped to be analyzed by Lipotype GMBH, Dresden, Germany. Samples for shotgun lipidomics were isolated, injected, and high-resolution Orbitrap mass analysis including internal lipid-class specific standards were performed. Lipid classes were identified via the LipotypeXplorer (Lipotype GMBH, Dresden, Germany) software. Data is presented in mol%. The nomenclature was as following: glycerophosphoplids subclass, the total length of fatty acids:number of saturated fatty acids:number of hydroxyl groups (eg PC 36:2:0). If no hydroxyl groups were detected for the subclass, we labeled the lipid with the fatty acid length and number of saturated fatty acids (ex. PC 36:2).

**Table 4:** List of analyzed lipid classes.

| Lipid class | MS mode | Structural detail level |
| --- | --- | --- |
| PA | MSMS | Species |
| PC | MSMS | Species |
| PE | MSMS | Species |
| PG | MSMS | Species |
| PI | MSMS | Species |
| PS | MSMS | Species |
| SM | MS | Species |
| CE | MSMS | Species |

**Table 5:** List of equipment and software used for shotgun lipidomics.

| Analysis step | Equipment or software |
| --- | --- |
| Extraction | Hamilton Robotics STARlet |
| Sample infusion | Advion Triversa Nanomate |
| Mass spectrometry | Thermo Scientific Q-Exactive |
| Lipid Identification | LipotypeXplorer |
| Data processing | Lipotype LIMS and LipotypeZoom |

### DESI-MS imaging

Xenograft tumor tissues were stored at −80°C prior to analysis. Frozen tissues were sectioned at 12 μm thickness, thaw-mounted onto glass slides, and immediately analyzed using a Q Exactive HF Orbitrap mass spectrometer (Thermo Fisher Scientific) coupled to a 2D OmniSpray stage (Prosolia Inc.) and equipped with a home-built DESI sprayer. DESI-MS imaging was performed on consecutive sections from each sample in both the positive and negative ionization modes using a mass resolving power of 70,000 and a spatial resolution of 200 μm. A histologically compatible solvent system comprised of 98% methanol was used as the DESI spray solvent at a flow rate of 5.0 μL/min. Ion images were assembled and visualized using Firefly (Prosolia, Inc.) and BioMap (Novartis) software. For ion identification, tandem mass spectrometry experiments were performed using HCD and high mass accuracy measurements. After DESI-MS imaging, the analyzed tissue sections were stained with H&E and annotated to delineate histopathological regions of interest. Mass spectral data was extracted from pixels corresponding spatially to tumor regions within the DESI-MS images using MSiReader software. Per-pixel ion intensities for selected metabolite and lipid species were normalized by the total ion count and averaged across each sample for subsequent statistical analyses.

### Seahorse analysis

Agilent Seahorse 96-well cell assay plates were coated with Poly-lysine (0.1% in H_2_O, Cat# P8920, Sigma, St. Louis, MO, USA) for 5 min, followed by two washes with PBS and dried. After sterilization via UV, cells were seeded (C4-2: 20,000 cells/well, RM-9: 10,000 cells/well) in RPMI 1640 no glucose media supplemented with dialyzed FBS (C4-2: 5%, RM-9: 0.5%) and glucose (RM-9, C4-2: 5 mM). 24 h after seeding, cells were treated as indicated and after 24 h of treatment Seahorse analysis were performed using the Xfe96 analyzer. The Glycolysis Stress assay was performed according to manufacturer’s manual using the following final concentrations per well: Glycolysis Stress assay RM-9 (glucose: 10 mM, oligomycin: 2 μM, 2DG: 50 mM), Glycolysis Stress assay C4-2 (glucose: 10 mM, oligomycin: 1 μM, 2DG: 50 mM). Results were analyzed using WAVE 2.6.3 and ECAR/OCR was normalized to relative cell number using Hoechst stain as described in proliferation section. Results were graphed using Graph Pad 9.3.1.

### Antibodies

**Table 6:** List of antibodies used for ATGL study.

| Antibody | Catalog number | Used for |
| --- | --- | --- |
| pATGL S406 | Abcam ab135093 | WB |
| V5 | Invitrogen #46-0705 | WB, IP |
| pAMPK (T172) | Cell signaling #2535 | WB |
| pAMPK Substrate Motif [LXRXX(pS/pT) MultiMab | Cell signaling #5759 | WB |
| AMPK | Cell signaling # 2532 | WB |
| CAMKK2 | BD biosciences Cat# 610544 | WB |
| ATGL | Cell signaling # 2138 | WB |
| pATGL S404 (developed for this study) | ABclonal (rabbit polyclonal) | WB |
| GAPDH | Sigma # G9545 | WB |
| Cleaved Caspase-3 | Cell Signaling Cat# 9661 | IHC |
| p-HH3 (Ser10) | Millipore Cat# 06-570 | IHC |

### Primers

**Table 7:**
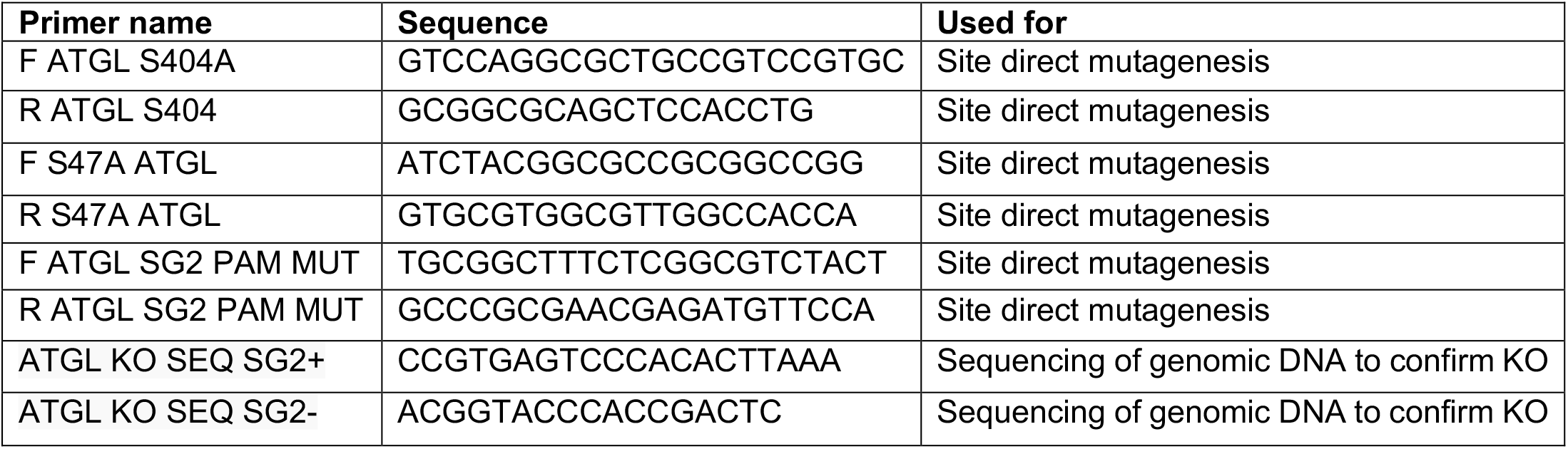
Primers used for the ATGL study.

### Plasmids

pLenti PGK Neo DEST (w531-1) was a gift from Eric Campeau & Paul Kaufman (plasmid # 19067, Addgene, Watertown, MA, USA). pLX304 was a gift from David Root (plasmid # 25890, Addgene, Watertown, MA, USA). lentiCRISPRv2 was a gift from Junjie Chen (UT MD Anderson Cancer Center).

**Table 8:**
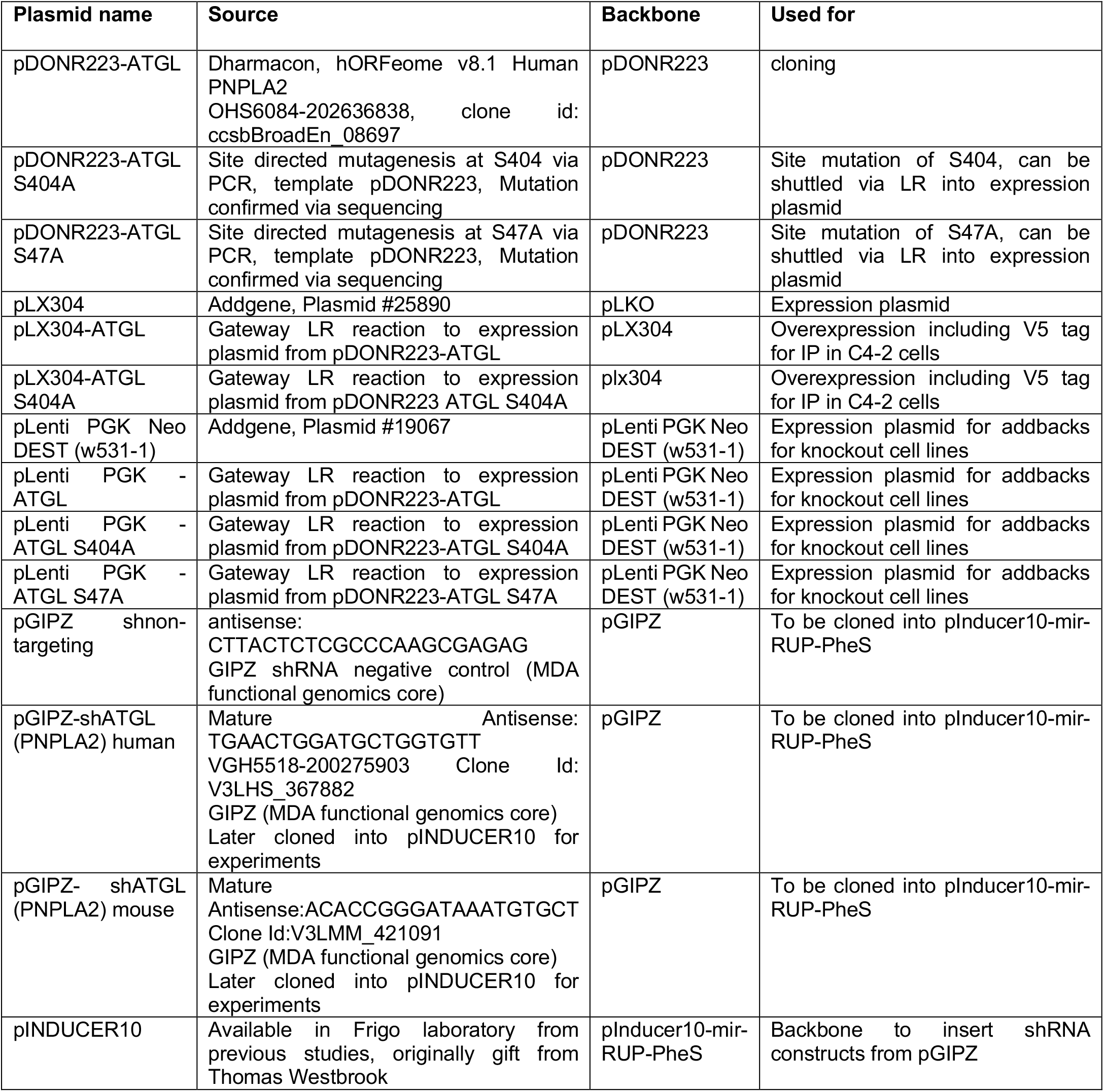
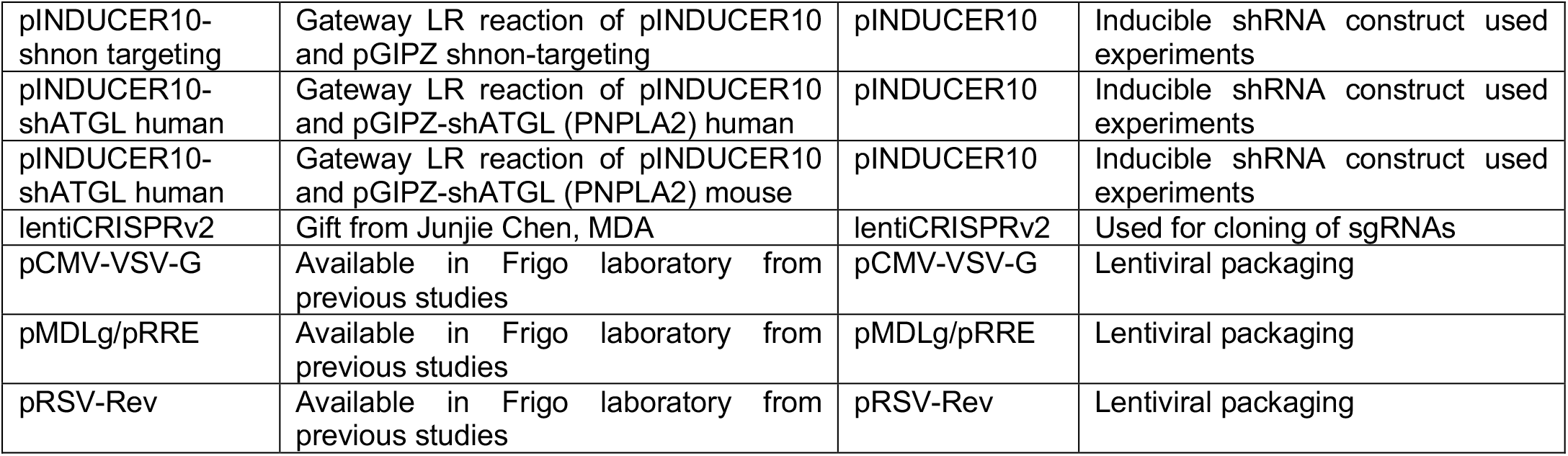
Plasmids used/created for this study.

### Statistical analysis

Statistical analyses were performed using Prism Graph Pad Version 9.3.1. One-way ANOVAs with Dunnett or Tukey (DESI-MS only) post-hoc as well as *t* tests (unpaired, two tailed) were performed as indicated for each figure.

### Synergy model calculations

CDI was calculated by the following formula: CDI = AB (AxB), where AB is the combination of drug treatment, A and B each of the drugs by itself ^96^. The CDI of x<1 means synergetic, x=1 means additive and x>1 means antagonistic. For the synergy model using cell viability, RM-9 cells were plated in 96-well plates (500 cells/well) in RPMI 1640 supplemented with 0.5% dialyzed FBS and 5 mM glucose. Cells were treated 24 h after seeding with atglistatin or AZ-PFKFB3-26 in various combination treatments. Cells were retreated every 48 h and cells were harvested after five days of treatment and cell viabilities were determined via CellTiter-Blue® assays. Synergy was determined using SynergyFinder 2.0^97^. Settings were applied as follows. Readout: viability. Detect outliers: Yes, Baseline correction: Yes, Curve fitting: LL4, Calculate synergy: HSA.

## References

1. Siegel, R. L., Miller, K. D., Fuchs, H. E. & Jemal, A. Cancer statistics, 2022. CA Cancer J. Clin. 72, 7–33 (2022).

2. Awad, D., Pulliam, T. L., Lin, C., Wilkenfeld, S. R. & Frigo, D. E. Delineation of the androgen-regulated signaling pathways in prostate cancer facilitates the development of novel therapeutic approaches. Current Opinion in Pharmacology (2018) doi:10.1016/j.coph.2018.03.002.

3. Litwin, M. S. & Tan, H.-J. The Diagnosis and Treatment of Prostate Cancer. JAMA (2017) doi:10.1001/jama.2017.7248.

4. Crona, D. & Whang, Y. Androgen Receptor-Dependent and -Independent Mechanisms Involved in Prostate Cancer Therapy Resistance. Cancers (2017) doi:10.3390/cancers9060067.

5. Frigo, D. E. et al. CaM kinase kinase β-mediated activation of the growth regulatory kinase AMPK is required for androgen-dependent migration of prostate cancer cells. Cancer Res. 71, 528–537 (2011).

6. Garcia, D. & Shaw, R. J. AMPK: Mechanisms of Cellular Energy Sensing and Restoration of Metabolic Balance. Molecular Cell (2017) doi:10.1016/j.molcel.2017.05.032.

7. Karacosta, L. G., Foster, B. A., Azabdaftari, G., Feliciano, D. M. & Edelman, A. M. A regulatory feedback loop between Ca2+/calmodulin-dependent protein kinase kinase 2 (CaMKK2) and the androgen receptor in prostate cancer progression. J. Biol. Chem. (2012) doi:10.1074/jbc.M112.370783.

8. Massie, C. E. et al. The androgen receptor fuels prostate cancer by regulating central metabolism and biosynthesis. EMBO J. (2011) doi:10.1038/emboj.2011.158.

9. Dadwal, U. C., Chang, E. S. & Sankar, U. Androgen receptor-CaMKK2 axis in prostate cancer and bone microenvironment. Frontiers in Endocrinology (2018) doi:10.3389/fendo.2018.00335.

10. Hawley, S. A. et al. Complexes between the LKB1 tumor suppressor, STRAD alpha/beta and MO25 alpha/beta are upstream kinases in the AMP-activated protein kinase cascade. J. Biol. 2, 28 (2003).

11. Woods, A. et al. LKB1 is the upstream kinase in the AMP-activated protein kinase cascade. Curr. Biol. 13, 2004–2008 (2003).

12. Shaw, R. J. et al. The tumor suppressor LKB1 kinase directly activates AMP-activated kinase and regulates apoptosis in response to energy stress. Proc. Natl. Acad. Sci. U. S. A. 101, 3329–3335 (2004).

13. Song, X. et al. AMP-activated protein kinase is required for cell survival and growth in HeLa-S3 cells in vivo. IUBMB Life 66, 415–423 (2014).

14. Ríos, M. et al. AMPK activation by oncogenesis is required to maintain cancer cell proliferation in astrocytic tumors. Cancer Res. 73, 2628–2638 (2013).

15. Mendoza, E. E. et al. Control of glycolytic flux by AMP-activated protein kinase in tumor cells adapted to low pH. Transl. Oncol. 5, 208–216 (2012).

16. Chhipa, R. R., Wu, Y. & Ip, C. AMPK-mediated autophagy is a survival mechanism in androgen-dependent prostate cancer cells subjected to androgen deprivation and hypoxia. Cell. Signal. 23, 1466–1472 (2011).

17. Jeon, S.-M., Chandel, N. S. & Hay, N. AMPK regulates NADPH homeostasis to promote tumour cell survival during energy stress. Nature 485, 661–665 (2012).

18. Tennakoon, J. B. et al. Androgens regulate prostate cancer cell growth via an AMPK-PGC-1α-mediated metabolic switch. Oncogene 33, 5251–5261 (2014).

19. Raj Chhipa, R. et al. AMP kinase promotes glioblastoma bioenergetics and tumour growth. Nat. Cell Biol. (2018) doi:10.1038/s41556-018-0126-z.

20. Lin, C. et al. Inhibition of CAMKK2 impairs autophagy and castration-resistant prostate cancer via suppression of AMPK-ULK1 signaling. Oncogene 40, 1690–1705 (2021).

21. Zimmermann, R. et al. Fat mobilization in adipose tissue is promoted by adipose triglyceride lipase. Science 306, 1383–1386 (2004).

22. Villena, J. A., Roy, S., Sarkadi-Nagy, E., Kim, K. H. & Hei, S. S. Desnutrin, an adipocyte gene encoding a novel patatin domain-containing protein, is induced by fasting and glucocorticoids: Ectopic expression of desnutrin increases triglyceride hydrolysis. J. Biol. Chem. (2004) doi:10.1074/jbc.M403855200.

23. Jenkins, C. M. et al. Identification, cloning, expression, and purification of three novel human calcium-independent phospholipase A2 family members possessing triacylglycerol lipase and acylglycerol transacylase activities. J. Biol. Chem. 279, 48968–48975 (2004).

24. Butler, L. M., Centenera, M. M. & Swinnen, J. V. Androgen control of lipid metabolism in prostate cancer: novel insights and future applications. Endocr. Relat. Cancer (2016) doi:10.1530/erc-15-0556.

25. Swinnen, J. V. Androgens markedly stimulate the accumulation of neutral lipids in the human prostatic adenocarcinoma cell line LNCaP. Endocrinology 137, 4468–4474 (1996).

26. Gorlov, I. P. et al. Candidate pathways and genes for prostate cancer: a meta-analysis of gene expression data. BMC Med. Genomics 2, 48 (2009).

27. Sharma, N. L. et al. The androgen receptor induces a distinct transcriptional program in castration-resistant prostate cancer in man. Cancer Cell 23, 35–47 (2013).

28. Pinthus, J. H. et al. Androgen-dependent regulation of medium and long chain fatty acids uptake in prostate cancer. Prostate (2007) doi:10.1002/pros.20609.

29. Liu, Y., Zuckier, L. S. & Ghesani, N. V. Dominant uptake of fatty acid over glucose by prostate cells: a potential new diagnostic and therapeutic approach. Anticancer Res. 30, 369–374 (2010).

30. Mitra, R., Chao, O., Urasaki, Y., Goodman, O. B. & Le, T. T. Detection of lipid-rich prostate circulating tumour cells with coherent anti-Stokes Raman scattering microscopy. BMC Cancer 12, 540 (2012).

31. Sacca, P. A. et al. Human periprostatic adipose tissue: Secretome from patients with prostate cancer or benign prostate hyperplasia. Cancer Genomics Proteomics 16, 29–58 (2019).

32. Taylor, R. A., Lo, J., Ascui, N. & Watt, M. J. Linking obesogenic dysregulation to prostate cancer progression. Endocr. Connect. 4, R68–80 (2015).

33. Xie, H. et al. Adipose triglyceride lipase activity regulates cancer cell proliferation via AMP-kinase and mTOR signaling. Biochim. Biophys. Acta Mol. Cell Biol. Lipids 1865, 158737 (2020).

34. Al-Zoughbi, W. et al. Loss of adipose triglyceride lipase is associated with human cancer and induces mouse pulmonary neoplasia. Oncotarget 7, 33832–33840 (2016).

35. Mitra, R., Le, T. T., Gorjala, P. & Goodman, O. B. Positive regulation of prostate cancer cell growth by lipid droplet forming and processing enzymes DGAT1 and ABHD5. BMC Cancer (2017) doi:10.1186/s12885-017-3589-6.

36. Chen, G. et al. Loss of ABHD5 promotes the aggressiveness of prostate cancer cells. Sci. Rep. (2017) doi:10.1038/s41598-017-13398-w.

37. Drake, J. M. et al. Phosphoproteome Integration Reveals Patient-Specific Networks in Prostate Cancer. Cell 166, 1041–1054 (2016).

38. Schaffer, B. E. et al. Identification of AMPK Phosphorylation Sites Reveals a Network of Proteins Involved in Cell Invasion and Facilitates Large-Scale Substrate Prediction. Cell Metab. (2015) doi:10.1016/j.cmet.2015.09.009.

39. Smirnova, E. et al. ATGL has a key role in lipid droplet/adiposome degradation in mammalian cells. EMBO Rep. (2006) doi:10.1038/sj.embor.7400559.

40. Kim, S.-J. et al. AMPK Phosphorylates Desnutrin/ATGL and Hormone-Sensitive Lipase To Regulate Lipolysis and Fatty Acid Oxidation within Adipose Tissue. Mol. Cell. Biol. (2016) doi:10.1128/MCB.00244-16.

41. Ahmadian, M. et al. Desnutrin/ATGL is regulated by AMPK and is required for a brown adipose phenotype. Cell Metab. (2011) doi:10.1016/j.cmet.2011.05.002.

42. Mason, R. R., Meex, R. C. R., Lee-Young, R., Canny, B. J. & Watt, M. J. Phosphorylation of adipose triglyceride lipase Ser(404) is not related to 5’-AMPK activation during moderate-intensity exercise in humans. Am. J. Physiol. Endocrinol. Metab. 303, E534–41 (2012).

43. White, M. A. et al. GLUT12 promotes prostate cancer cell growth and is regulated by androgens and CaMKK2 signaling. Endocr. Relat. Cancer 25, (2018).

44. Schweiger, M., Lass, A., Zimmermann, R., Eichmann, T. O. & Zechner, R. Neutral lipid storage disease: genetic disorders caused by mutations in adipose triglyceride lipase/PNPLA2 or CGI-58/ABHD5. Am. J. Physiol. Endocrinol. Metab. 297, E289–96 (2009).

45. Lord, C. C. & Brown, J. M. Distinct roles for alpha-beta hydrolase domain 5 (ABHD5/CGI-58) and adipose triglyceride lipase (ATGL/PNPLA2) in lipid metabolism and signaling. Adipocyte 1, 123–131 (2012).

46. Vande Voorde, J. et al. Improving the metabolic fidelity of cancer models with a physiological cell culture medium. Sci. Adv. 5, eaau7314 (2019).

47. Fu, H. et al. MicroRNA-224 and its target CAMKK2 synergistically influence tumor progression and patient prognosis in prostate cancer. Tumour Biol. 36, 1983–1991 (2015).

48. Butler, L. M. et al. Lipidomic Profiling of Clinical Prostate Cancer Reveals Targetable Alterations in Membrane Lipid Composition. Cancer Res. 81, 4981–4993 (2021).

49. Chen, M. et al. An aberrant SREBP-dependent lipogenic program promotes metastatic prostate cancer. Nat. Genet. 50, 206–218 (2018).

50. Lu, X. et al. HOXB13 suppresses de novo lipogenesis through HDAC3-mediated epigenetic reprogramming in prostate cancer. Nat. Genet. 54, 670–683 (2022).

51. Hishikawa, D., Hashidate, T., Shimizu, T. & Shindou, H. Diversity and function of membrane glycerophospholipids generated by the remodeling pathway in mammalian cells. J. Lipid Res. 55, 799–807 (2014).

52. Fujimoto, T. & Parton, R. G. Not just fat: the structure and function of the lipid droplet. Cold Spring Harb. Perspect. Biol. 3, a004838–a004838 (2011).

53. Mayer, N. et al. Development of small-molecule inhibitors targeting adipose triglyceride lipase. Nat. Chem. Biol. 9, 785–787 (2013).

54. Iglesias, J. et al. Simplified assays of lipolysis enzymes for drug discovery and specificity assessment of known inhibitors. J. Lipid Res. 57, 131–141 (2016).

55. Schweiger, M. et al. Pharmacological inhibition of adipose triglyceride lipase corrects high-fat diet-induced insulin resistance and hepatosteatosis in mice. Nat. Commun. 8, 14859 (2017).

56. Honeder, S. et al. Adipose Triglyceride Lipase Loss Promotes a Metabolic Switch in A549 Non-Small Cell Lung Cancer Cell Spheroids. Mol. Cell. Proteomics 100095 (2021).

57. Beg, M., Zhang, W., McCourt, A. C. & Enerbäck, S. ATGL activity regulates GLUT1-mediated glucose uptake and lactate production via TXNIP stability in adipocytes. J. Biol. Chem. 296, 100332 (2021).

58. Di Leo, L. et al. Forcing ATGL expression in hepatocarcinoma cells imposes glycolytic rewiring through PPAR-α/p300-mediated acetylation of p53. Oncogene 38, 1860–1875 (2019).

59. Boyd, S. et al. Structure-based design of potent and selective inhibitors of the metabolic kinase PFKFB3. J. Med. Chem. (2015) doi:10.1021/acs.jmedchem.5b00352.

60. St-Gallay, S. A. et al. A High-Throughput Screening Triage Workflow to Authenticate a Novel Series of PFKFB3 Inhibitors. SLAS Discovery (2018) doi:10.1177/2472555217732289.

61. Gindlhuber, J. et al. Deletion of Adipose Triglyceride Lipase Links Triacylglycerol Accumulation to a More-Aggressive Phenotype in A549 Lung Carcinoma Cells. J. Proteome Res. (2018) doi:10.1021/acs.jproteome.7b00782.

62. Zagani, R., El-Assaad, W., Gamache, I. & Teodoro, J. G. Inhibition of adipose triglyceride lipase (ATGL) by the putative tumor suppressor G0S2 or a small molecule inhibitor attenuates the growth of cancer cells. Oncotarget 6, (2015).

63. Crawford, S. E. et al. Adipose Triglyceride Lipase (ATGL) Expression Is Associated with Adiposity and Tumor Stromal Proliferation in Patients with Pancreatic Ductal Adenocarcinoma. Anticancer Res. (2017) doi:10.21873/anticanres.11366.

64. Liu, X. et al. Long non-coding RNA NEAT1-modulated abnormal lipolysis via ATGL drives hepatocellular carcinoma proliferation. Mol. Cancer 17, (2018).

65. Chen, G., Zhou, G., Lotvola, A., Granneman, J. G. & Wang, J. ABHD5 suppresses cancer cell anabolism through lipolysis-dependent activation of the AMPK/mTORC1 pathway. J. Biol. Chem. 296, 100104 (2021).

66. Lass, A. et al. Adipose triglyceride lipase-mediated lipolysis of cellular fat stores is activated by CGI-58 and defective in Chanarin-Dorfman Syndrome. Cell Metab. 3, 309–319 (2006).

67. Nardi, F. et al. DGAT1 Inhibitor Suppresses Prostate Tumor Growth and Migration by Regulating Intracellular Lipids and Non-Centrosomal MTOC Protein GM130. Sci. Rep. 9, (2019).

68. Notari, L. et al. Identification of a lipase-linked cell membrane receptor for pigment epithelium-derived factor. J. Biol. Chem. 281, 38022–38037 (2006).

69. Sathyanarayan, A., Mashek, M. T. & Mashek, D. G. ATGL Promotes Autophagy/Lipophagy via SIRT1 to Control Hepatic Lipid Droplet Catabolism. Cell Rep. 19, 1–9 (2017).

70. Patel, R. et al. ATGL is a biosynthetic enzyme for fatty acid esters of hydroxy fatty acids. Nature 606, 968–975 (2022).

71. Mayer, N. et al. Structure-activity relationship studies for the development of inhibitors of murine adipose triglyceride lipase (ATGL). Bioorganic and Medicinal Chemistry 28, 115610 (2020).

72. Jumper, J. et al. Highly accurate protein structure prediction with AlphaFold. Nature 596, 583–589 (2021).

73. Schweiger, M. et al. The C-terminal region of human adipose triglyceride lipase affects enzyme activity and lipid droplet binding. J. Biol. Chem. 283, 17211–17220 (2008).

74. Xie, X. et al. Identification of a novel phosphorylation site in adipose triglyceride lipase as a regulator of lipid droplet localization. Am. J. Physiol. Endocrinol. Metab. 306, E1449–59 (2014).

75. Pagnon, J. et al. Identification and functional characterization of protein kinase A phosphorylation sites in the major lipolytic protein, adipose triglyceride lipase. Endocrinology 153, 4278–4289 (2012).

76. Eidelman, E., Twum-Ampofo, J., Ansari, J. & Siddiqui, M. M. The Metabolic Phenotype of Prostate Cancer. Front. Oncol. (2017) doi:10.3389/fonc.2017.00131.

77. Haemmerle, G. et al. Defective lipolysis and altered energy metabolism in mice lacking adipose triglyceride lipase. Science 312, 734–737 (2006).

78. Schreiber, R. et al. Cold-Induced Thermogenesis Depends on ATGL-Mediated Lipolysis in Cardiac Muscle, but Not Brown Adipose Tissue. Cell Metab. 26, 753–763.e7 (2017).

79. Schoiswohl, G. et al. Adipose triglyceride lipase plays a key role in the supply of the working muscle with fatty acids. J. Lipid Res. 51, 490–499 (2010).

80. Haemmerle, G. et al. ATGL-mediated fat catabolism regulates cardiac mitochondrial function via PPAR-α and PGC-1. Nat. Med. 17, 1076–1085 (2011).

81. Schreiber, R. et al. Hypophagia and metabolic adaptations in mice with defective ATGL-mediated lipolysis cause resistance to HFD-induced obesity. Proc. Natl. Acad. Sci. U. S. A. 112, 13850–13855 (2015).

82. Grabner, G. F. et al. Small-molecule inhibitors targeting lipolysis in human adipocytes. J. Am. Chem. Soc. 144, 6237–6250 (2022).

83. Freedland, S. J. & Abrahamsson, P.-A. Androgen deprivation therapy and side effects: are GnRH antagonists safer? Asian J. Androl. 23, 3–10 (2021).

84. Higano, C. S. Update on cardiovascular and metabolic risk profiles of hormonal agents used in managing advanced prostate cancer. Urol. Oncol. 38, 912–917 (2020).

85. Aparicio, A. et al. Neuroendocrine prostate cancer xenografts with large-cell and small-cell features derived from a single patient’s tumor: morphological, immunohistochemical, and gene expression profiles. Prostate 71, 846–856 (2011).

86. Baley, P. A., Yoshida, K., Qian, W., Sehgal, I. & Thompson, T. C. Progression to androgen insensitivity in a novel in vitro mouse model for prostate cancer. J. Steroid Biochem. Mol. Biol. 52, 403–413 (1995).

87. Cong, L. & Zhang, F. Genome engineering using CRISPR-Cas9 system. Methods Mol. Biol. 1239, 197–217 (2015).

88. Ellwood-Yen, K. et al. Myc-driven murine prostate cancer shares molecular features with human prostate tumors. Cancer Cell 4, 223–238 (2003).

89. Drost, J. et al. Organoid culture systems for prostate epithelial and cancer tissue. Nat. Protoc. 11, 347–358 (2016).

90. Karthaus, W. R. et al. Identification of multipotent luminal progenitor cells in human prostate organoid cultures. Cell 159, 163–175 (2014).

91. Pakula, H. et al. Protocols for studies on TMPRSS2/ERG in prostate cancer. Methods Mol. Biol. 1786, 131–151 (2018).

92. Sikkeland, J., Lindstad, T. & Saatcioglu, F. Analysis of androgen-induced increase in lipid accumulation in prostate cancer cells. Methods Mol. Biol. 776, 371–382 (2011).

93. Tellman, T. V., Cruz, L. A., Grindel, B. J. & Farach-Carson, M. C. Cleavage of the perlecan-Semaphorin 3A-Plexin A1-Neuropilin-1 (PSPN) Complex by matrix metalloproteinase 7/matrilysin triggers prostate cancer cell dyscohesion and migration. Int. J. Mol. Sci. 22, 3218 (2021).

94. Hubka, K. M., Carson, D. D., Harrington, D. A. & Farach-Carson, M. C. Perlecan domain I gradients establish stable biomimetic heparin binding growth factor gradients for cell migration in hydrogels. Acta Biomater. 97, 385–398 (2019).

95. Fong, E. L. S. et al. A 3D in vitro model of patient-derived prostate cancer xenograft for controlled interrogation of in vivo tumor-stromal interactions. Biomaterials 77, 164–172 (2016).

96. Rice, M. A. et al. SU086, an inhibitor of HSP90, impairs glycolysis and represents a treatment strategy for advanced prostate cancer. Cell Rep Med 3, 100502 (2022).

97. Ianevski, A., Giri, A. K. & Aittokallio, T. SynergyFinder 2.0: visual analytics of multi-drug combination synergies. Nucleic Acids Res. 48, W488–W493 (2020).

