## Supplemental Figures for "Adipose triglyceride lipase is regulated by CAMKK2-AMPK signaling and drives advanced prostate cancer"

Supplemental Figure S1

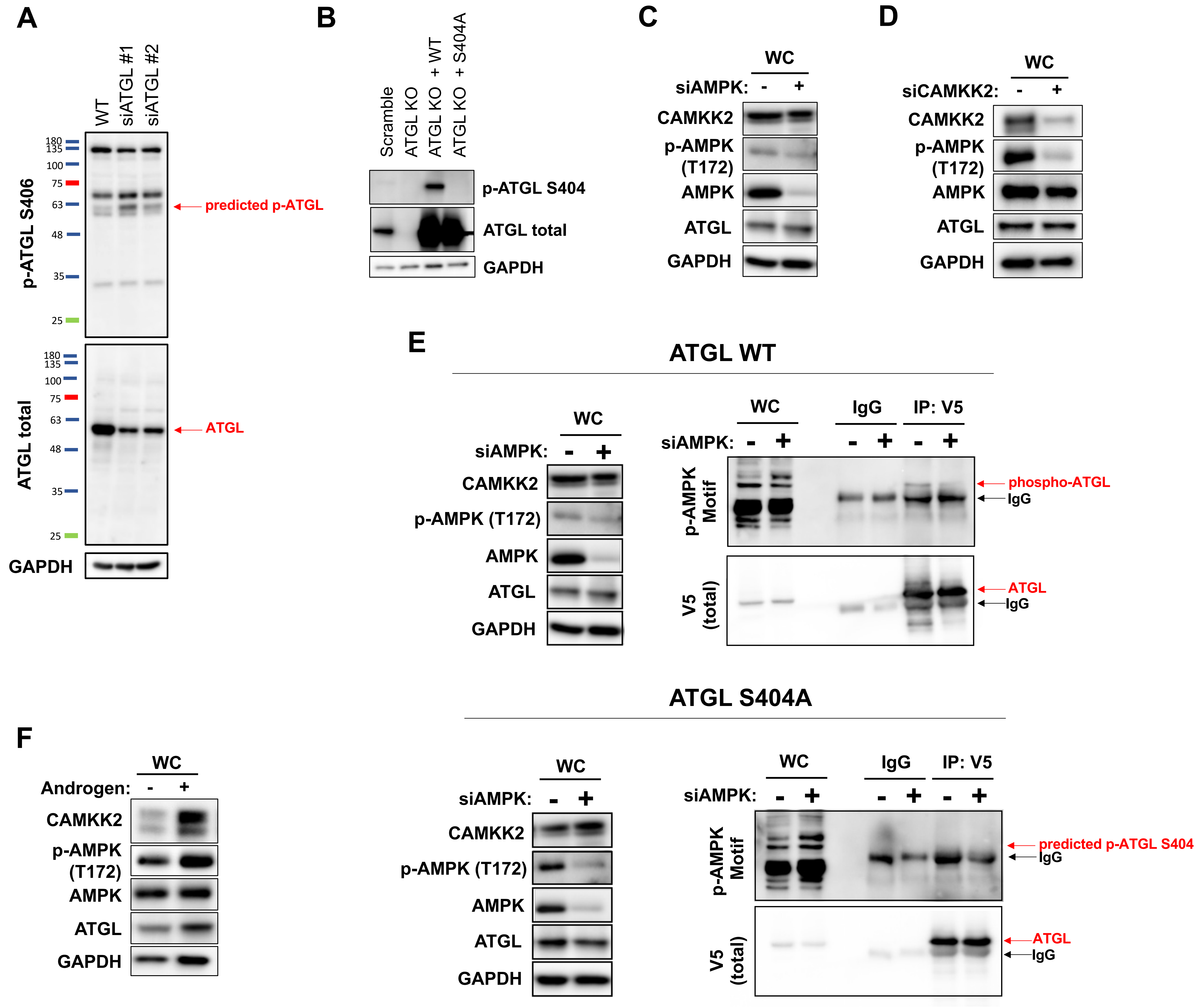


**Supplemental Figure S1: Evaluation of existing and new anti-p-ATGL antibodies, validation of siRNAs and androgen-mediated effects on CAMKK2-AMPK signaling, and orthogonal confirmation of ATGL(S404) as a downstream target of AMPK in prostate cancer cells.** (A) LNCaP cells were treated with siRNAs targeting *PNPLA2* (encoding ATGL) to test murine-specific p-ATGL antibody (p-ATGL S406, Abcam ab135093). (B) Validation of the new human-recognizing p-ATGL S404 antibody using C4-2 Scramble, ATGL KO and addback cell derivates. (C-D) Western blot controls for Figures 1D-E, respectively, from C4-2 cells transfected with siRNAs targeting scramble control (-), the AMPK $\alpha$ catalytic subunits (*PRKAA1/2*) or *CAMKK2* (siAMPK or siCAMKK2) for 72h. WC, whole cell lysates. (E) Immunoprecipitated ATGL-V5 wildtype (WT) overexpressed in C4-2 cells demonstrated a decrease in phosphorylation in the presence of chemicals siRNA targeting AMPK ($\alpha$ catalytic subunits *PRKAA1/2*) when using a broad p-AMPK substrate motif antibody (LXRXXS). The p-ATGL signal was specific to S404, since no signal could be detected when ATGL (S404A)-V5 was used for the immunoprecipitation. (F) Western blot control for Figure 1F from C4-2 cells treated with vehicle or synthetic androgen (R1881) for 72h.

Supplemental Figure S2

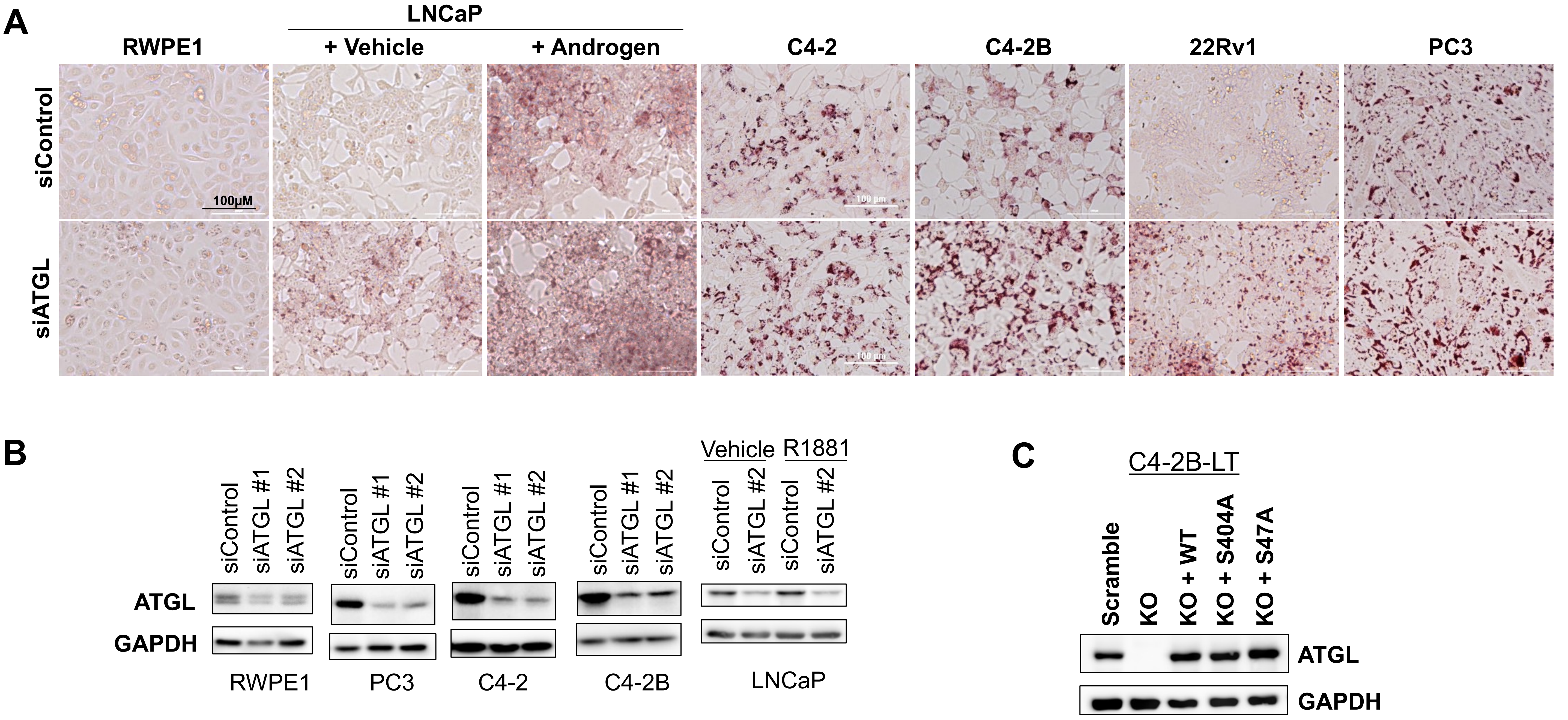


**Supplemental Figure S2: ATGL controls access to intracellular lipids in multiple prostate cancer cell models.** (A) Panel of cell models of non-transformed prostate epithelial cells (RWPE1), hormone-sensitive, AR+ prostate cancer (LNCAP), AR+ CRPC (C4-2, C4-2B, 22Rv1) and AR- CRPC (PC3) show strong accumulation of triglycerides in prostate cancer cells only when treated with chemical siRNA targeting *PNPLA2* (encoding ATGL) as assessed by Oil Red O staining. (B) Immunoblot validation of chemical siRNAs used for Supplemental Fig. S2A. SiATGL #2 was used for Supplemental Fig. S2A. (C) Western blot control for C4-2B-LT ATGL KO and add-back cell models. Images are representative results of at least three independent experiments.

Supplemental Figure S3

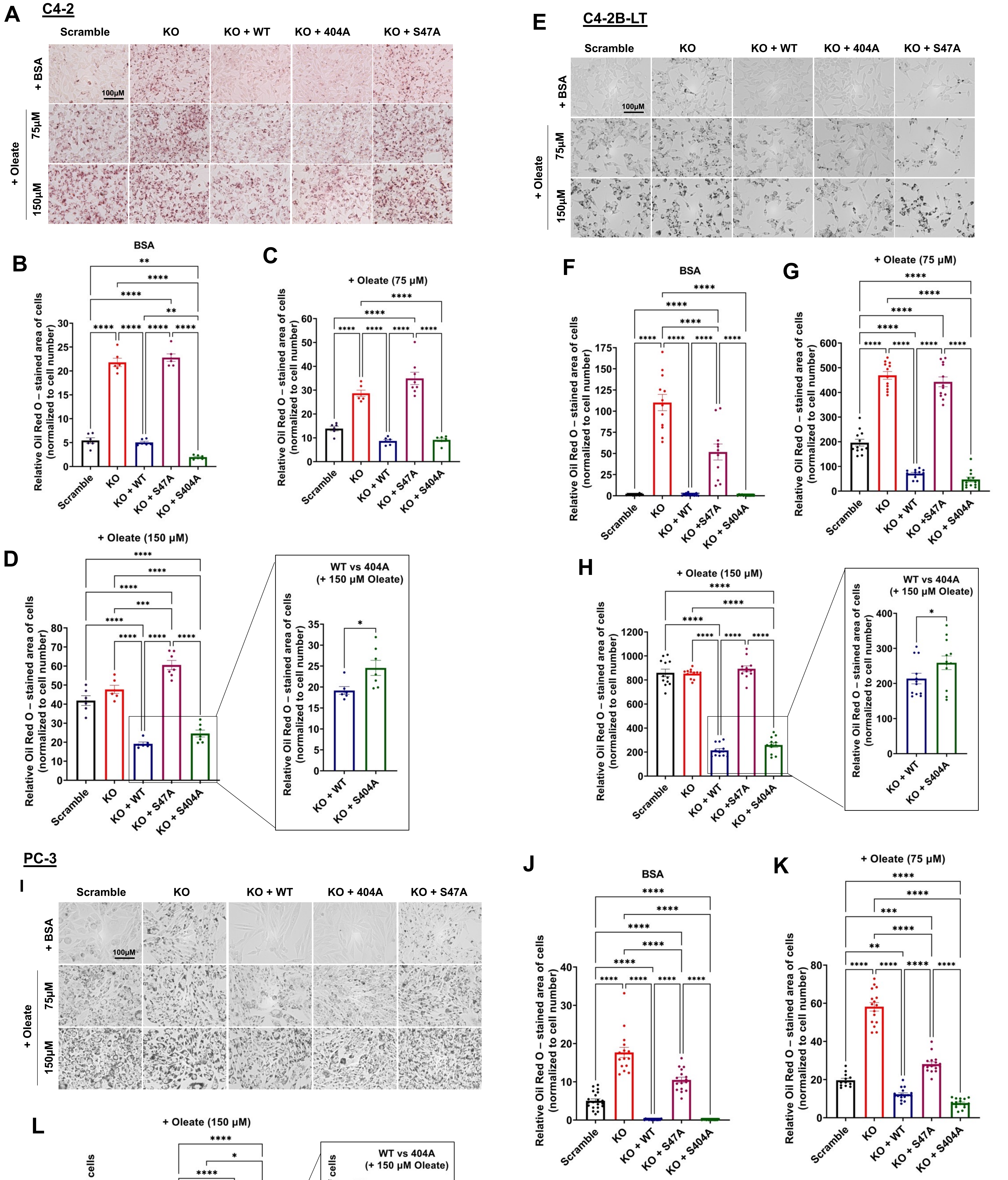


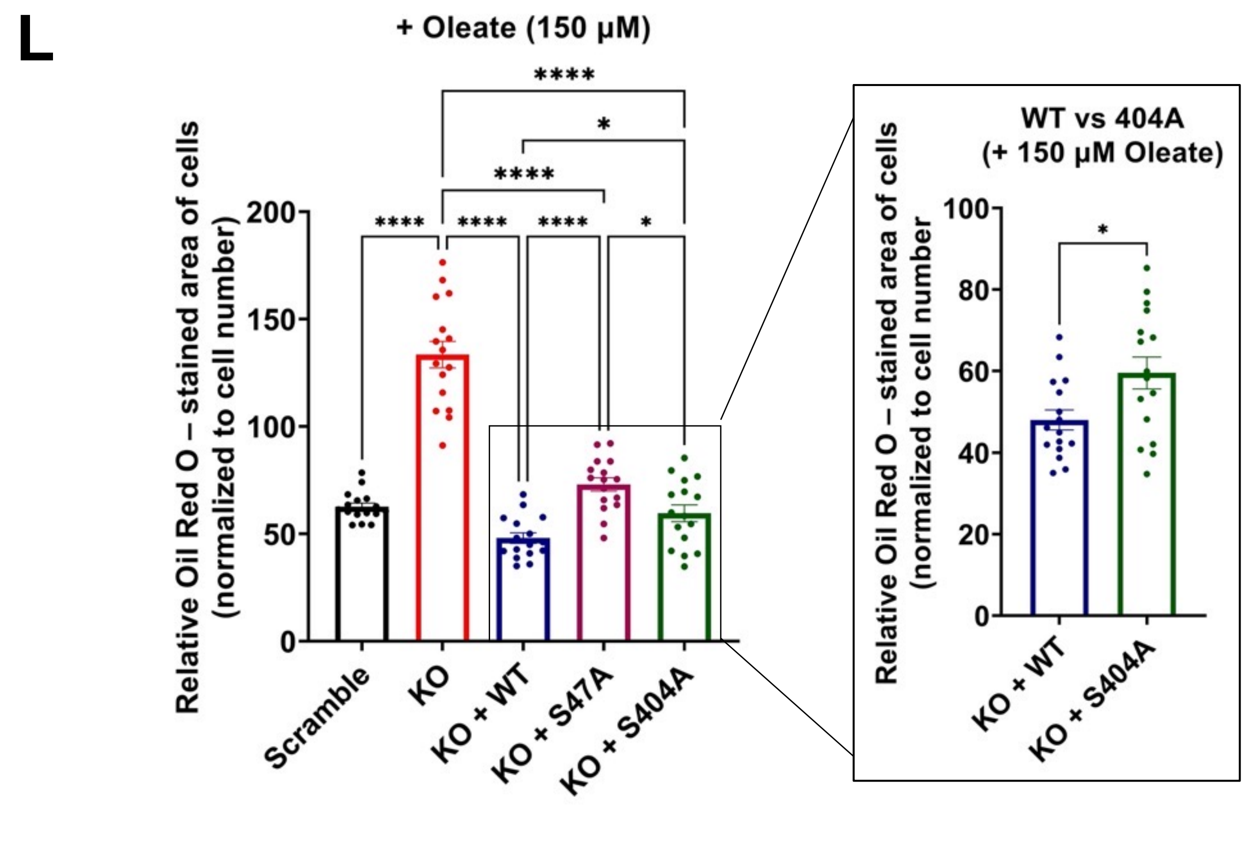


**Supplemental Figure S3: ATGL S404 fine-tunes access to intracellular lipids in multiple prostate cancer cell models in the presence of high fatty acid levels.** Representative images and quantification of Oil Red O stains performed with (A-D) C4-2, (E-H; Western blot control shown in Supplemental Fig. S2C) C4-2B-LT and (I-L) PC-3 cells. These data supplement the Oil Red O staining reported in Figure 1. Images and graphs are representative results of at least three independent experiments. One-way ANOVA or unpaired two-tailed t-test. *$P < 0.05$, **$P < 0.01$, ***$P < 0.001$, ****$P < 0.0001$, ns = no significance.

Supplemental Figure S4

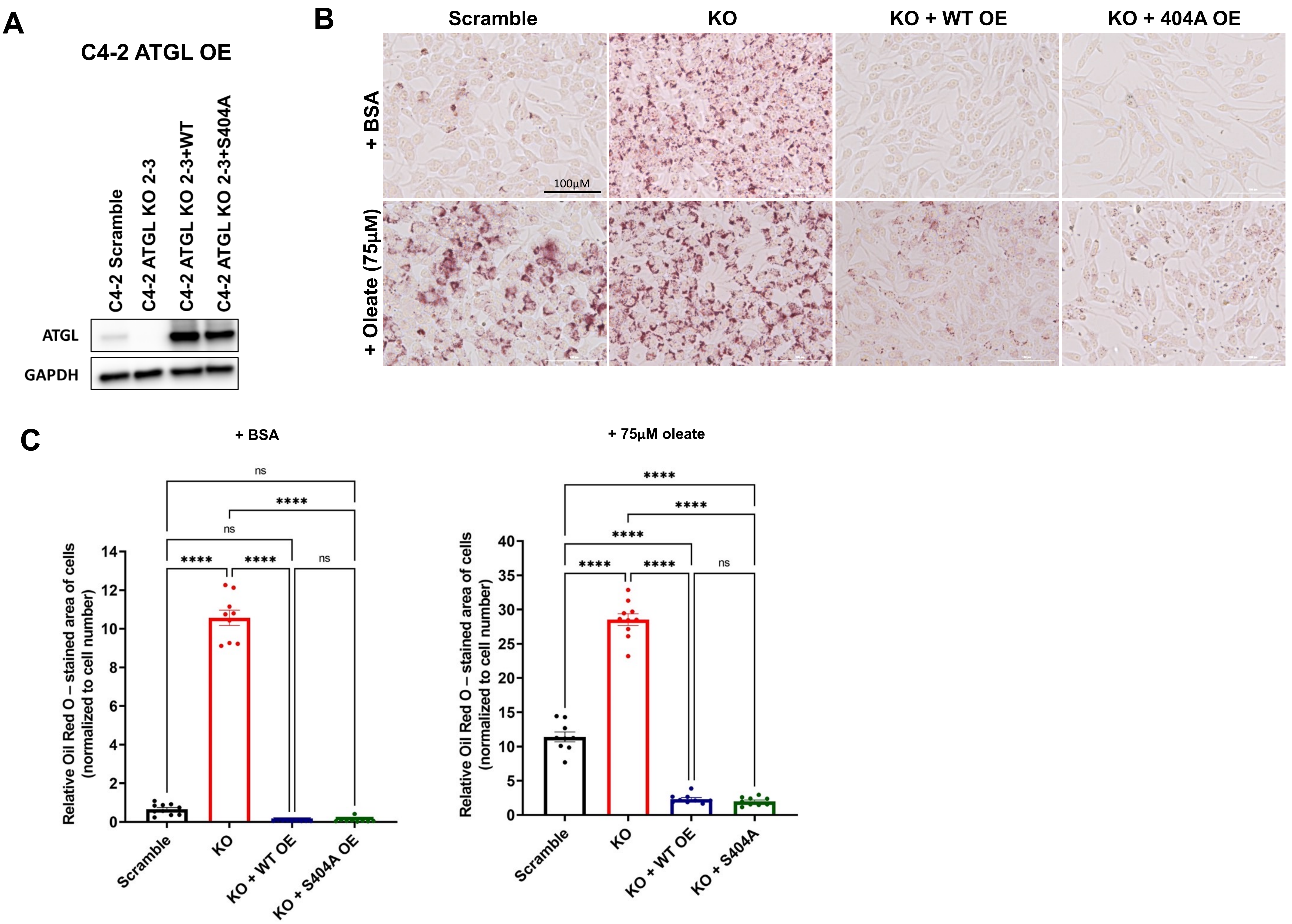


**Supplemental Figure S4: Overexpression of ATGL WT or S404A mutant lower neutral lipids below baseline levels.** (A-C) Overexpression (OE) of ATGL WT or 404A decreases Oil Red O staining, suggesting that increased protein level can supersede the effects of phosphorylation. (A) Immunoblot validation of ATGL KO and ATGL OE C4-2 models. (B) Representative images and (C) quantification of Oil Red O staining. Images and graphs are representative results of at least three independent experiments. One-way ANOVA. **P* < 0.05, ***P* < 0.01, ****P* < 0.001, *****P* < 0.0001, ns = no significance.

Supplemental Figure S5

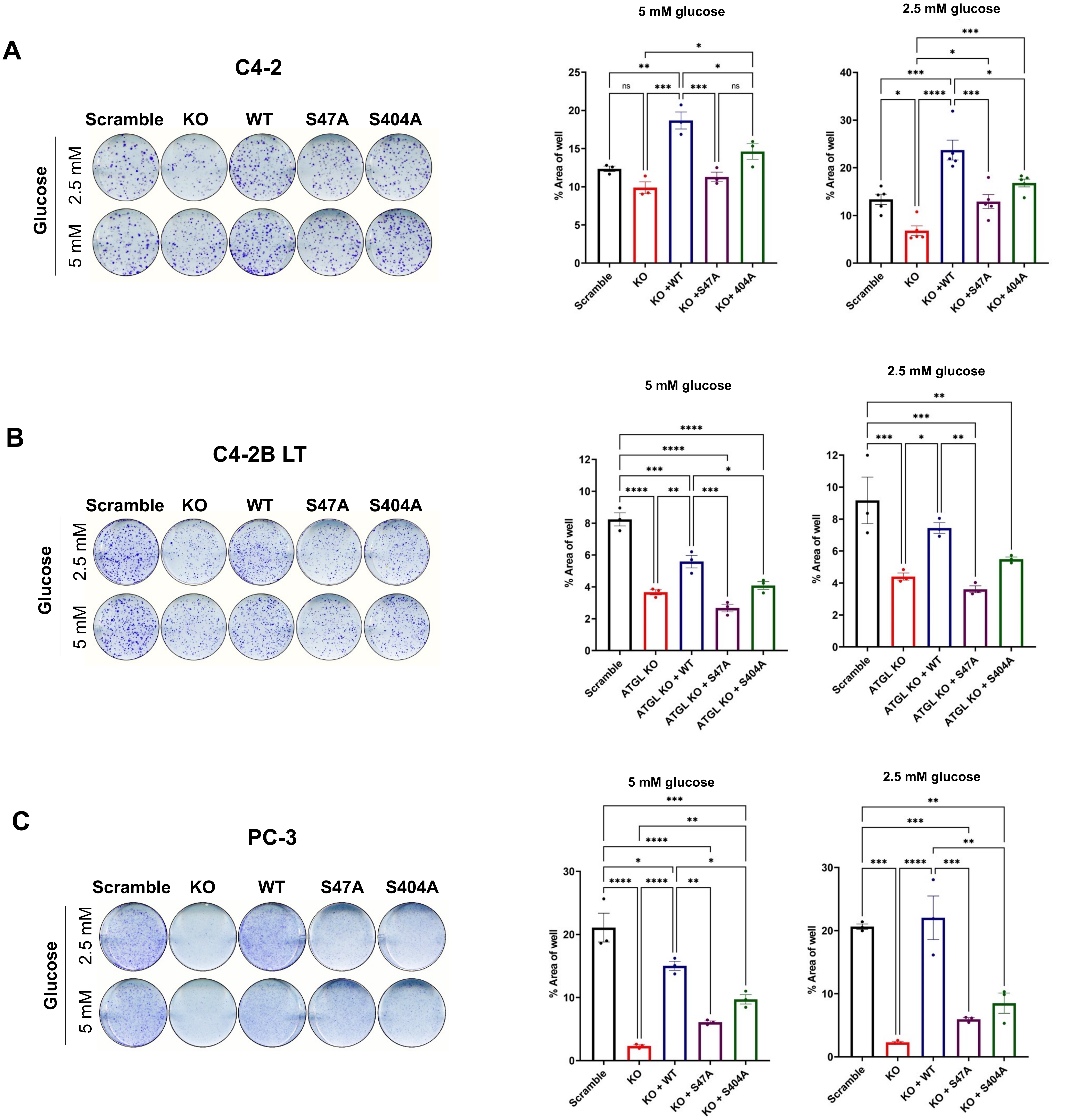


**Supplemental Figure S5: ATGL is required for CRPC cell model colony formation.** (A-C) Supplementing information for Figure 3C. Colony formation of Scramble, *PNPLA2* knockout (ATGL KO), and ATGL KO addback cell lines. (A) C4-2 and (B) C4-2BLT were grown in 5% dialyzed FBS supplemented with 2.5 mM or 5 mM glucose, (C) PC-3 were grown in 0.5% dialyzed FBS supplemented with 2.5 mM or 5 mM glucose. Images (*left*) and quantifications (*right*) are representative results of at least three independent experiments. One-way ANOVA. **P* < 0.05, ***P* < 0.01, ****P* < 0.001, *****P* < 0.0001, ns = no significance.

Supplemental Figure S6

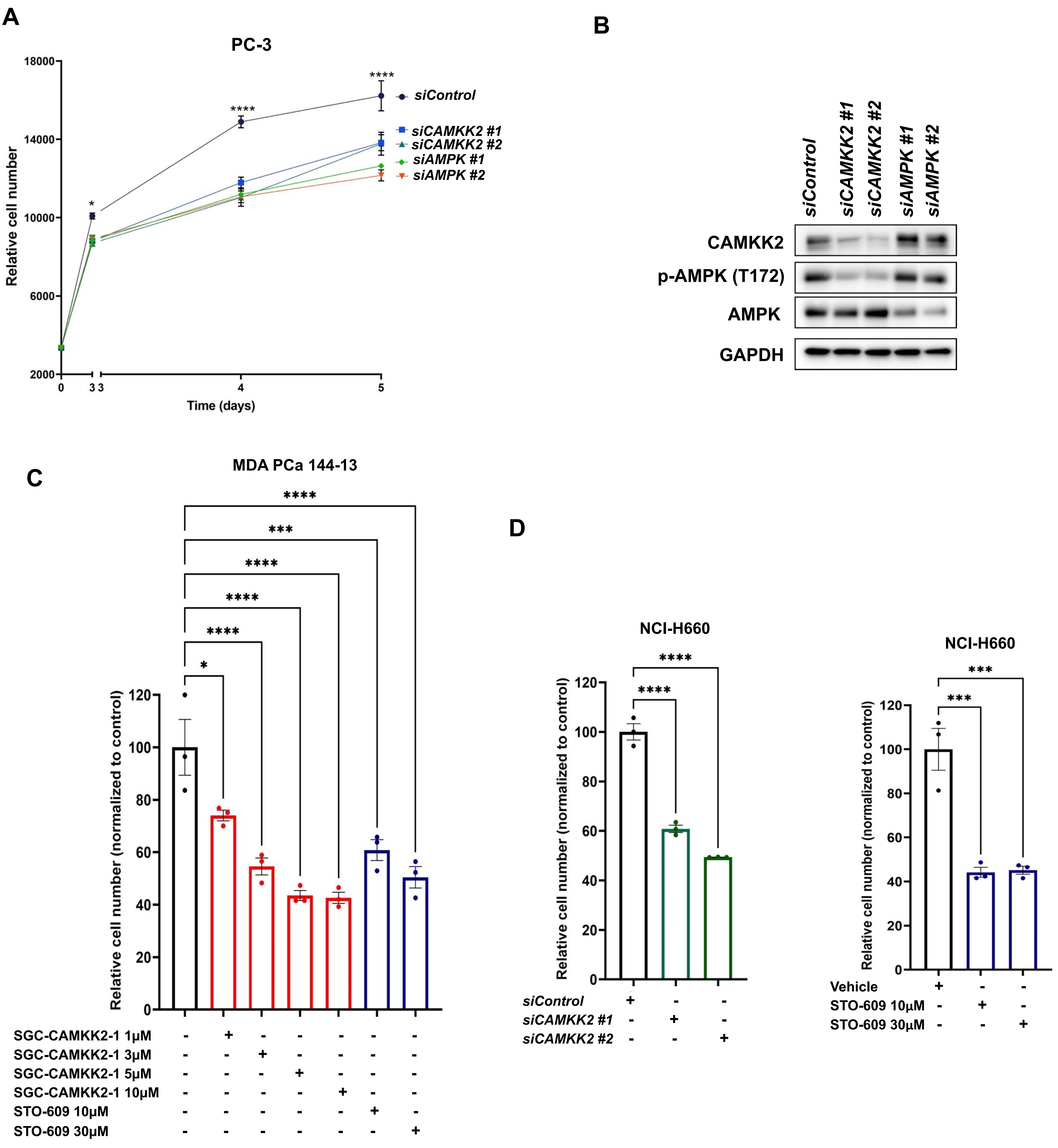


**Supplemental Figure S6: CAMKK2-AMPK signaling promotes AR- prostate cancer growth cell growth.** (A) Proliferation assay (Hoechst stain) of AR- PC-3 cells treated with chemical siRNA targeting *CAMKK2* or AMPK's α catalytic subunits *(PRKAA1/2)*. (B) Immunoblot control of CAMKK2 and AMPK knockdown in PC-3 cells. (C) Cell viability assay (resazurin) of NEPC MDA-PCa-144-13 cells treated with the CAMKK2 inhibitors SGC-CAMKK2-1 and STO-609. (D) Cell viability assay (resazurin) of NEPC NCI-H660 cells treated with siRNAs targeting *CAMKK2* or STO-609. Data are representative results of at least three independent experiments. One-way ANOVA. *$P$ < 0.05, **$P$ < 0.01, ***$P$ < 0.001, ****$P$< 0.0001, ns = no significance.

Supplemental Figure S7

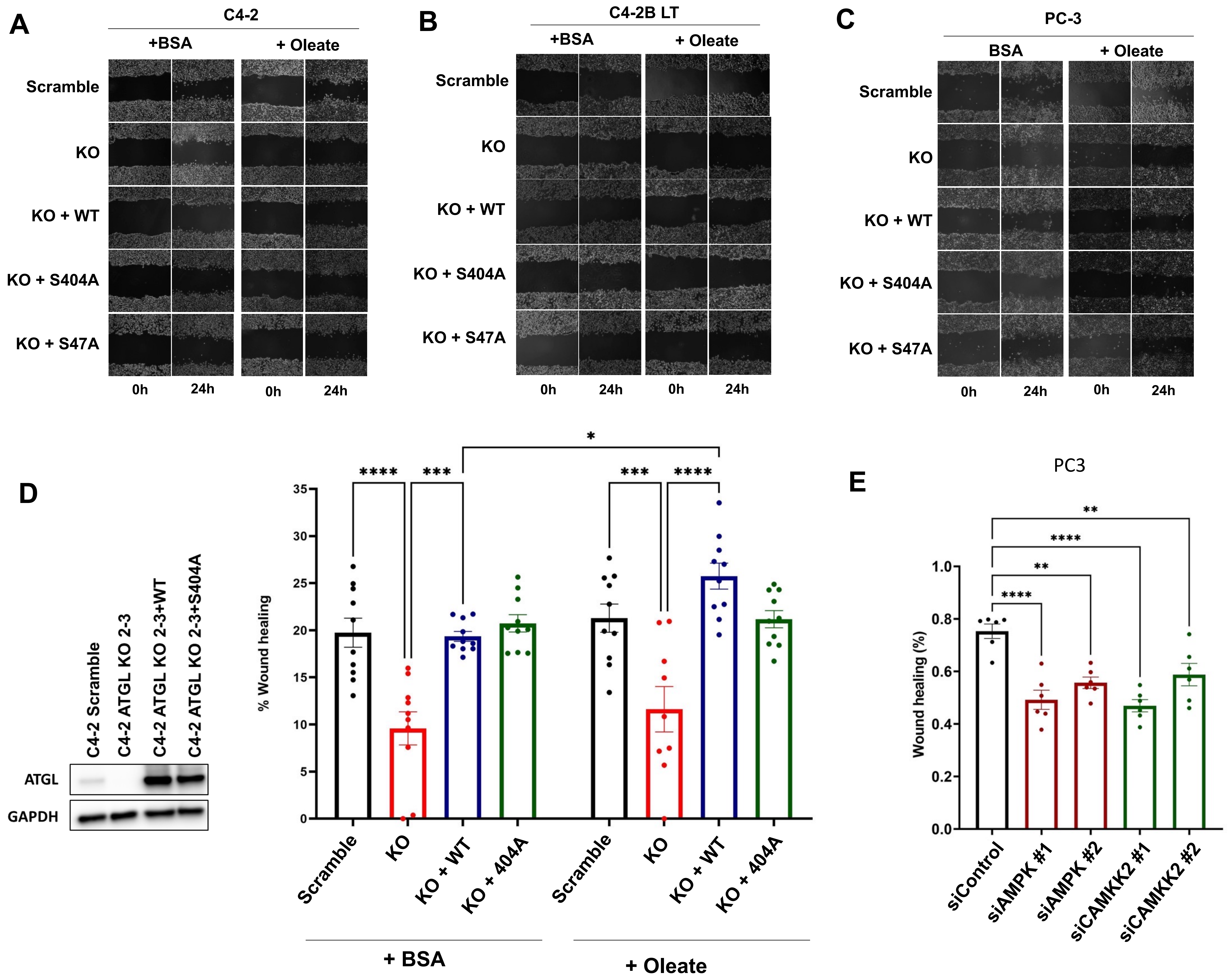


**Supplemental Figure S7: ATGL mediates prostate cancer cell migration.** (A-C) Representative images for wound healing/scratch assays from Figures 4A-C. (D) Migration (scratch) assay of C4-2 ATGL KO and addback cell lines with high expression of ATGL wildtype or ATGL S404A mutant. (E) Migration (scratch) assay of AR- PC-3 cells over 24h treated with chemical siRNA targeting *CAMKK2* or combines *PRKAA1+2* for 72h. One-way ANOVA. **P* < 0.05, ***P* < 0.01, ****P* < 0.001, *****P* < 0.0001, ns = no significance.

Supplemental Figure S8

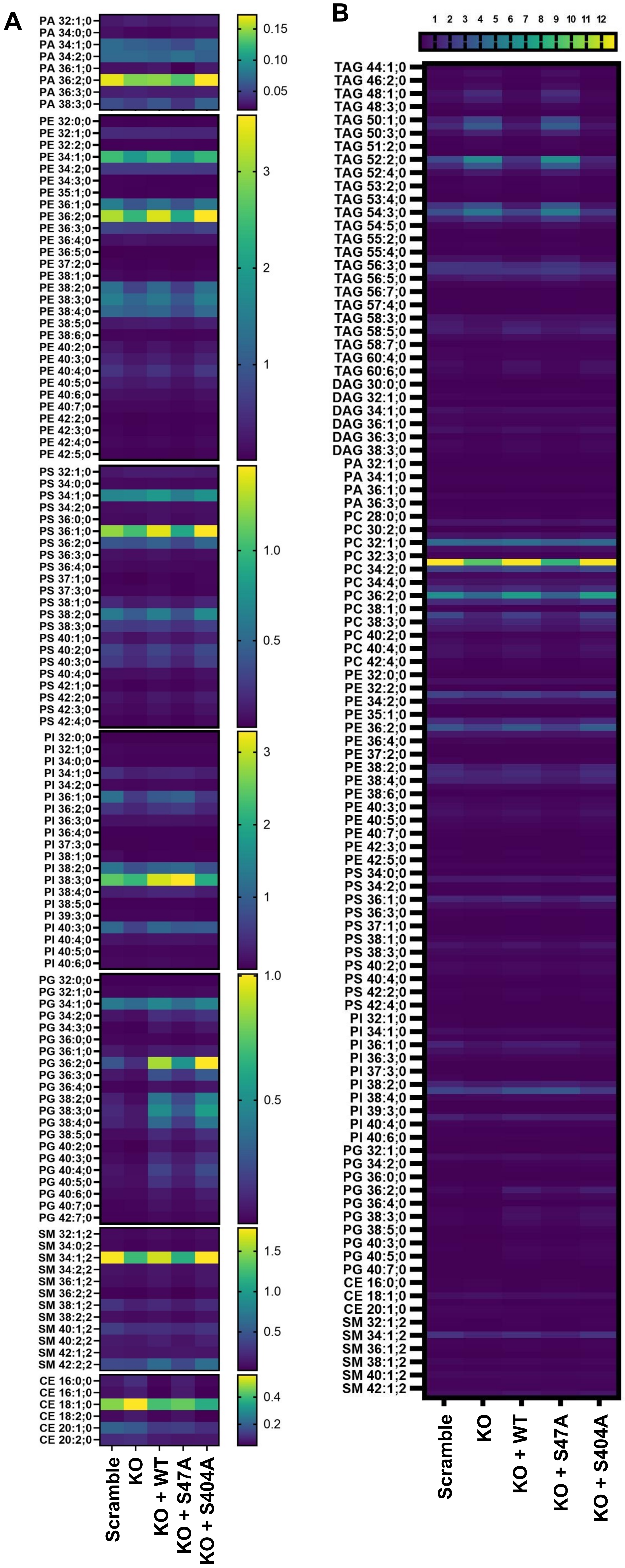


**Supplemental Figure S8: Heatmaps of shotgun lipidomics for CRPC ATGL KO and addback cell models.** C4-2 cells with Scramble sgRNA, *PNPLA2* knockout (ATGL KO) or ATGL KO with addback of ATGL wildtype (WT) or mutants (S47A, S404A) were analyzed via shotgun lipidomics. Cells were supplemented with 75μM BSA-coupled oleate overnight (A) Heatmaps of each lipid class to supplement Figure 5. Phosphatidic acid (PA), Phosphatidylethanolamine (PE), Phosphatidylinositol (PI), Phosphatidylglycerol (PG), Phosphatidylserine (PS), Sphingomyelin (SM), Cholesteryl ester (CE). (B) Unnormalized combined heatmap to show overall abundance of lipids detected in shotgun lipidomics. Sample size n = 3. One-way ANOVA. *$P < 0.05$, **$P < 0.01$, ***$P < 0.001$, ****$P < 0.0001$, ns = no significance.

### Supplemental Figure S9

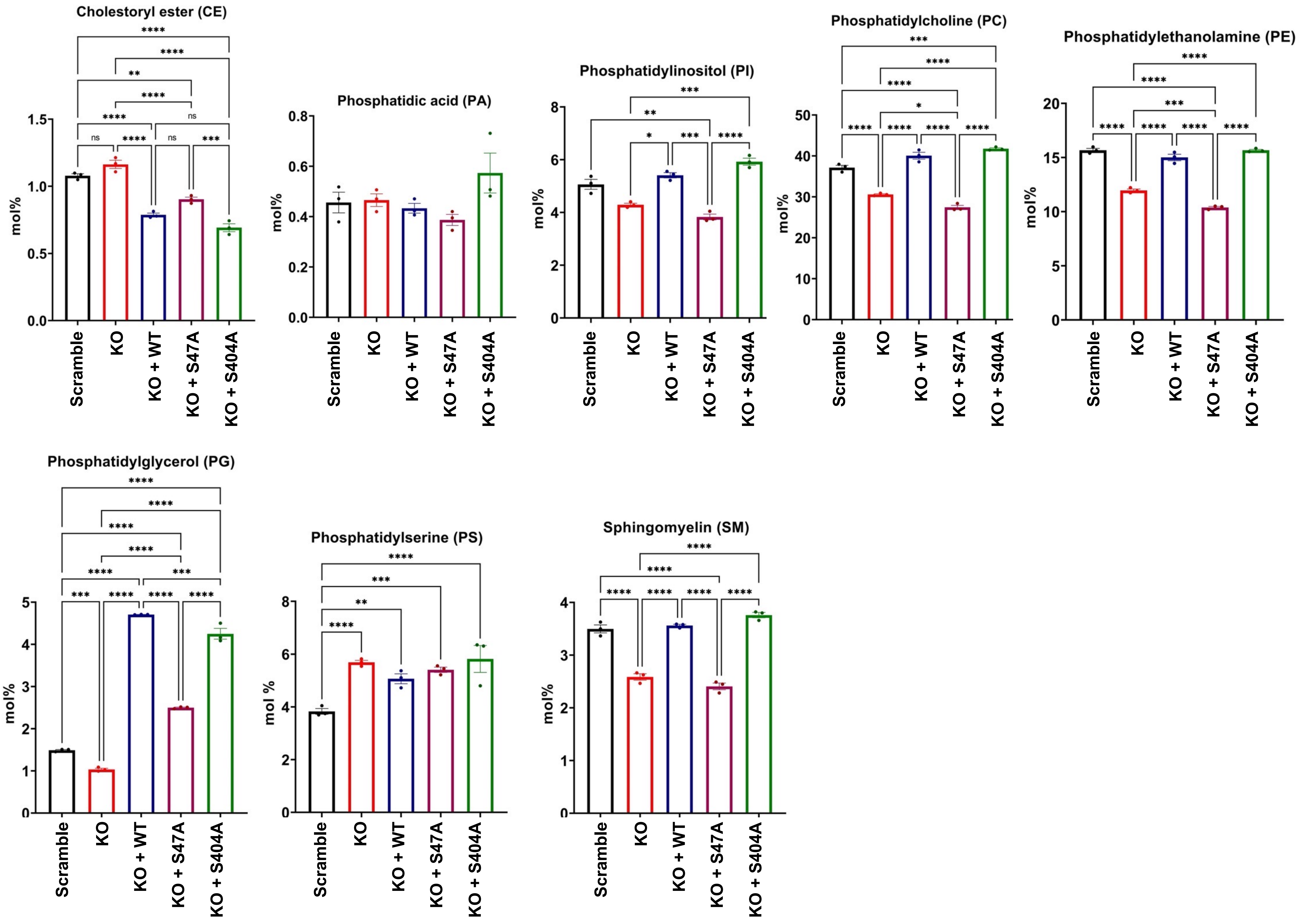


**Supplemental Figure S9: Shotgun lipidomics reveals significant changes in all detected glycerophospholipids upon ATGL knockout.** Quantification of the sum of individual lipid classes shown in Supplemental Figure S8 as heatmaps. Sample size n = 3. One-way ANOVA. **P* < 0.05, ***P* < 0.01, ****P* < 0.001, *****P* < 0.0001, ns = no significance.

Supplemental Figure S10

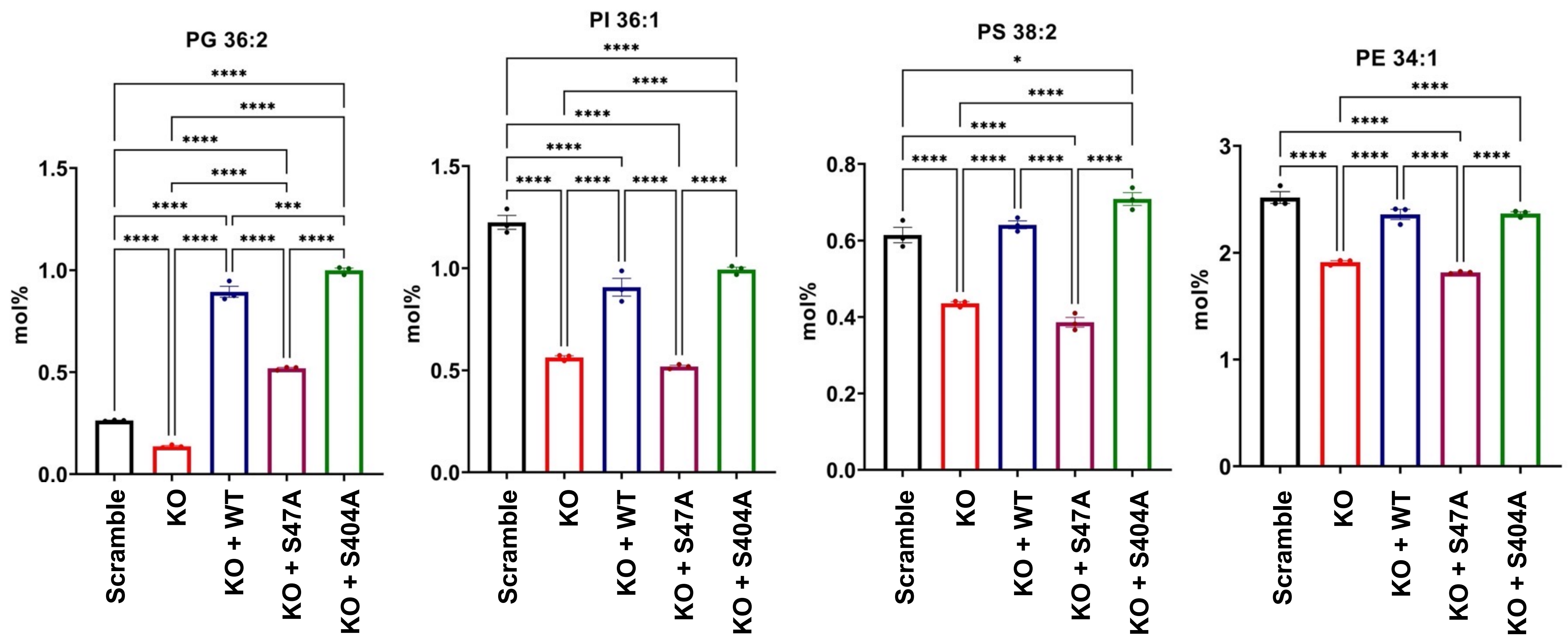


**Supplemental Figure S10: Additional lipid classes modulated by ATGL that are associated with prostate cancer.** C4-2 ATGL KO and addback cell lines were analyzed using shotgun lipidomics. Shown are additional lipid subclasses detected using shotgun lipidomics that are associated with prostate cancer (Butler et al. 2021). Data supplement Figure 4E. Phosphatidylethanolamine (PE), Phosphatidylinositol (PI), Phosphatidylglycerol (PG), Phosphatidylserine (PS). Sample size n = 3. One-way ANOVA. **P* < 0.05, ***P* < 0.01, ****P* < 0.001, *****P* < 0.0001, ns = no significance.

Supplemental Figure S11

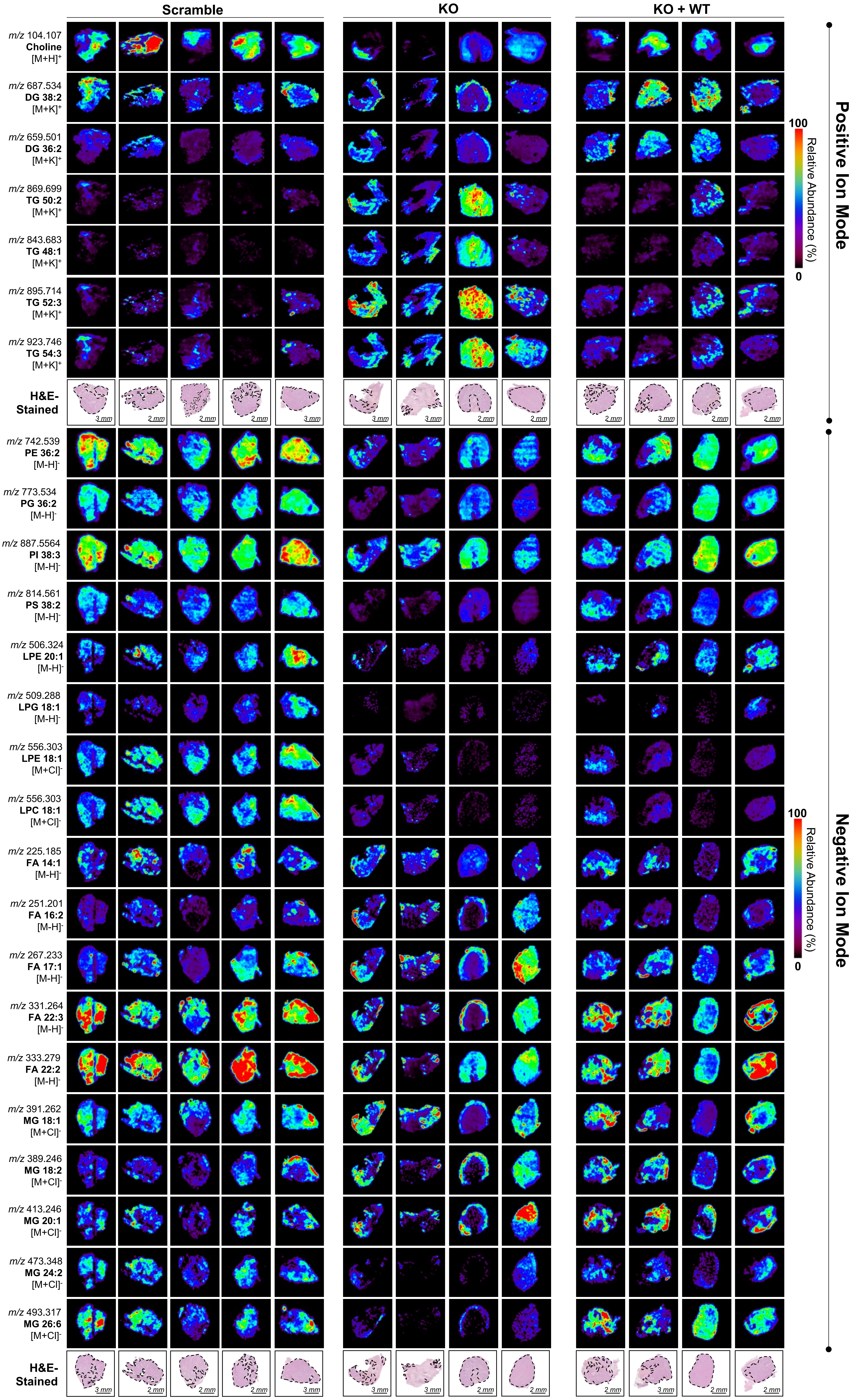


**Supplemental Figure S11: Additional DESI-MS images obtained from human CRPC xenografts.** Representative DESI-MS images of human CRPC xenografts described in Figure 3. Scramble control (n = 5), ATGL KO (n = 4), ATGL KO + WT (n=4).

Supplemental Figure S12

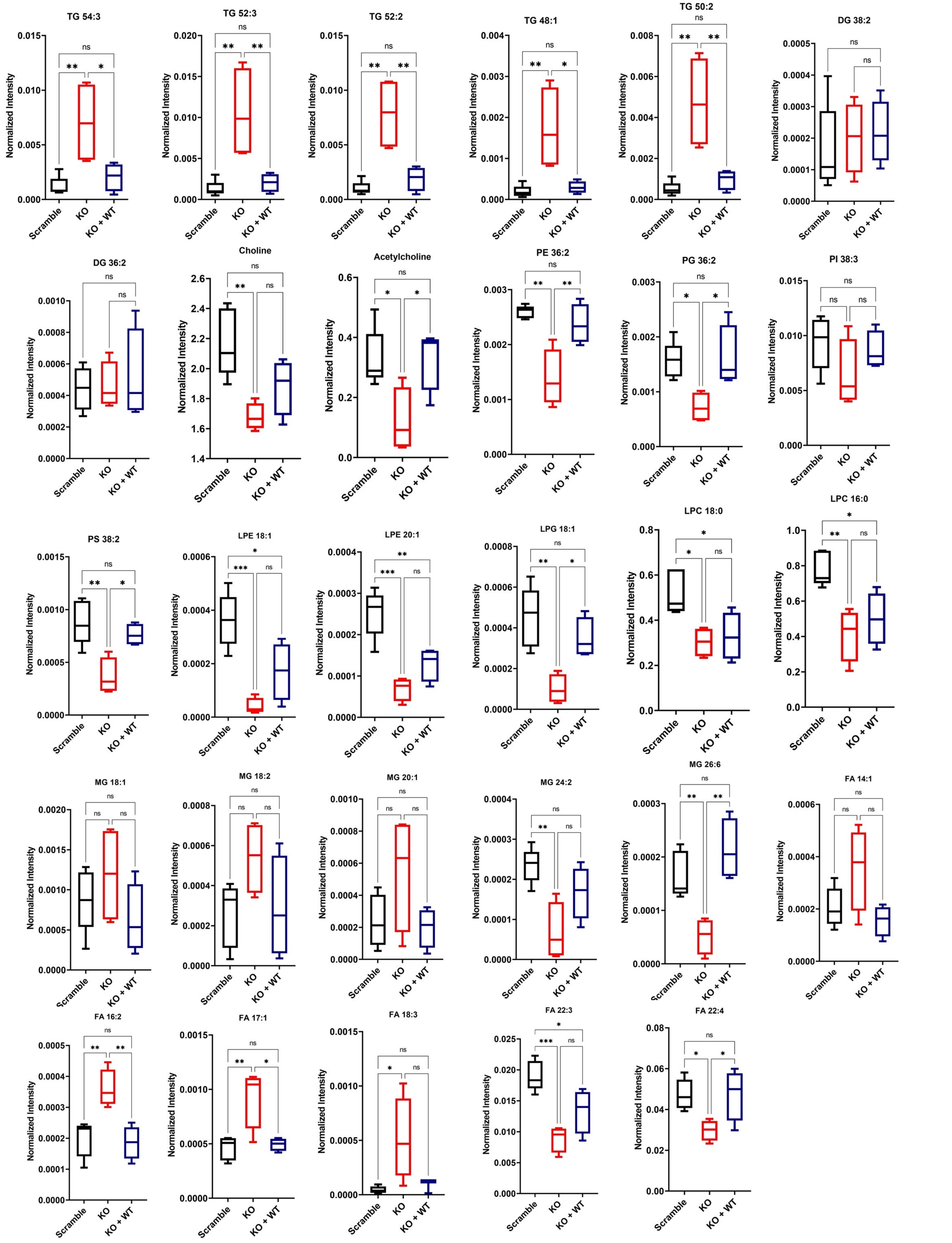


**Supplemental Figure S12: Ion intensity values extracted from tumor regions of ATGL KO and ATGL WT rescue CRPC xenografts.** Normalized intensities for metabolite and lipid species shown in Supplemental Figure S11 that were detected from human CRPC xenografts. Scramble control (n = 5), ATGL KO (n = 4), ATGL KO + WT (n=4). One-way ANOVA + Tukey. **P* < 0.05, ***P* < 0.01, ****P* < 0.001, *****P* < 0.0001, ns = no significance.

Supplemental Figure S13

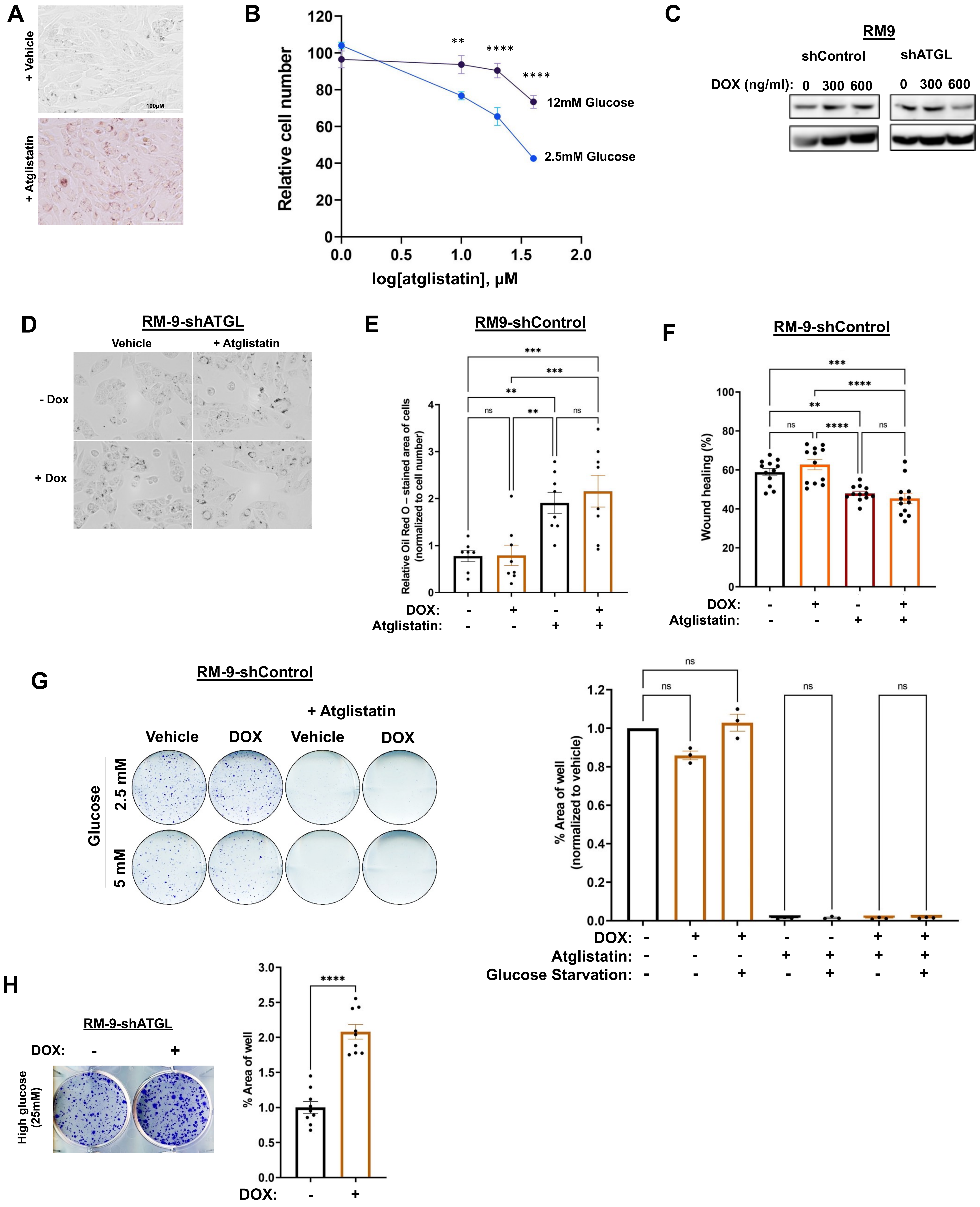


**Supplemental Figure S13: Knockdown of *PNPLA2* (encoding ATGL) or targeting ATGL pharmacologically leads to an increase in triglycerides and decrease in colony formation in murine prostate RM-9 cells.** These representative images and graphs are supporting information for Figure 7. (A) Brightfield images of Oil Red O-stained RM-9 cells treated with atglistatin from graph shown in Figure 7A. (B) 7-day cell growth curve of RM-9 cells treated with atglistatin in the presence of high or low glucose conditions (C) Immunoblot conformation of RM-9 stable cells with doxycycline (DOX)-inducible shRNAs targeting scramble control (shControl) or *Pnpla2* (shATGL). (D) Representative images of Oil Red O stains used for quantification shown Figure 7E. RM-9-shControl cells were used to test the effect on cells when treated with 600 ng/ml DOX: (E) Oil Red O stain, (F) migration (scratch assay) and (G) colony formation assay from cells treated with DOX and/or 10 µM atglistatin with (2.5 mM) or without (5 mM) glucose starvation. (H) Cells grown in the presence of high glucose media (DMEM high glucose, containing 25 mM glucose) exhibit increased colony formation following ATGL knockdown. One-way ANOVA. *$P < 0.05$, **$P < 0.01$, ***$P < 0.001$, ****$P < 0.0001$, ns = no significance.

Supplemental Figure S14

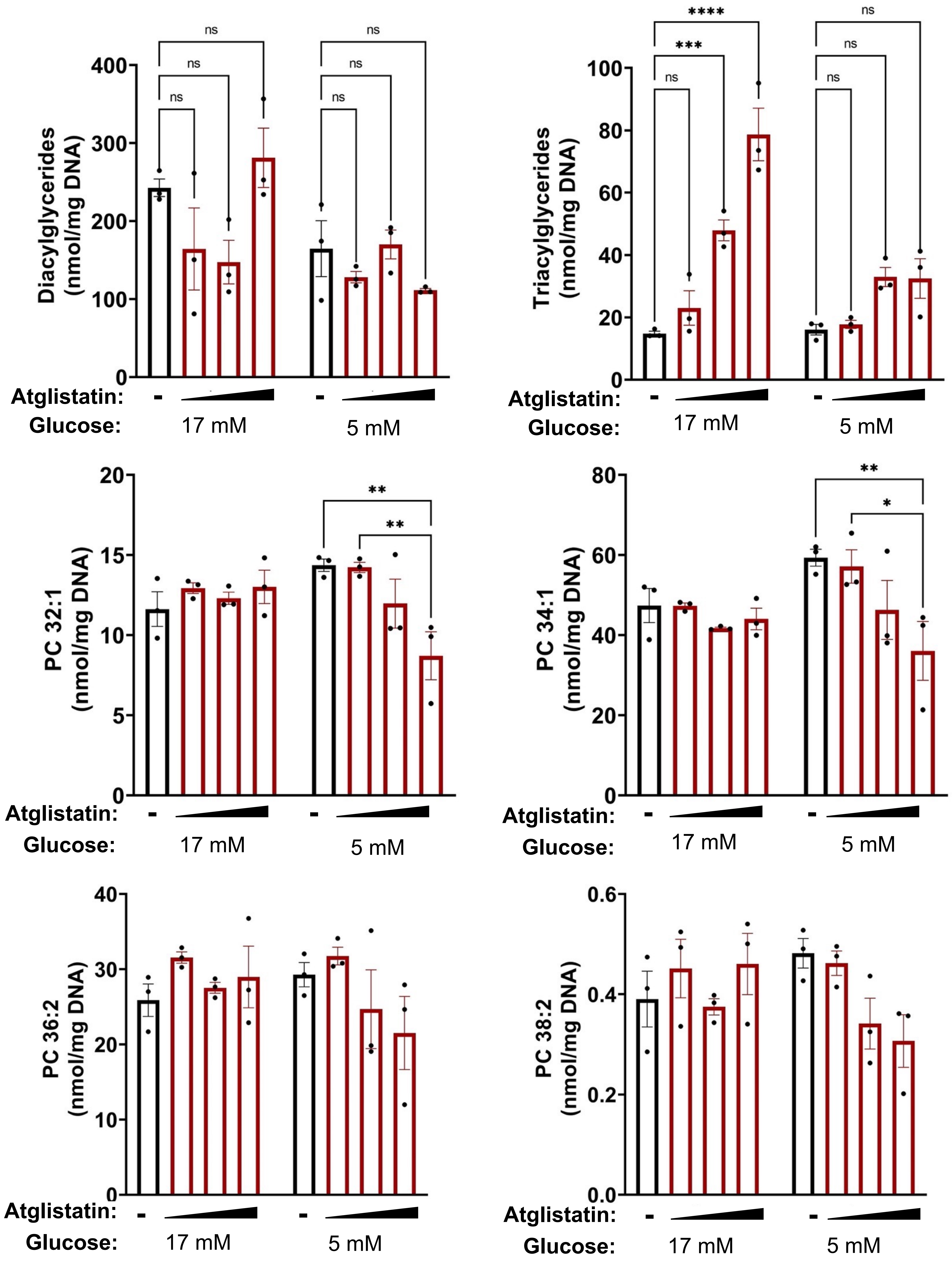


**Supplemental Figure S14: Lipidomics of organoids.** Quantification of lipidomics analysis of Hi-MYC organoids grown in 17 mM (standard organoid media) or 5 mM glucose (low glucose). Shown are triglycerides (TG), diglycerides (DG) and prostate-cancer associated phosphatidylcholines (PC) (Butler et al. 2021). Two-way ANOVA. **P* < 0.05, ***P* < 0.01, ****P* < 0.001, *****P* < 0.0001, ns = no significance.

Supplemental Figure S15

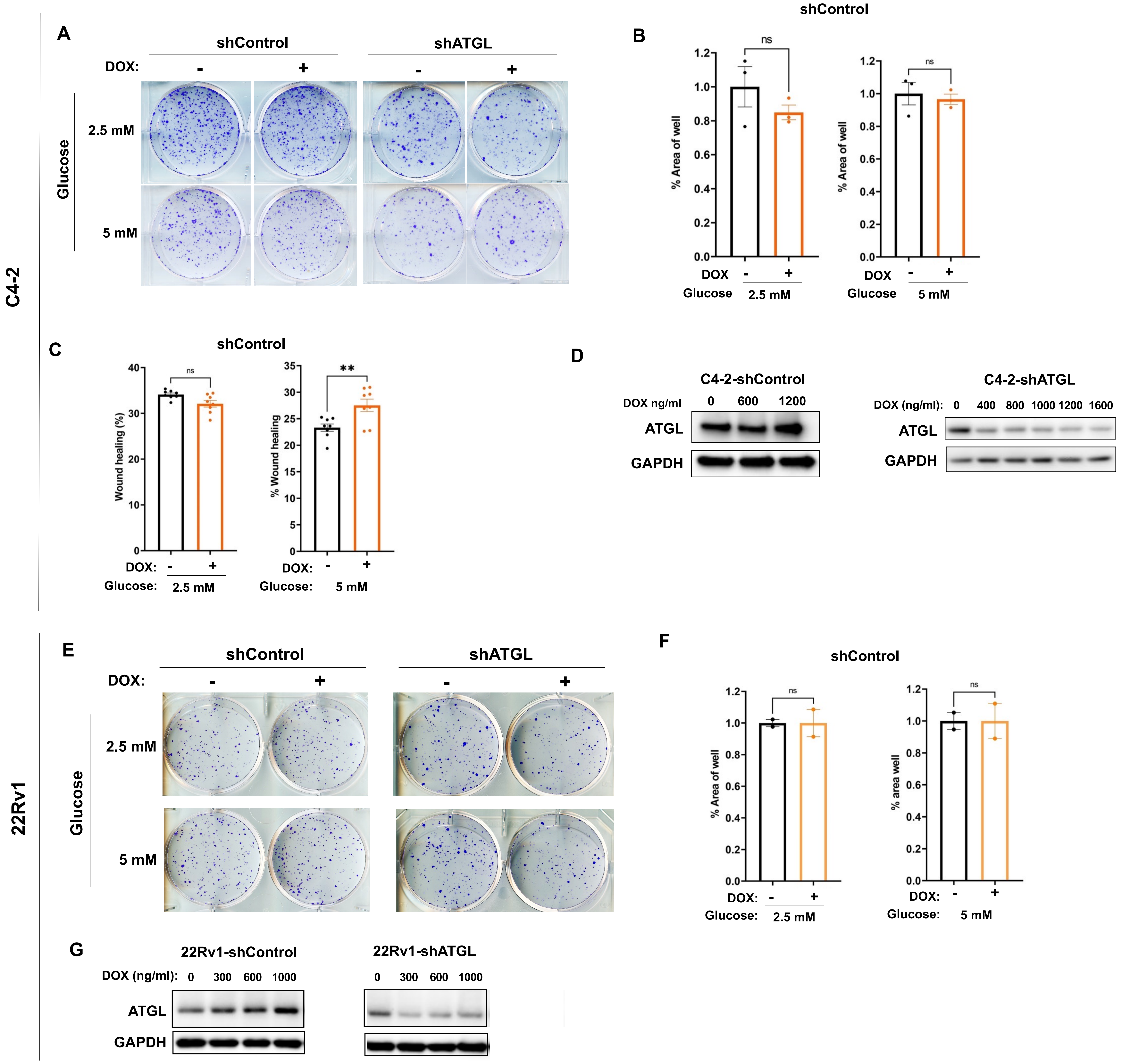


**Supplemental Figure S15: Molecular knockdown of *PNPLA2* (encoding ATGL) decreases colony formation in human CRPC cell models.** These representative images and graphs are supporting information for Figure 6. (A) Representative images of the colony formation assay of C4-2 shATGL cells shown in Figure 6N. Cells were treated with 600 ng/ml doxycycline (DOX). (B) Quantification of colony formation of C4-2 shControl shown in (A). (C) Quantification of migration of C4-2 shControl cells treated with or without 600 ng/ml DOX. (D) Immunoblot validation of ATGL knockdown via shRNA targeting *PNPLA2* but not scramble control in C4-2 stable cells. (E) Representative images of the colony formation assay of 22Rv1-shATGL cells shown in Figure 6O and (F) quantification of the colony formation of 22Rv1-shControl cells (control for DOX). (G) Immunoblot validation of ATGL knockdown via shRNA targeting *PNPLA2* but not scramble control in 22Rv1 stable cells. One-way ANOVA. **P* < 0.05, ***P* < 0.01, ****P* < 0.001, *****P* < 0.0001, ns = no significance.

Supplemental Figure S16

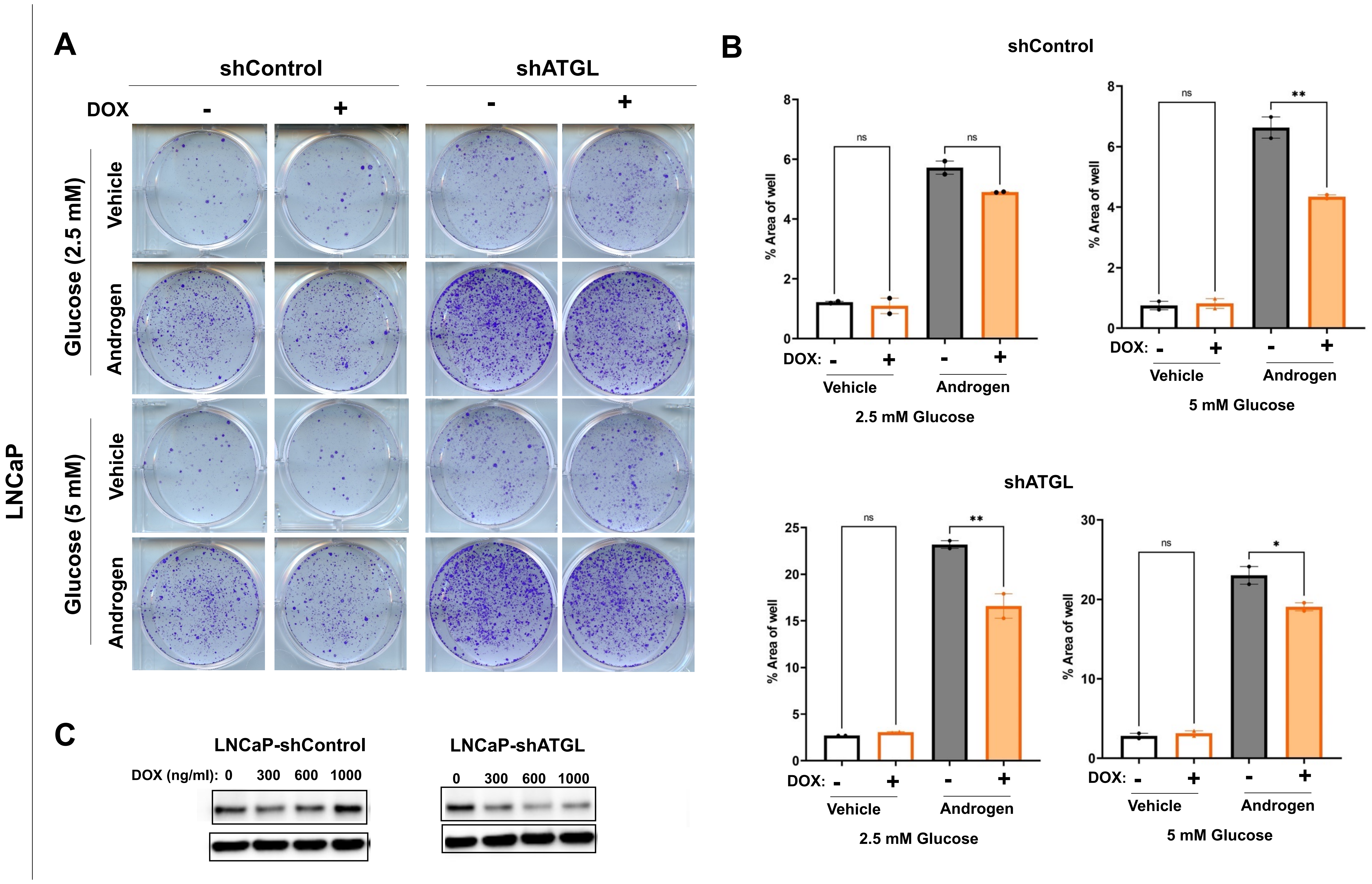


**Supplemental Figure S16: ATGL does not contribute to hormone-sensitive prostate cancer cell growth.** (A-C) LNCaP cells stably expressing inducible shRNAs targeting scramble control (shControl) or *PNPLA2* (shATGL) were treated with or without 600 ng/ml doxycycline (DOX) with vehicle or 1 nM of the synthetic androgen R1881. Cells were subjected to (A) colony formation, (B) wound healing, and (C) immunoblot control assays as indicated. One-way ANOVA. *$P$ < 0.05, **$P$ < 0.01, ***$P$ < 0.001, ****$P$ < 0.0001, ns = no significance.
